# Proteo-transcriptomic reprogramming and resource reallocation define the aging mammalian brain

**DOI:** 10.1101/2025.08.14.669896

**Authors:** Nisha Hemandhar-Kumar, Verena Kluever, Svenja V. Kaufmann, Cornelius Bergmann, Kanaan Mousaei, Miguel Tomas, Miguel Correa Marrero, Avika Chopra, Misa Hirose, Mercè Pallas, Coral Sanfeliu, Saleh M. Ibrahim, Andre Fischer, Tiago F. Outeiro, Henning Urlaub, Tatjana Tchumatchenko, Carlos López Otín, Eugenio F. Fornasiero

**Affiliations:** Department of Neuro- and Sensory Physiology, University Medical Center Göttingen, 37073 Göttingen, Germany; Cologne Excellence Cluster for Aging and Aging-Associated Diseases (CECAD), University of Köln, Köln, Germany; Max Planck Institute for Multidisciplinary Science, Bioanalytical Mass Spectrometry, Am Fassberg 11, 37077 Göttingen, Germany; University Medical Center Göttingen, Institute of Clinical Chemistry, Bioanalytics, Robert Koch Straße 40, 37075 Göttingen, Germany; Institute of Experimental Epileptology and Cognition Research, University of Bonn Medical Center, Venusberg-Campus 1, 53127 Bonn, Germany; Institute of Molecular Systems Biology, Department of Biology, ETH Zurich, Otto-Stern-Weg 3, Zurich, 8093, Zurich, Switzerland; Swiss Institute of Bioinformatics, 1015 Lausanne, Switzerland; University Medical Center Göttingen, Department of Experimental Neurodegeneration, Center for Biostructural Imaging of Neurodegeneration, Göttingen, Germany; Lübeck Institute of Experimental Dermatology, University of Lübeck; Institute of Neurobiology, University of Lübeck; Departemt Pharmacology and Therapeutic Chemistry, Institut de Neurociències-Universitat de Barcelona, Barcelona, Spain; CIBERNED, Instituto de Salud Carlos III, Madrid, Spain; Department Neuroscience and Experimental Therapeutics, Institute of Biomedical Research of Barcelona (IIBB), Consejo Superior de Investigaciones Científicas, Barcelona, Spain; Center for Biotechnology, Department of Medical Sciences, Khalifa University, Abu Dhabi, United Arab Emirates; Cluster of Excellence “Multiscale Bioimaging: from Molecular Machines to Networks of Excitable Cells” (MBExC), University of Göttingen, Göttingen, Germany; German Center for Neurodegenerative Diseases, Göttingen, Germany; Translational and Clinical Research Institute, Newcastle University, UK; Departamento de Bioquímica y Biología Molecular, Instituto Universitario de Oncología, Universidad de Oviedo, Spain; Facultad de Ciencias de la Vida y la Naturaleza, Universidad Nebrija, Madrid, Spain; Centre de Recherche des Cordeliers, Université de Paris Cité, Sorbonne Université, INSERM U1138, France; Department of Life Sciences, University of Trieste, 34127 Trieste, Italy

**Keywords:** Proteo-transcriptome analysis, multiomics data integration, nuclear retention, protein aggregation, proteostasis, neurodegeneration

## Abstract

Brain aging is a major risk for neurodegeneration, yet the underlying molecular mechanisms remain poorly understood. Here we performed an integrative proteo-transcriptomic analysis of the aging mouse brain, uncovering molecular signatures of aging through the assessment of protein aggregation, mRNA relocalization, and comparative proteomics across eight models of premature aging and neurodegeneration. We identified dynamic changes in physiological aging highlighting differences in synaptic maintenance and energy-allocation. These were linked to changes associated with fundamental protein biochemical properties such as size and net charge. Network analysis highlighted a decrease in mitochondrial complex I proteins not compensated at the mRNA level. Aggregation of 60S ribosome subunits indicated deteriorating translation efficiency and was accompanied by mitochondrial and proteasomal imbalance. The analysis of the nine models revealed key similarities and differences between physiological aging and pathology. Overall, our study provides an extensive resource on molecular aging, and offers insights into mechanisms predisposing to neurodegeneration, easily accessible at our Brain Aging and Molecular Atlas Project (BrainAging-MAP) website.

## Introduction

Brain aging affects the regulation of DNA transcription, mRNA translation, and protein homeostasis (proteostasis) and is linked to an increased risk for age-related neurodegenerative diseases (NDDs) such as Parkinson’s and Alzheimer’s disease^1–5^. Omics approaches have been employed to simultaneously measure mRNA and protein level changes during brain aging^5–10^ showing that proteostasis regulation is essential for healthy aging^4,11–18^ and its decline can lead to incorrect protein folding and accumulation of toxic unfolded proteins within cells^15,19–22^. Protein accumulation can affect the composition and localization of protein complexes^23^, resulting in overall changes in protein abundance, decoupling of mRNA and protein level and contributing to aging phenotypes. Indeed, evidence from non-mammalian models suggest an age-dependent loss of correlation between mRNA and protein levels along with changes in translation and protein degradation^11,12^. However, a deep concomitant characterization of mammalian transcriptomes and proteomes during brain aging considering the effects of protein aggregation and mRNA localization is currently lacking.

Despite extensive research on age-related changes in protein and transcript levels^5,11,14,24,25^, our understanding of the regulatory mechanisms governing the different molecular layers, as well as the relationships between layers remains limited^9,13,24,26–28^. Key questions remain as to whether systematic proteo-transcriptomic features, such as mRNA sequence, amino acid composition, and biochemical properties shape interactions between omics layers. In this context, evidence from our lab suggests that amino acid cost, a fundamental biochemical constraint, influences protein production and stability in the aging brain^5^. Given the interdependence between mRNA composition, protein levels, stability, and cellular phenotypes^29–32^, a detailed proteo-transcriptome analysis could elucidate their collective role in brain aging and potentially facilitate the development of therapeutic targets for age-related brain disorders.

In this study, we performed a comprehensive analysis in young (6 months - 6m), middle (12m) and aged (24m) mouse brains including several omics layers such as mRNA and protein abundances, nuclear mRNAs abundance and levels of aggregated proteins. We also studied the changes of the total and the aggregated proteome for 8 mouse models and their respective controls recapitulating accelerated aging (aAGE) and NDDs to understand similarities and differences between these perturbations and physiological aging.

Our multifaceted analysis, whose data serves as a comprehensive, updatable and easy to access resource for the community (BrainAging-MAP) revealed several important new aspects of brain aging. For example, we observed that in mammals, proteo-transcriptomic changes for several pathways follow dynamic, non-linear trajectories. We also revealed a possible energy-reallocation switch where synapse maintenance might get compromised to ensure survival-oriented energy conservation. Importantly, transcriptional alterations do not always translate into protein level changes, as in the case of nuclear-encoded mitochondrial Complex I genes, where mRNA level increase is not reflected at the protein level, leading to late-aging translational inefficiency and providing new mechanistic insights into why energy metabolism declines in the aging brain. The additional comparative analysis of aAGE and NDD models with physiological aging allowed us to identify common altered modules such as accumulation of cytoskeletal components in accelerated aging and aggregation of 60S ribosomes in the insoluble fraction of NDD models, while differences point to possible adaptations that might reflect resilience mechanisms.

Collectively, our findings challenge the assumption that transcriptional regulation can also be among drivers of aging-related dysfunction and underscore the importance of proteo-transcriptome analyses as a major aspect for understanding changes occurring during brain aging.

## Results

### Gene expression changes in the aging brain

To obtain a comprehensive analysis of the proteo-transcriptome changes observed during brain aging we initially considered physiologically aged mice and measured in parallel protein and mRNA levels at 6m (young adult), 12m (middle-aged) and 24m (aged; see methods for the age choice). We designed an integrated workflow that enabled the quantification of these aspects in parallel starting from the same tissue samples (**Fig. 1a**). Rather than microdissecting brain subregions, we chose to use whole-brain tissue using bulk measurements to capture global molecular trends. While this approach lacks cellular resolution, it ensures consistent sampling across age groups and facilitates integration of diverse molecular readouts from the same biological source. Furthermore, while single-cell/single-nuclei measurements are invaluable for understanding cellular trajectories in aging^33–37^, they are currently limited in capturing extranuclear components (such as synaptic mRNAs and proteins), key elements in age-related neuronal changes. Importantly, this strategy allowed us to reduce sample-to-sample variability and preserve the integrity of parallel measurements, which is critical for a quantitative systems-level integration across molecular layers.

**Figure 1:**
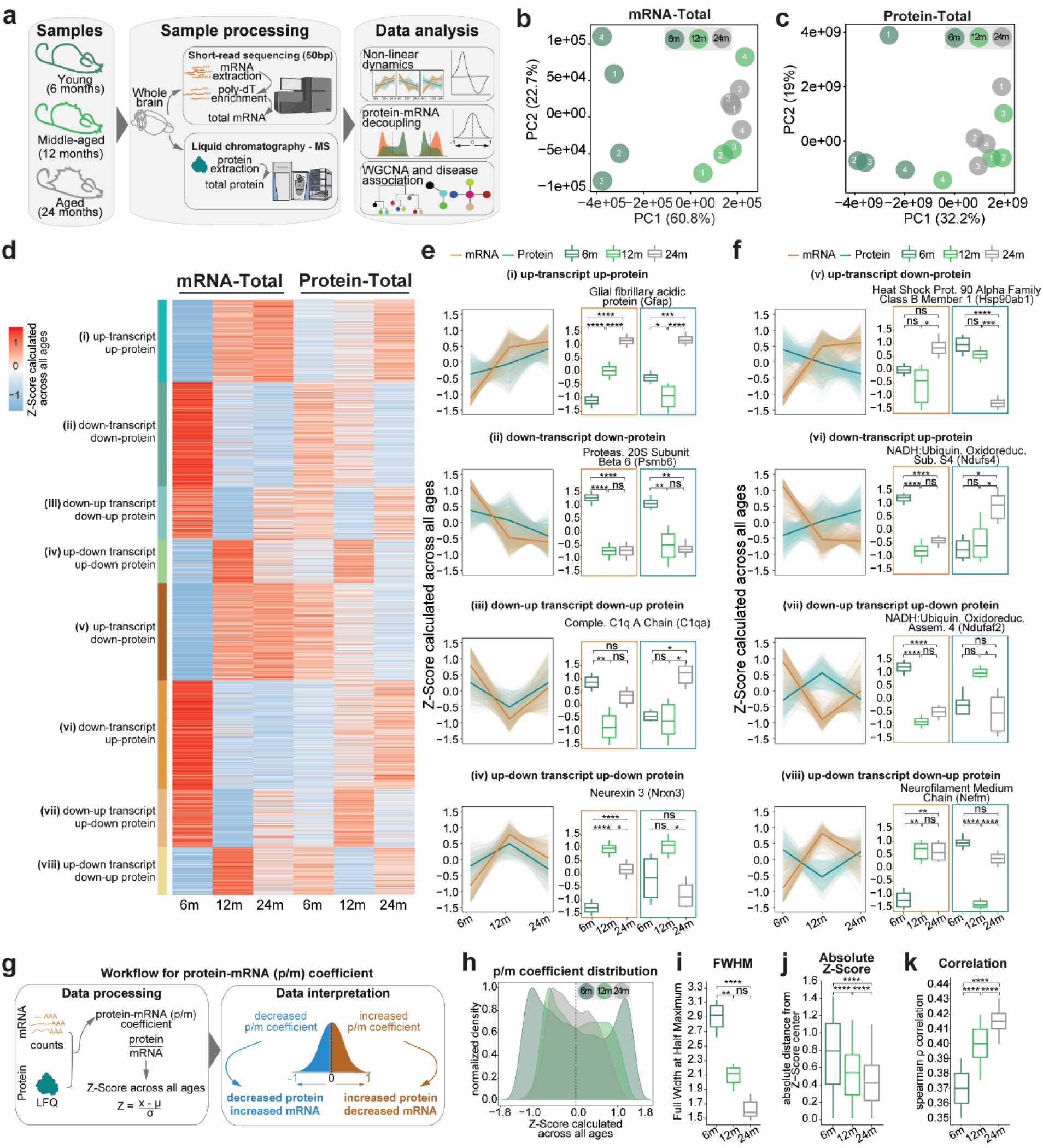
Non-linear dynamics and protein-mRNA coefficients in the aging mouse brain. (**a**) Schematic of the experimental and analytical workflow assessing mRNA and protein levels in the total fraction of aging mouse brains across animals aged 6-months (6m), 12-months (12m), and 24-months (24m). (**b, c**) Dot plots depict biological replicate variability via principal component analysis (PCA) for the (**b**) mRNA-total and (**c**) protein-total datasets. Dark green represents 6m, light green is 12m and gray is 24m (**d**) Heatmap illustrating eight distinct patterns in mRNA and protein levels across ages. Color gradient for the Z-scores across all ages (red increased and blue decreased). (**e, f**) Line plots of mRNA and protein trajectories showing (**e**) similar and (**f**) opposite trends across molecular layers. Brown represents mRNAs and teal proteins. (**g**) Workflow for analysis and categorization of protein-mRNA coefficient (p/m coefficient). Positive coefficient values (brown) indicate increased protein levels relative to mRNA, suggesting efficient translation; negative values (blue) indicate reduced protein levels relative to mRNA, indicating inefficient translation. (**h**) Density distribution of p/m coefficient for 6m, 12m and 24m. (**i**) Full width half maximum (FWHM) for the density distribution for 6m, 12m and 24m in Fig. 1h. (**j**) Absolute distance from Z-Score center for the density distribution for 6m, 12m and 24m in Fig. 1h. (**k**) Protein-mRNA spearman ρ correlation for each age. P-values indicate the results of paired t-test followed by Tukey post hoc test P-value * ≤ 0.05, ** ≤ 0.01, *** ≤ 0.001 and **** ≤ 0.0001.

We first assessed inter-sample variability and correlation among replicates using principal component analysis (PCA) and spearman’s ρ correlation (**Fig. 1b** and **c, Extended Data Fig. 1a and b**). As expected, biological replicates exhibit distinct grouping patterns based on their respective ages in the mRNA-total fraction (**Fig. 1b, Extended Data Fig. 1a**). Specifically, samples from 12m and 24m cluster together, while those at 6m form a separate cluster. In our proteomics dataset, the total fraction shows similar age-based grouping patterns (**Fig. 1c, Extended Data Fig. 1b**) and biological replicates within each age group are consistent. These findings show that the proteo-transcriptome landscape undergoes more substantial changes from young adulthood to middle age.

We then performed differential gene and protein expression analysis by comparing mRNA and protein abundance across ages obtained with short-read next-generation sequencing and Liquid Chromatography Mass Spectrometry (LC-MS/MS). To estimate the reliability of our protein quantifications, we also performed parallel reaction monitoring (PRM) and assessed levels for 8 proteins whose levels were significantly changed in our dataset (Ina, Lsamp, Nefh, Plec, Ermn, Vim, Gfap, C1qb) confirming the changes measured in our quantification method based on BoxCar^38^ (**Extended Data Fig. 1c**).

While extremely informative and detailed, due to space limitations these analyses are summarized in the supplementary section of the work (**Supplementary Text 1**; **Extended Data Figs 2-11** and **Supplementary Table 1**). In short, using mRNA and protein level comparisons between young and aged mouse brains, significant upward and downward trends were observed, especially within genes related to synaptic function and immune response, with a marked increase in astrocyte-specific markers such as Gfap and vimentin (Vim) at 24m. Also consistent with previous observations^5,24,39,40^, GO analyses suggest adaptive aging responses in the brain.

One of the most interesting novel observations was that several intermediate filament proteins, including neurofilament proteins (Nefl, Nefm, Nefh), Vim, plectin (Plec), and α-internexin (Ina), showed a down-up trend with significantly lower abundance at 12m than both 6m and 24m (**Extended Data** Fig. 10, 11a, c). This is a clear example of a non-linear process occurring in brain aging, highlighting the importance of considering middle-aged animals in longitudinal datasets.

### Non-linear dynamics and bidirectional trends in the aging brain highlight complexity in proteo-transcriptome changes

To explore the complex patterns of proteo-transcriptome dynamics during aging, we grouped genes into eight unique temporal trends based on both their mRNA and protein levels at different ages (Fig. 1d). We sub-selected ∼3500 genes based on defined filtering criteria to ensure reliability (see Methods for details). We thus identified four patterns that show similar trends between mRNAs and proteins (i-iv, Fig. 1e) and four that show opposite trends (v-viii, Fig. 1f).

When analyzing the patterns that showed similar trends in the protein and mRNA datasets, we found that among the consistently upregulated genes (i in Fig. 1d**,e**, 14%, N = 492) in the aged brain, several hits are linked to neuron projection development, polymeric cytoskeletal fibers (GO:0031175, GO:0099513**, Extended Data** Fig. 11d**, Supplementary Table 1**) and glial function. For example, Gfap showed continuous increase in both mRNA and protein datasets. Among the 18% of mRNA-protein couples that were continuously decreased during aging (ii; N = 624, Fig. 1d, e**, Supplementary Table 1**), we found hits corresponding to proteasome, ribosome (GO:0042254**, Extended Data** Fig. 11d**, Supplementary Table 1**), and mitochondrial function (GO:0033108**, Extended Data** Fig. 11d**, Supplementary Table 1**). One example is the proteasome 20S subunit beta 6 (Psmb6, Fig. 1e). Overall, these findings suggest mitochondrial dysfunction, loss of ribosome stoichiometry and impaired proteostasis, as observed in previous studies in the aging brain^5,11,12,14,39–41^.

We also found non-linear trends linked to aging that are characterized by bidirectional changes. For example, in one group relatively high levels were found at 6m, followed by a decrease at 12m, and an increase again at 24m (iii, down-up, 9%, N = 319). We observed another group showing the inverse pattern of up-down regulation (iv, 7%, N = 263; Fig. 1d, e**, Supplementary Table 1**). Some notable examples for these non-linear patterns include complement proteins such as C1qa (Fig. 1e), involved in immune function, which exhibited a down-up trend, and neurexin 3 (Nrxn3, Fig. 1e**, Supplementary Table 1**), a cell adhesion molecule enriched in neurites involved in synaptic and cognitive function which showed an up-down pattern. Collectively, these findings suggest that only for ∼50% of gene products there is a clear longitudinal coupling between mRNA and proteins.

Importantly, in our analysis we also found four patterns that show opposite trends between mRNA and protein levels (v-viii, Fig. 1d, Fig. 1f**, Supplementary Table 1**). Among these pairs we observed 16% of genes (N = 574) that during age increase in mRNA but are decreased in protein abundances (up-transcript, down-protein, v), while 18% (N = 648) showed the inverse pattern (down-transcript, up-protein, vi). We also observed two groups with non-linear bidirectional patterns with 10% of genes (N = 346) exhibiting a down-up pattern in mRNA with an up-down pattern in protein (vii), and 8% of genes (N = 289) with the opposite non-linear trend (up-down transcript and down-up in protein, viii).

Among the up-transcript and down-protein group (v), we observed a clear enrichment of genes encoding for protein folding and degradation machineries and for protein kinase A binding (GO:0019900, GO:0051018, **Extended Data** Fig. 11d**, Supplementary Table 1**). For example, the Heat Shock Protein 90 Alpha Family Class B Member 1 (Hsp90ab1, Fig. 1e**, Supplementary Table 1**), increases its level in mRNA during aging but is downregulated at the protein level (v). We found that genes involved in the chaperone complex and several metabolic and catabolic processes (GO:0101031, GO:0016054, GO:0044283, **Extended Data** Fig. 11d**, Supplementary Table 1**) showed a down-up mRNA pattern and an up-down protein pattern (vii), suggesting that part of the proteostasis imbalance observed during aging is due to an inefficient compensation of these machineries at the protein level. For example, the gene NADH:Ubiquinone oxidoreductase complex assembly factor 2 (Ndufaf2, Fig. 1e**, Supplementary Table 1**), which is involved in mitochondrial complex I assembly, showed this pattern.

Examples of genes decreasing during aging in mRNAs but with increasing protein levels (vi) or showing up-down mRNA and down-up protein patterns (viii) include NADH:Ubiquinone oxidoreductase subunit s4 (Ndufs4, Fig. 1e**, Supplementary Table 1**), another mitochondrial complex I protein^42^, and neurofilament medium chain (Nefm, Fig. 1e**, Supplementary Table 1**). This finding was supported by enrichment of GO terms such as ‘intermediate filament-based process’ and ‘actin cytoskeleton’ (GO:0045103, GO:0015629, **Extended Data** Fig. 11d**, Supplementary Table 1**). These findings suggest that for the remaining ∼50% of gene products the proteo-transcriptome is much more complex than expected, prompting additional analyses. In addition, having perfectly matched datasets allowed us to perform novel isoform discovery, where after combining information from transcriptomics and proteomics we were able to identify novel isoforms expressed at different ages, including two novel proteoforms for synaptotagmin 7 (Syt7), which regulates asynchronous neurotransmitter release (see Methods, **Supplementary text 1 and Extended Data** Fig. 12).

### Protein-mRNA (p/m) coefficient suggests loss of molecular regulation in aging

To obtain further understanding of mRNA-protein discrepancies, we calculated the protein-mRNA coefficient (p/m coefficient, **Supplementary Table 2**) which provides a single value to describe the relationship between mRNA and protein levels. We first checked the distribution of p/m coefficients and observed that it became narrower with age (Fig. 1h), as evidenced by a significant reduction in the full width at half maximum (FWHM, Fig. 1i) of the peaks and a decrease in the absolute distance from the Z-score center (Fig. 1j). The narrowing was accompanied by an increase in correlation between protein and mRNA levels with aging (Fig. 1k, Spearman ρ correlation (median) 6m: ∼0.37, 12m: ∼0.40 and 24m: ∼0.41). As the variability in the p/m coefficient goes down with age, the correlation between protein and mRNA levels increases. This suggests that expression is dynamically regulated in young brains, with multiple mechanisms influencing translation and degradation. However, with aging, these processes may become less efficient, leading to a higher correlation between mRNA and protein levels and resulting in a loss of fine-tuned gene expression control, potentially causing a flattening of expression regulation in aged brains.

### Co-regulation modules reveal hidden alterations of mitochondrial, synaptic, ribosomal and proteasomal genes within the aging mouse brain

To obtain a biology-oriented understanding of how p/m coefficients change we evaluated p/m co-regulation using weighted gene co-expression network analysis (WGCNA)^43^. This method allows us to divide gene products with similar expression patterns into modules. We identified eight mutually exclusive modules of different sizes ranging from 42 to 1001 genes (Fig. 2a, b**, Extended Data** Fig. 13 **and Supplementary Table 2**) with two that were increased at 12m and 24m compared to 6m but decreased at 24m compared to 12m (M1, M2, M3, M7 Fig. 2c).

**Figure 2:**
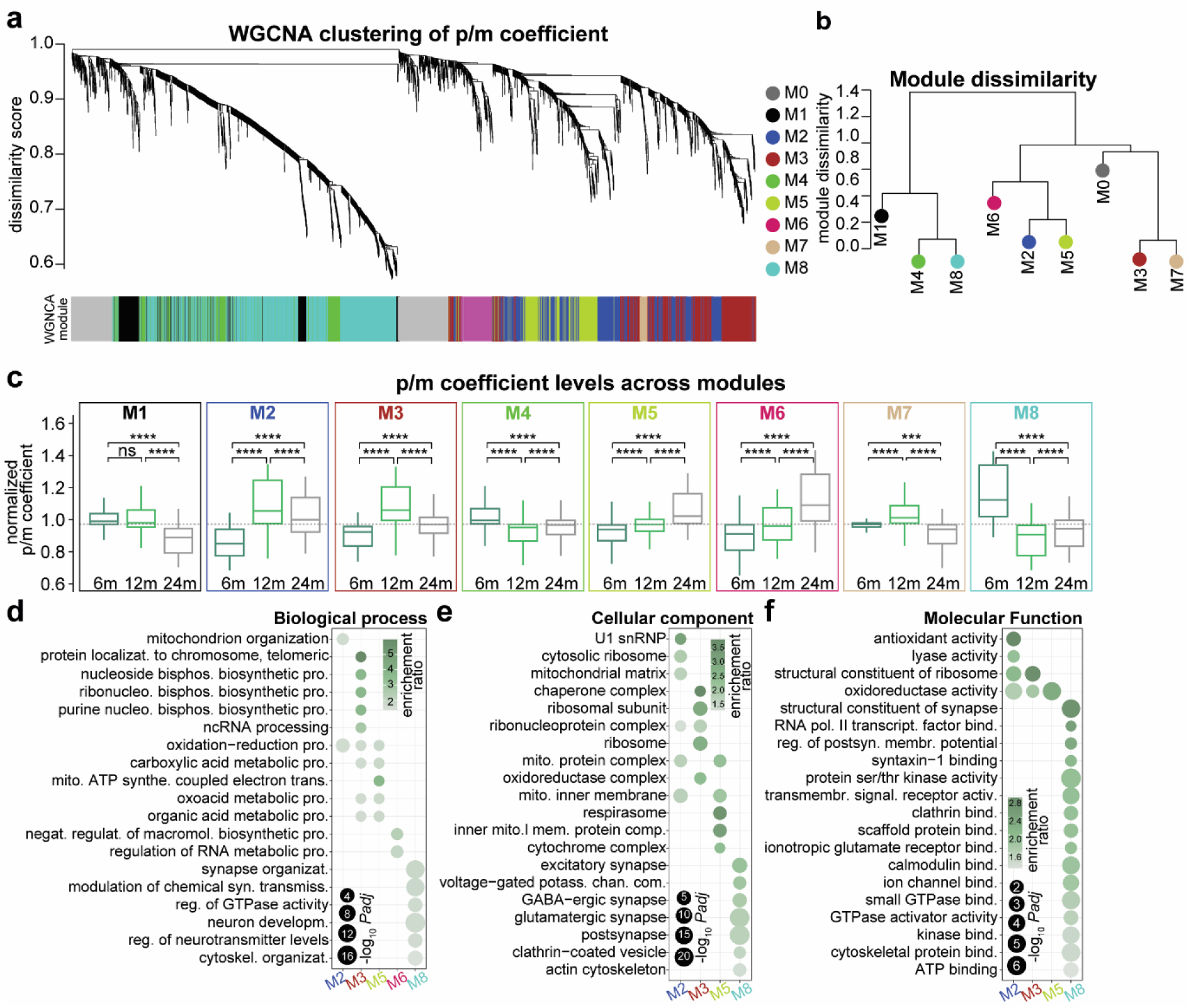
The analysis of the p/m coefficient identifies altered proteo-transcriptome modules in the aging mouse brain. (**a**) WGNCA dendrogram with highlighted modules (lower colored bar). Genes were clustered based on dissimilarity measures. The branches are modules of closely correlated proteo-transcriptome groups that have a similar p/m coefficient. Eight significant modules and M0 corresponding to ∼3500 genes were detected with WGCNA. M0 is a module with a less correlated gene group. (**b**) Module dissimilarity based on module eigengene distances. Modules are grouped based on their p/m coefficient. (**c**) Boxplots of normalized p/m coefficient in 6m, 12m and 24m mouse brain, grouped by their modules detected in the WGNCA method. Tukey post hoc test P-value * ≤ 0.05, ** ≤ 0.01, *** ≤ 0.001 and **** ≤ 0.0001. (**d-f**) Gene Ontology over-representation analysis for the five selected modules (M2, M3, M5, M6 and M8). For all modules in detail refer to **Supplementary Table 2**. Dot sizes in the enrichment graphs correspond to the *padj* and color scales represent the enrichment ratio for each GO term, dark green – highly enriched and gray – less enriched.

The p/m coefficients for modules M4 and M8 were decreased at 12m and 24m compared to 6m but increased at 24m compared to 12m (Fig. 2c). Modules M5 and M6 were continuously increased with aging (Fig. 2c). Over-representation analysis (ORA) using GO annotations identified several significantly enriched biological processes for each module (Fig. 2d**-f****, Supplementary Table 2**), allowing to isolate modules related to mitochondrial and ribosomal function (M2, M3, M5), RNA splicing (M6) and neuronal and synaptic biology (M5, M8). Although the data for all modules is available (**Supplementary Table 2**), due to space limitations, we focused on the four modules with the most striking patterns: (i) p/m coefficient for the M2 module was increased at 12m and 24m compared to 6m but decreased at 24m compared to 12m. In other words, the protein produced per unit of mRNA increased at 24m and 12m compared to 6m but decreased at 24m compared to 12m.

This module strikingly includes ribonucleoproteins and mitochondrial genes such as Ndufa10, Ndufa11, Ndufa12, Ndufa13, Ndufa8, Ndufaf2, Ndufb7, Ndufs3, and Ndufs6, which are all components of the mitochondrial complex I. Here, the p/m coefficient is dominated by the protein levels, since the mRNA levels show an opposite pattern (**Extended Data** Fig. 14a, b). Module M3 (ii) is similarly increased at 12m and 24m compared to 6m but decreased at 24m compared to 12m in the aged brain and includes a large number of ribosomal and chaperone genes that are important for maintaining protein homeostasis such as Cct2, Cct3, Cct4, and Cct5 of the chaperone CCT complex and Cops3, Cops4, and Cops5 of the COP9 signalosome which showed a similar trend as the module (**Extended Data** Fig. 15a**, Supplementary Table 3**). Module M5 (iii), which shows a continuous increase in p/m coefficient in the aging brain, encodes several components involved in metabolic and catabolic processes in the mitochondria. In the M5 module the protein produced per unit of mRNA decreases with aging (iv) while for M8 the p/m coefficient is decreased at 12m and 24m compared to 6m but increased at 24m compared to 12m in the aged brain. M8 includes genes related to cytoskeletal organization and synaptic function such as major structural components of the neuronal cytoskeleton, Nefh, Nefl and Nefm^44^ (Fig. 2c**-f**). Together, these changes show that aging disrupts the coordinated regulation of mitochondrial, ribosomal, proteasomal, and synaptic genes in the brain with dynamic alterations in protein-to-mRNA ratios that may impact neuronal homeostasis.

We then explored the association of these modules with human pathologies by curating genes associated with brain disease (**Extended Data** Fig. 15b). This analysis showed that module M8, related to synaptic function, are clearly over-represented in brain diseases. Furthermore, the synaptic module M8 shows an increase in p/m correlation with age (Spearman ρ correlation (median) 6m: ∼0.37, 12m: ∼0.38 and 24m: ∼0.40) similar to the overall trend, whereas we observed a decrease in the mitochondrial modules M2 and M5 (**Extended Data** Fig. 15c, M2 - Spearman ρ correlation (median) 6m: ∼0.37, 12m: ∼0.36 and 24m: ∼0.37 and M5 - Spearman ρ correlation (median) 6m: ∼0.62, 12m: ∼0.62 and 24m: ∼0.61). Taken together, these results suggest that the synaptic genes are dysregulated at both the mRNA and protein levels, but there may be discrepancies for the mitochondrial genes.

### Changes in p/m coefficient and alterations in mitochondrial and synaptic genes

Given the previously reported age-related dysfunction in mitochondrial and synaptic function^1,4,11,12,14,15,17,18,39–41,45,46^ and the finding that changes in the modules M2, M5 and M8 directly point to these alterations, we decided to study in more detail how the p/m coefficient for mitochondrial and synaptic genes changes during aging. For this purpose, we curated two specific gene lists using the MitoCarta^47^ for mitochondrial genes and the SYNGO^48^ database for synaptic genes. Considering these two groups, we found a remarkable switching behavior during aging (Fig. 3a**-c**). At 6m the synaptic genes show overall high p/m coefficient, possibly suggesting very efficient translation (Fig 3a**; Extended Data** Figure 16a-c**; Extended Data** Figure 17a-d**; and Supplementary Table 3**). This was evident for genes encoding key neuronal components such as potassium channels (Kcna2, Kcnd3), calcium channels (Cacna1e, Cacna2d1), components of the NMDA receptor complex (Grin2a), and glutamate receptors (Grik3). In contrast, mitochondrial genes showed a low p/m coefficient indicating a relatively lower translation at 6m when compared to other ages (Fig 3a**; Extended Data** Figure 16a-c**; Extended Data** Figure 17a-d**; and Supplementary Table 3**). This was observed for mitochondrial ribosomes (Mrpl1, Mrps9, and Mrpl13).

**Figure 3:**
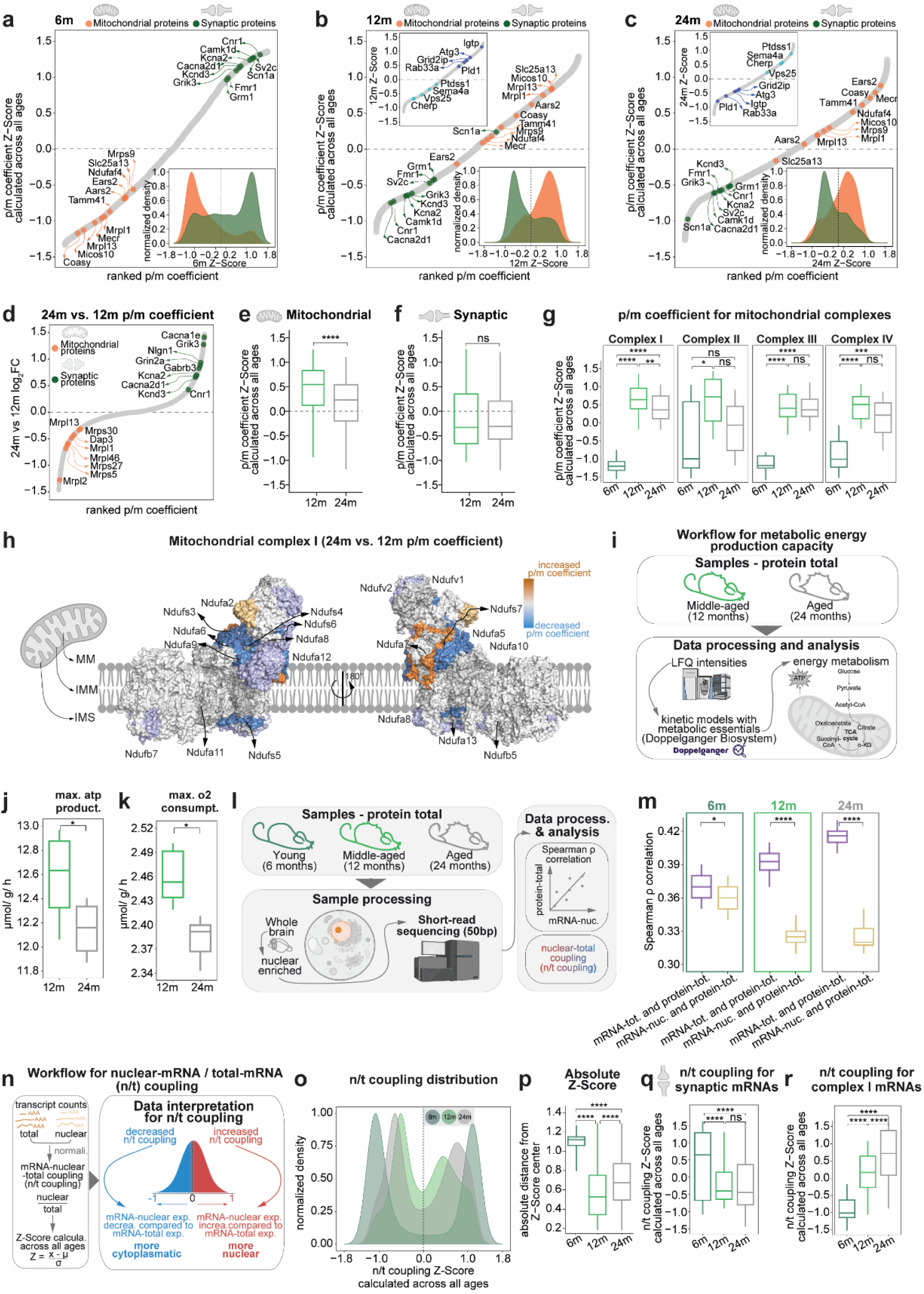
p/m coefficient and n/t coupling changes in the aging mouse brain – alterations in mitochondrial and synaptic genes. (**a-c**) Scatter plots for the genes ranked by their p/m coefficient for (**a**) 6m, (**b**) 12m and (**c**) 24m using the Z-Score calculated across all ages. Insets are density distribution of p/m coefficient for 6m, 12m and 24m for mitochondrial^47^ (orange) and synaptic^48^ (green) proteins. (**d**) Scatter plot for the genes ranked by their p/m coefficient for 24m vs. 12m using the log_2_FC between 24m and 12m. Positive values indicate an increased p/m coefficient in 24m, and negative value is a decreased p/m coefficient at 24m. (**e,f**) Boxplots of the (**e**) mitochondrial and (**f**) synaptic proteins for 12m and 24m. P-values indicate the results of paired t-test. (**g**) Boxplot for p/m coefficient for mitochondrial complexes. P-values indicate the results of paired t-test followed by Tukey post hoc test. (**h**) Structural representation for the genes from mitochondrial complex I for 24m vs. 12m using the log_2_FC between 24m and 12m (**i**) Schematic of the experimental workflow assessing the metabolic energy production capacity in 12m and 24m using protein-total dataset. Boxplots for (**j**) Mitochondrial ATP production capacity and (**k**) Mitochondrial oxygen consumption rate. P-values indicate the results of paired t-test. (**l**) Schematic of the experimental and analytical workflow assessing mRNA-nuclear fraction of aging mouse brains across 6m, 12m, and 24m. (**m**) spearman ρ correlation for mRNA-total and protein-total compared to ρ for mRNA-nuclear and protein-total for each age. P-values indicate the results of paired t-tests. (**n**) Workflow for analysis and categorization of nuclear-total mRNA coupling (n/t coupling) in aging brain. Positive coupling values (red) indicate increased nuclear-mRNA levels relative to total-mRNA, suggesting increased nuclear localization or these mRNAs are more in the nuclear; negative values (blue) indicate decreased nuclear-mRNA levels relative to total-mRNA, indicating decreased nuclear localization or more cytoplasmic. (**o**) Density distribution of p/m coefficient for 6m, 12m and 24m. (**p**) Absolute distance from Z-Score center for the density distribution for 6m, 12m and 24m in Fig. 3o. P-values indicate the results of paired t-test followed by Tukey post hoc test. (**q,r**) Boxplots for n/t coupling for (**q**) synaptic mRNAs and (**r**) mitochondrial complex I mRNAs. P-values indicate the results of paired t-test followed by Tukey post hoc test P-value * ≤ 0.05, ** ≤ 0.01, *** ≤ 0.001 and **** ≤ 0.0001.

Interestingly, we found that at 12m and 24m the synaptic genes showed the opposite pattern with decreased p/m coefficient for synaptic and increased for mitochondrial (Fig. 3b, c**; Extended Data** Figure 16a-c**; Extended Data Figure 17e; and Supplementary Table 3)**.While these changes appear to occur similarly in 12m and 24m old mice compared to 6m mice (suggesting a phenotype arising already at middle age), we were interested in understanding what happens at later aging stages when mice start showing memory deficits^49^. We thus compared specifically 24m vs. 12m and found that at 24m the mitochondrial genes showed a decreased p/m coefficient (Fig. 3d, e). This was also evident for several complex I genes (Fig. 3g), including 10 Ndufs proteins that showed a decreased p/m coefficient at 24m compared to 12m (Fig. 3h). The protein level changes for complex I followed a similar trend as the p/m coefficient, while the mRNA expression showed an opposite trend, suggesting that the system tries to a certain extent to compensate for the decreased levels of protein (**Extended Data** Fig. 18a). Overall, these results indicate that the mitochondrial proteins produced per unit of mRNA was significantly reduced at 24m compared to 12m and point to inefficient translation. In general, synaptic genes as a whole did not show significant differences at 24m vs 12m, even though we found few examples such as Grid2ip, Rab33a which are involved in synaptic biology with decreased p/m coefficient (Fig. 3b, c **inset and** Fig. 3d). As an important note, we also identified genes involved in immune system regulation and signaling (Ptdss1, Sema4a, Vps25, and Cherp; Fig. 3b, c **inset**) which showed an increased p/m coefficient at 24m compared to 12m, suggesting increased translational output for these genes.

Given the obvious importance of mitochondrial Complex I in energy metabolism, we asked whether the small but widespread differences in protein abundances that we observed at 24m vs 12m could lead to substantial age-specific changes in mitochondrial energy metabolism. To this end, we used an *in silico* approach that predicts mitochondrial dysfunction based on protein levels^50–52^ (Fig. 3l). This analysis revealed that the identified protein level changes can lead to significantly decreased ATP production (Fig. 3m **and Supplementary Table 4**) and oxygen consumption (Fig. 3n, o **and Supplementary Table 4**). Collectively, these results suggest that there is an energy imbalance due to a small but widespread decrease in mitochondrial complex I protein levels that is attempted to be compensated for increased mRNA levels (which are increased, **Extended Data** Fig. 18a).

### Nuclear RNA analysis reveals age-related defects in transcript localization

We then specifically investigated whether mRNA export from the nucleus could allow us to more meaningfully interpret changes in p/m ratios. We thus isolated nuclear fractions from whole mouse brains from the same exact tissue used for bulk mRNA and protein measurements and prepared mRNA from isolated nuclei (Fig. 3l). RT-qPCR and western blot validated the efficiency of our fractionation protocol. For instance, Malat1 and Lamin A/C were as expected, localized to the nucleus and cytosolic enriched mt-Co1and α-Tubulin showed an enrichment in their respective fraction (**Extended Data** Fig. 19a, b). We additionally compared our results to the published SINC-seq data^53^ and the RNALocate database^54^ (**Extended Data** Fig. 19c, d) and observed a clear enrichment of the nuclear mRNAs validating the reliability of our sequencing results. Extensive analysis on differential expression analysis can be found in the **Supplementary Text 2 (Supplementary Table 5)**.

To analyze the effect of nuclear mRNAs on proteostasis we first evaluated the correlation of nuclear mRNA abundance and protein abundance. This analysis indicated that, while during aging there is an increase in the correlation between total mRNA abundance and protein abundance (as seen in Fig. 1k), the correlation between nuclear mRNA abundance and protein abundance is substantially decreased in the aged brain (Fig. 3m). One possible interpretation is that mRNAs that are not efficiently transported outside the nucleus in the aged brain do not get efficiently translated into proteins. Moreover, this may also suggest a shift toward more nuclear mRNA accumulation with advancing age.

To more clearly address discrepancies between nuclear mRNA and total mRNA, we calculated the nuclear-mRNA vs total-mRNA coupling (n/t coupling, Fig. 3n**, Supplementary Table 5**) which provides a single value to describe this relationship (see methods). We first checked the distribution of the n/t coupling and observed that it has a pronounced bimodal distribution that becomes narrower at 24m and 12m compared to 6m (Fig. 3o) Given our previous observation of decreased protein production efficiency for mRNAs at 24m and 12m compared to 6m (Fig. 3a**-h**), we checked the n/t coupling for synaptic and mitochondrial mRNAs using Z-Scores calculated across all ages. For synaptic genes, we observed a significant decrease in n/t coupling at 24m and 12m compared to 6m, but no significant difference at 24m compared to 12m. This indicates that nuclear localization does not lead to late-aging effect on synaptic mRNAs, rather indicating that synaptic mRNAs tend to be enriched outside the nucleus already at 12m. On the contrary, by examining changes in n/t coupling for the mRNAs encoding mitochondrial proteins in the nucleus, we found that the mRNAs for complex I and IV showed a significant increase in n/t coupling at 24m compared to 12m (Fig. 3p**-r****; Extended Data** Fig. 24a**; and Supplementary Table 5**). Taken together, these results highlight age-specific changes in the nuclear export for nuclear encoded mitochondrial mRNAs which could affect their availability and translation efficiency.

### Age-related dysregulation of mTOR signaling

Analysis of mTOR-related genes such as Ak1, Tsc1, and Tsc2 showed a decreased p/m coefficient **(Extended Data** Fig. 25a). The decreased p/m coefficient indicates a decreased protein production per unit of mRNA, in other words a decreased protein expression and increased mRNA expression at 24m **(Extended Data** Fig. 25b, c). Hspa5, important for ER folding showed a decreased p/m coefficient while Eif2a showed an increased p/m coefficient **(Extended Data** Fig. 25a), which indicates an activation of the integrated stress response and suggests a compensatory mechanism to the possible ER stress in the aged brain^55^.

### Age-related changes in protein solubility and dysregulation of mTOR signaling

Obvious possible protein complexes implicated in proteostasis regulation are ribosomes and the proteasomes. When considering the p/m coefficient for these complexes we observed an up-down trend (Fig. 4a**, Supplementary Table 3**), suggesting a complex dynamic regulation during aging. Given the tendency to accumulate misfolded proteins during brain aging^56^, we decided to consider the SDS-insoluble fraction of the proteome to quantitatively picture this regulation and we performed LC-MS/MS to measure protein abundances in the SDS-insoluble fractions (see methods) for the same exact samples for which we previously measured mRNA (total and nuclear) and total protein abundances (Fig. 4b). To confirm the enrichment of the SDS-insoluble fraction we relied on comparisons with published datasets^11^ and confirmed reliable enrichment of aggregated proteins (**Extended Data** Fig. 26a, b). Additional detailed analyses revealed several interesting findings discussed in the supplements (**Extended Data** Fig. 26d **and Supplementary Table 6**: see also **Supplementary text 3**).

**Figure 4:**
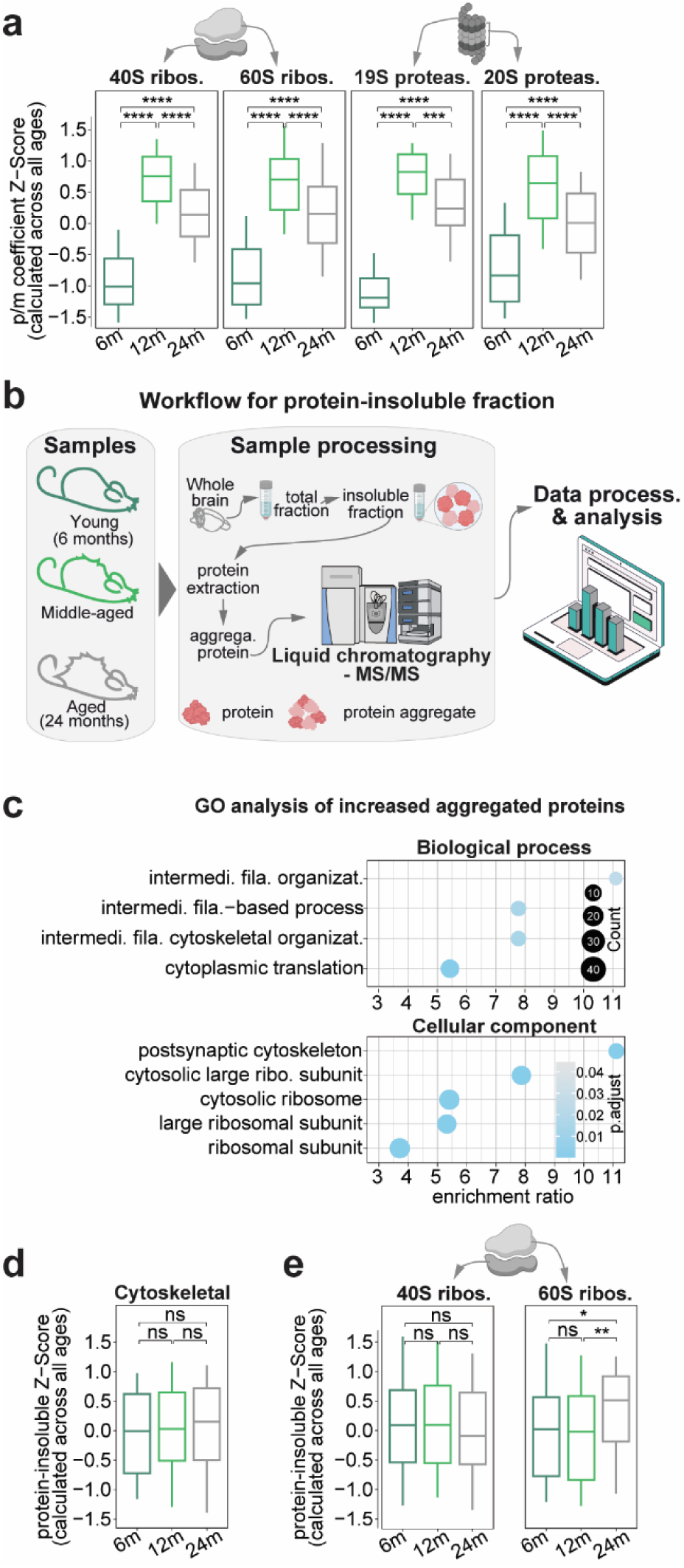
Shifts in-age related changes in protein solubility. (**a**) Boxplot for p/m coefficient for ribosomal and proteasome complexes. P-values indicate the results of paired t-test followed by Tukey post hoc test. (**b**) Schematic of the experimental and analytical workflow assessing protein-insoluble fraction of aging mouse brains across 6-month (6m), 12-month (12m), and 24-month (24m). (**c**) GO-ORA of the top150 proteins that are upregulated in 24m vs 12m in the protein aggregate fraction. (**d,e**) Boxplots for protein aggregate for, (**d**) Cytoskeletal proteins and (**e**) 40S and 60S ribosome. P-values indicate the results of paired t-test followed by Tukey post hoc test P-value * ≤ 0.05, ** ≤ 0.01, *** ≤ 0.001 and **** ≤ 0.0001.

A clear aspect that could be observed in the SDS-insoluble fraction during late aging (24m vs 12m) is that neurofilaments (*i.e.*, Nefl, Nefh, and Nefm) and ribosomal proteins become less soluble with age, as also highlighted by GO analysis (Fig. 4c). A more general assessment of all cytoskeletal proteins did not show this phenotype, confirming specificity for neurofilaments (Fig. 4d). Detailed analysis of ribosomal proteins indicates that the large subunit is the one clearly affected during late aging (Fig. 4e).

### In late aging, changes in biochemical parameters suggest hampered translation and metabolic imbalance

We reasoned that an increase in aggregation of 60S subunits might cause shortage of functioning ribosomes and lead to less efficient translation. Since positively charged amino acids, such as lysine, are known to interact with the ribosome’s exit tunnel^57^, the translation of proteins rich in positive charges should be especially slow during the elongation process and less favorable. In other words, if translation is impeded by non-functional 60S subunits in the aged brain, any further deferral from a “sticky” positive nascent peptide can make the process worse.

As a result, mRNAs encoding proteins with a high content of positively charged residues should have a relatively lower p/m coefficient at 24m vs. 12m (since for the same amount of mRNA the protein produced should be less abundant). To test this hypothesis and, in general to explore the possibility that simple biochemical parameters can influence proteo-transcriptome balance during late brain aging, we computed several biochemical protein features (**Supplementary Table 7**) and tested whether any of these were linked to changes in the p/m coefficient in late aging (Fig. 5a). This analysis revealed that the relative percentage of positive amino acids in the protein sequence are negatively correlated to p/m coefficient in late aging (Fig. 5b **and Extended Data** Fig. 24), with lysine content indeed being the likely cause of this effect (Fig. 5c).

**Figure 5:**
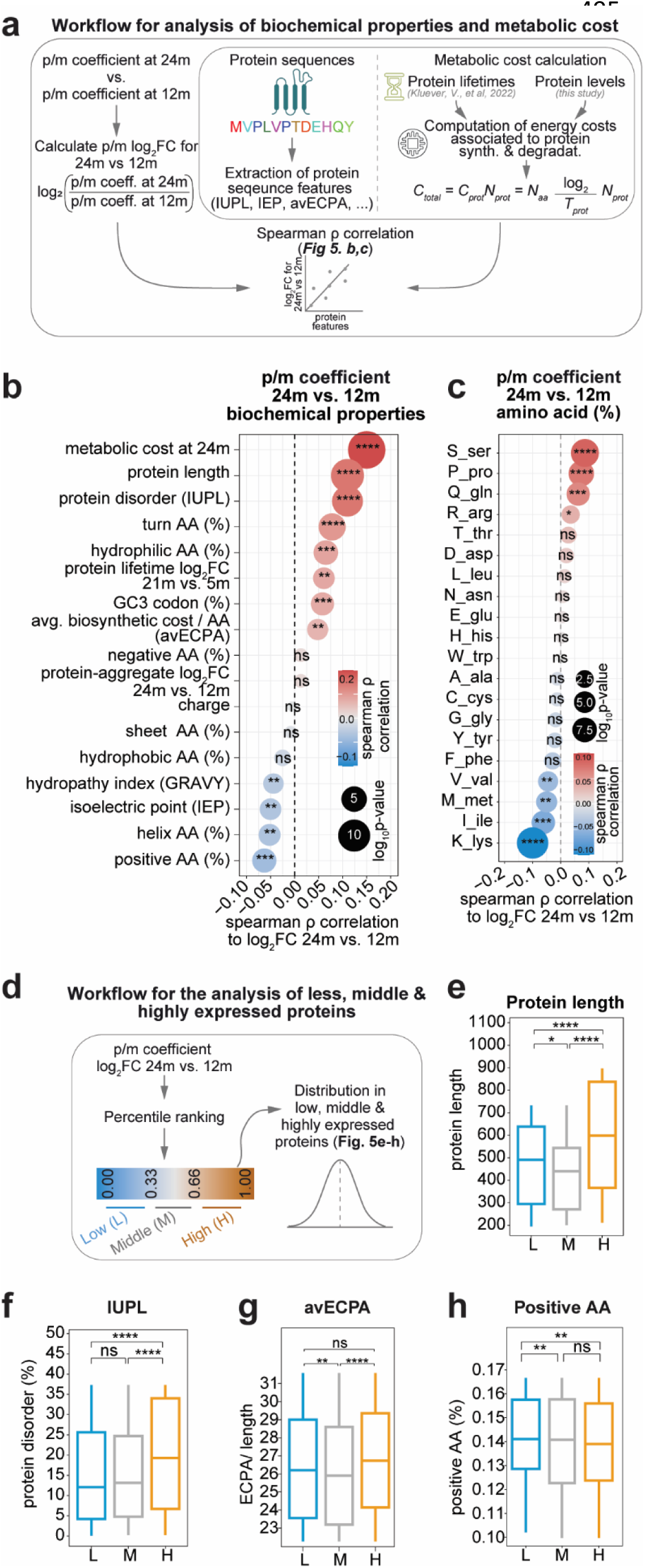
Biochemical property alterations and translational energy imbalance in the aged brain. (**a**) Analytical workflow for the correlation between protein biochemical properties, metabolic cost (see methods for details and p/m coefficient in late aging. (**b,c**) Spearman ρ correlation for 24m vs. 12m to (**b**) biochemical properties and (**c**) amino acid percentage. Colors indicate Spearman ρ correlation, where red signifies positive correlation, and blue signifies negative correlation. (**d**) Percentile ranking of the log_2_FC of the p/m coefficient for the 24m vs. 12m analysis of low (blue; 0.00 to <0.33), medium (gray; 0.33 to ≤ 0.66) and high (orange; >0.66 and ≤1.00) expressed proteins. (**e-h**) Boxplot for proteins for high (H, increased), low (L, decreased) and middle (M, unchanged) in 24m compared to 12m. (**e**) Protein length, (**f**) average biosynthetic cost per amino acid (avECPA) normalized by protein length similar to our previous work^5^.(**g**) Protein disorder using Intrinsically Unstructured Protein-Like (IUPL), and (**h**) Amino acid percentage for positive amino acids. P-values for paired t-test followed by Tukey post hoc test. P-value * ≤ 0.05, ** ≤ 0.01, *** ≤ 0.001 and **** ≤ 0.0001. (**i**) Workflow for the model to predict the metabolic cost calculations.

Following a hint by previous proteomic work from our lab^5^ we considered the normalized biosynthetic cost of amino acid synthesis (avECPA) and we noticed that there is a significant positive correlation between aminoacidic biosynthetic cost and p/m coefficient suggesting that translation of the “more expensive” proteins is favored in late aging. These results were largely recapitulated when we considered the relative difference in p/m coefficient after grouping (Fig. 5d) suggesting that specifically those whose increased p/m coefficient is higher in late aging are those mostly relevant for these differences (Fig. 5e**-h**).

We thus decided to explore whether protein species with an overall higher metabolic burden than other protein species would behave differently in the aged brain. To do so, we calculated the metabolic cost per proteo-transcriptome species based on the protein turnover parameters that we measured in a previous work^5^ and combined the protein-level measurements obtained in this study. Our findings revealed a positive correlation between higher metabolic cost per proteo-transcriptome species and the p/m coefficient in late aging. This suggests that high-cost proteins experience an increase in translational efficiency with age (Fig. 5b). However, we observed contrasting changes in overall aging (i.e. 6m vs. 24m) indicating this is a late-aging effect (see **Supplementary Text 4** for details).

Collectively, these findings highlight an age-specific shift in metabolism which reshapes the proteome and can be at least in part reconducted to simple biochemical properties, such as protein size and charge.

### Identifying resilience and compensations in physiological brain aging by comparative proteomics of eight additional mouse models and their controls

The molecular changes observed in the aging brain so far reflect an intricate interaction between metabolic constraints and proteome re-organization, which could also potentially include resilience processes against the deterioration caused by advanced age. If biochemical properties, such as protein size and charge, influence translational efficiency and metabolic cost, then comparing proteomic changes in physiological aging with mouse models of neurodegenerative diseases (NDD) and accelerated aging (aAGE) could allow to identify possible differences and identify where resilience mechanisms might emerge and whether they are linked to possible protective adaptations against neurodegeneration.

To address this, we performed an additional large set of comparative proteomic measurements and established an extensive database for total and aggregated proteins for eight cohorts of mice and their matched controls. These models and ages were chosen based on their lifespan and genetic background to capture age-related changes but preceded the most severe pathological changes (Fig. 6a **and Extended Data** Fig. 30). Each mouse model was compared to a matched control line to avoid biases due to genetic background variability. The eight cohorts can be divided into two sub-groups of aAGE and NDD. The aAGE mice included: (1) the Senescence-Accelerated Mouse Prone 8 (SAMP8, aAGE_1, phenotypic aging^58,59^), (2) the AIRAPL-deficient (aAGE_2, proteostasis aging^60^), (3) the Lamin A knock-in (LAKI, aAGE_3, nuclear aging^61^), (4) the Zmpste24-deficient (aAGE_4, nuclear aging^62^). The NDD mice included: (1) the APPPS1 (NDD_1, AD mouse model^63^), (2) the Line 61 (NDD_2, PD mouse model^64^), (3) the HM2 (NDD_3, PD mouse model^65^), and (4) the UCP2 knockout (NDD_4, PD mouse model^66^).

**Figure 6:**
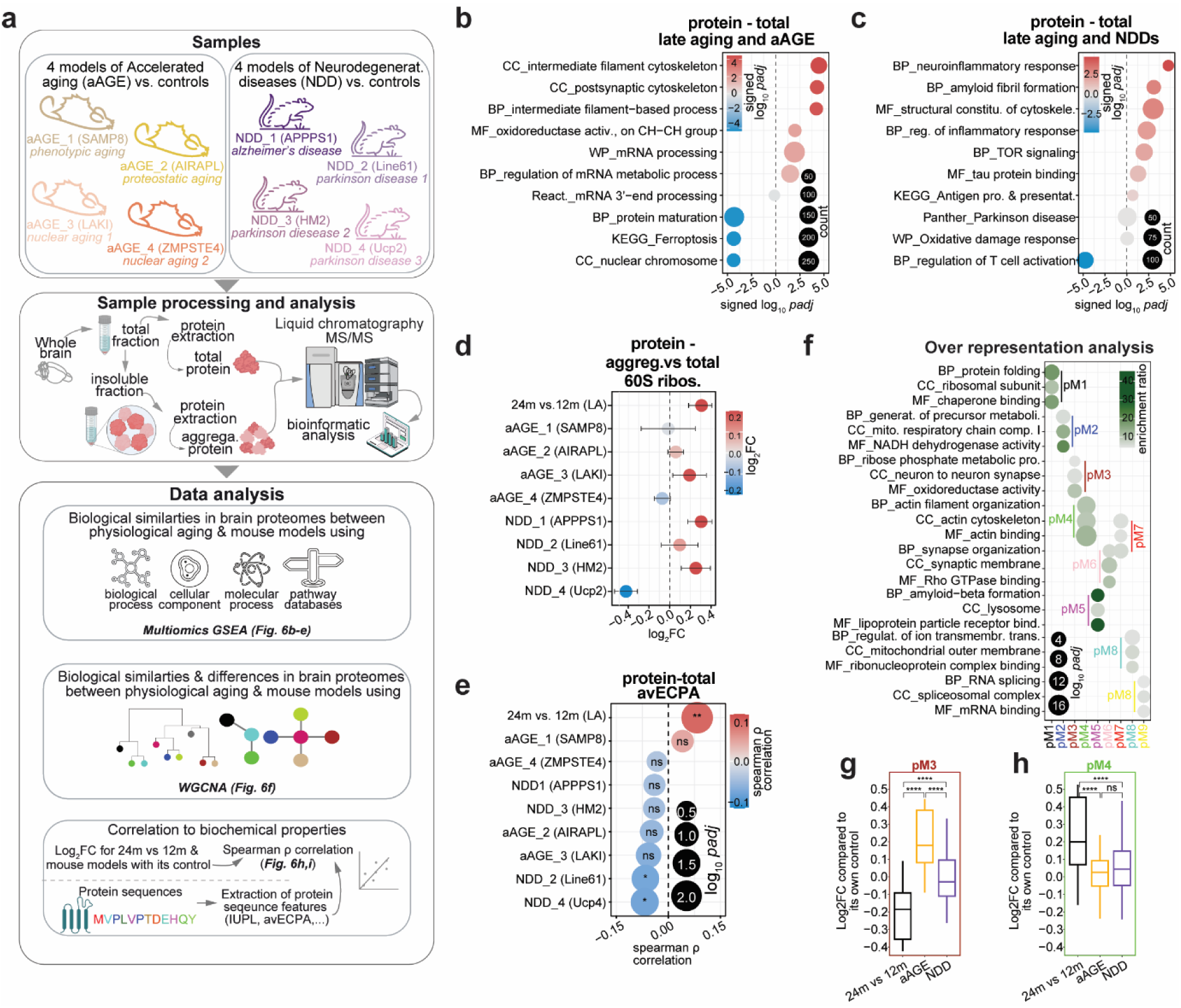
Identification of shared protein abundance changes in late aging, accelerated aging, and neurodegenerative disease mouse models. **(a)** Schematic of the experimental and analytical workflow assessing protein-total and -insoluble fraction of mouse models for accelerated aging (aAGE) and neurodegenerative diseases (NDD). aAGE models include SAMP8 (aAGE_1), AIRAPL (aAGE_2), LAKI (aAGE_3) and ZMPSTE4 (aAGE_4). NDD models include APPPS1 (NDD_1), Line61 (NDD_2), HM2 (NDD_3) and Ucp2 (NDD_4). (**b,c**) Multi-list GSEA for protein total fraction comparing (**b**) late aging and aAGE and (**c**) late aging and NDD (**Supplementary Table 9**). Positive *padj* values (red) indicate increased enrichment; negative *padj* values (blue) indicate decreased enrichment. The size of the dots corresponds to count, bigger the size of the bubble means they are a greater number of genes. (**d**) Dot plot for proteins in 60S ribosome for protein-aggregate fraction normalized by the protein-total fraction. (**e**) Spearman ρ correlation for 24m vs. 12m, aAGE and NDD to avECPA in protein-total fraction. Colors based spearman ρ correlation, where red signifies positive correlation, and blue signifies negative correlation. (**f**) Gene ontology over-representation analysis (GO-ORA) for the modules identified using WGCNA. Here are the top 3 GO terms based on the enrichment ratio. For all modules in detail refer to **Supplementary Table 10**. Dot sizes in the enrichment graphs correspond to the padj and color scales represent the enrichment ratio for each GO term, dark green – highly enriched and gray – less enriched. (**g, h**) Boxplot for log_2_FC in 24m vs. 12m, average of aAGE and NDD models for (**g**) pM3 and (**h**) pM4 modules. P-values for paired t-test followed by Tukey post hoc test. P-value * ≤ 0.05, ** ≤ 0.01, *** ≤ 0.001 and **** ≤ 0.0001

Having collected these datasets, we first performed differential protein expression analysis in the aging brain by comparing protein abundance to their control obtained with mass spectrometry. Due to space limitations all these analyses can be found in the supplementary section of the work (**Supplementary Text 5**; **Extended Data Figs 31** and **Supplementary Table 9**). In short, important pathways that were affected included gliosis and inflammation (Gfap, Vim) that are upregulated in late physiological aging (24m vs 12m), several aAGE models, and NDD models. Among other notable proteins we found a cytoskeletal maintenance protein (Plec), components of the protein homeostasis machinery (Sae1, Hsp90ab1), and proteins related to lipid metabolism (Osbpl1a, Oxct1) that showed notable differences (See **Supplementary Text 5** for details).

Next, using multi-omics gene set enrichment analysis (GSEA), we looked at similarities between physiological late aging and either aAGE models or NDD models (Fig. 6b**,c****; Supplementary Table 9**). We found as an example that late aging and aAGE models have a similar increase in cytoskeletal proteins and decrease in protein maturation co-factors (Fig. 6b).

Late aging also shared with NDD models an increased inflammatory response (coupled with decreased T cell activation) and increased amyloid fibril formation tendency (Fig. 6c). Analysis of shared pathways in SDS-resistant proteins normalized by the total protein indicated that the aggregated phenotype of 60S ribosomes is conserved in 6 out of 8 models identifying a clear common alteration observed in brain aging (Fig. 6d). We also checked if the avECPA that we observed positively correlated to the p/m coefficient in late aging (Fig. 5b; possibly suggesting a more efficient translation of longer transcripts with high biosynthetic cost), was correlated to increased protein levels (Fig. 6e). Only late aging showed a significant positive correlation that was not observed in the mouse models, suggesting that increased translation of longer transcripts with high biosynthetic cost may be a resilience mechanism unique to physiological aging.

Following a similar reasoning we performed WGCNA for models of late aging based on total protein measurements, to identify possible changes that are put in place by physiological aging to compensate maladaptive alterations (**Extended data** Fig. 32a**, Supplementary Table 10**). This analysis identified nine mutually exclusive modules (**Extended Data** Fig. 32a), which include protein folding (pM1), mitochondrial changes (pM2 and pM8), metabolic and synaptic processes (pM3), cytoskeletal organization (pM4), amyloid beta (pM5), synaptic and cytoskeleton (pM6 and pM7) and mRNA metabolism (pM9). We identified the modules in which changes in the total protein abundance were clearly different in late aging compared to both sets of mouse models (see **Extended data** Fig. 32b and Fig. 6g, h). We reasoned that possible resilience mechanisms in late aging might be absent from these mouse models as they would constitute pathological adaptations in aAGE and NDD mice. This analysis revealed that both pM3 and pM4 have clear significant changes setting late aging apart from aAGE and NDD models (Fig. 6g, h). In detail pM3, the metabolic and synaptic module, show decreased protein levels in physiological aging while pM4, the cytoskeletal module, is increased. Overall, this suggests that aging follows trajectories that might have protective effects for maladaptive alterations.

Thus, we reasoned that if these modules indeed mirror underlining possible resilience adaptations that are put in place during aging, then examining protein levels across all ages would show non-linear dynamic trajectories during aging (Extended data Fig. 32c). Indeed, these two modules clearly show such trajectories, with pM3 increasing at 12m and decreasing at 24m and pM4 showing the opposite trend (Extended Data Fig. 32c). Interestingly, we checked and found an increase of neurofilament proteins (Nefh, Nefl, Nefm) among the proteins of pM4, while pM3 contained metabolic and synaptic proteins (Ogdh, Pdhx, Homer3). Furthermore, the pM4 module was positively correlated with protein length (ρ = 0.22, *p.value* = 0.03) and avECPA (ρ = 0.24, *p.value* = 0.02) and pM3 was negatively correlated with LA (protein length: ρ = −0.2, *p.value* = 0.03; avECPA: ρ = −0.2, *p.value* = 0.03) and no significant differences for aging models (**Supplementary Table 10**). From these results we could speculate that the proteins in pM4 and pM3 could be involved in a potential resilience mechanism with aging.

## Discussion

The brain undergoes extensive molecular changes during aging^67,68^, that affect its function and increase its susceptibility to age-related neurodegenerative diseases^69,70^. How different molecular layers interact and influence each other in the aging brain remains elusive. In this study we comprehensively analyzed four molecular layers in parallel, namely mRNA and protein abundance, nuclear mRNA abundance, and levels of SDS-insoluble (aggregated) proteins in young, middle-aged and aged mice. By combining eight additional proteomic datasets of mouse model of disease and accelerated aging, applying careful data curation and detailed bioinformatic analyses, we performed a comprehensive characterization of the proteo-transcriptome in the aging brain. This large database (BrainAging-MAP; Fig. 7a**, Extended Data Figs. 33-37**) can be easily explored by the scientific community.

**Figure 7:**
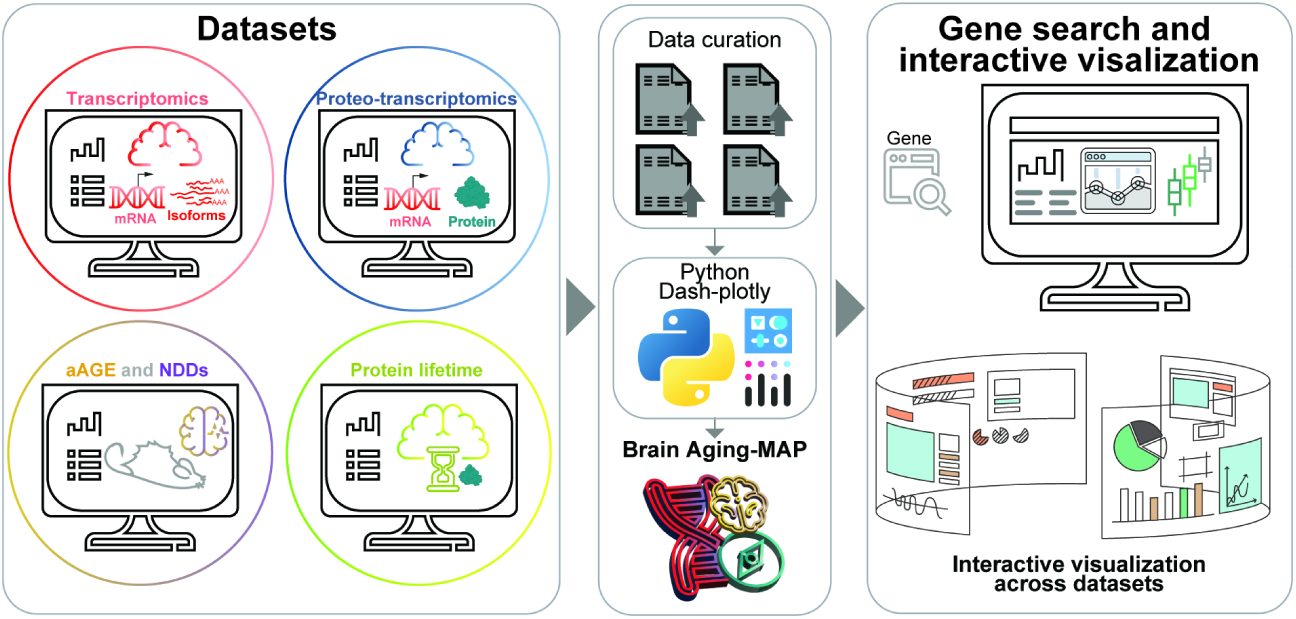
Overview of the BrainAging-MAP dash application. A web application for multi-layered proteo-transcriptomic datasets.

Several interesting observations arise from our analyses. We have confirmed widespread low levels of inflammation and gliosis ^71–73^, highlighted by an upregulation of astrocyte-specific gene such as GFAP (Fig. 1e). These are known hallmarks of brain aging and our results are well in line with previous findings^27,74^.

Further, we observed here opposing translational trajectories of synaptic and mitochondrial proteins. While synaptic proteins that are efficiently translated in young brains become translationally constrained with age, mitochondrial proteins follow the opposite trajectory. This possible tug of war for resources is a fundamental new finding. Mitochondrial impairment is a hallmark of the aging brain^1,7,26,39–41,45^ but our study points to a specific decrease in translation for mitochondrial complex I proteins and an increased nuclear retention of the mRNA for these proteins in late aging (Fig. 3r). This indicates that the molecular defect is not just a loss of mitochondrial gene expression, but an inability to translate key components efficiently, which disrupts ATP production and oxygen consumption as also supported by other works^75,76^. This challenges the conventional view that transcriptional compensation is sufficient for maintaining mitochondrial function in aging and highlights translational control as a critical failure point.

Synaptic dysfunction is another key feature in the aging brain^26,39–41^, in our study we show that synaptic protein-mRNA balance already changes strikingly between 6m and 12m but does not significantly worsen at 24m. This suggests that neurons undergo an early-stage molecular synaptic reorganization in mid-life, followed by a plateau, rather than a linear decline. This contrasts with mitochondrial proteins, which continue to deteriorate after 12m, implying that synapses adapt to an extent, but the adaptation does not occur for mitochondria. A possible explanation could be that in middle age there is a reallocation of energy resources to mitochondrial maintenance instead of translation of synaptic proteins. However, this compensation mechanism fails at 24m, which is supported by a relative decrease in the p/m coefficient at 24m vs. 12m and is specifically evident for mitochondrial complex I proteins.

Protein degradation and synthesis are affected in the aging brain^5,11,77^. This is supported by a reduction in proteasomal and ribosomal activity, and in particular an aggregation of ribosomes, which are conserved in the aging brain in both killifish and mouse^11^, similar to our observations in late aging (Fig. 4). Our study reveals that the 60S ribosome is predominantly involved in this process. Moreover, ribosomal subunit aggregation is also observed in several mouse models, suggesting an important role in aging and neurodegeneration. This is further supported by a differential regulation of the mTOR pathway, indicating a modulation in protein synthesis. Furthermore, several studies have reported alterations in translation control mechanisms in the aging brain^11,78,79^. Our data also supports this by the tightening of the protein-mRNA (p/m) coefficient with age (Fig. 1 h-j), suggesting that with aging comes at the cost of a loss of regulation to differentially regulate gene expression leading to a paradoxical increase in mRNA-protein correlation (Fig. 1k). In line with this, some of the captivating findings of our work are that the positively charged amino acids show a relatively lower p/m coefficient at 24m vs. 12m (Fig. 5 b,h), suggesting that translation of proteins rich in positive charges might specifically slow down the elongation process. The proteins that are longer and have high biosynthetic costs (Fig. 6e) are efficiently translated with aging, but not in aAGE or NDD models, suggesting the existence of a resilience mechanism in physiological aging and indicating a possible reallocation of metabolic resources that prevents pathological changes that would lead to neurodegeneration.

In conclusion, our work collectively challenges the notion that age-related dysfunction is solely due to transcriptional changes and positions translational control and proteome prioritization as a possible therapeutic target for age-related neurodegeneration. Overall, our data serves as a solid foundation for more systematic pathway-oriented work that will elucidate the molecular causes of brain deterioration during aging.

## Methods

### Animal cohorts

All locally conducted animal experiments (physiological aging cohort) were approved by the Lower Saxony State Office for Consumer Protection and Food Safety (Niedersächsisches Landesamt für Verbraucherschutz und Lebensmittelsicherheit). Mouse lines are frequently employed in aging research because of their capacity to produce disease-free and genetically consistent animal cohorts^5,24,80^. In both mice and humans, adolescence is marked by several changes in synaptic refinement and brain connection. The human brain reaches full cerebral maturity by the end of the second decade of life, whereas the mouse brain continues to mature until six months of age^81–83^. To compare them to the older mice, we used 6-month-old (6m) mice as a control group in the present study. Furthermore, aging is a nonlinear process, and as significant changes in humans take place in the fourth to sixth decade of life ^84^, it is essential to research the middle aged and aged adult window to comprehend the nonlinear aging process. Whereas, in mice this transitional window would correspond between ages of 12-month-old (12m) and 24-month-old (24m)^85^ Therefore, we used 12m mice as a middle-aged group to compare with young (6m) and older (24m) mice to explain the early and late events in the aging process. We also compared 24m to 6m to explain the overall aging process. We decided to concentrate on male mice, similar to what we did in the past^13,24,86^. The cohorts of mice used for physiological aging (C57Bl/6JRj) were purchased from Janvier labs and routinely checked for absence of the following four classes of pathogens: <u>1) Bacteria and fungi</u>: *Bordetella bronchiseptica*, CAR bacillus, *Citrobacter rodentium*, *Clostridium piliforme*, *Corynebacterium bovis*, *Corynebacterium kutscheri*, *Dermatophytes*, *Encephalitozoon cuniculi*, *Helicobacter* spp, *Mycoplasma pulmonis*, *Pasteurellaceae*, *Actinobacillus* spp., *Haemophilus* spp., *Mannheimia haemolytica*, *Pasteurella* spp., *Pasteurella multocida*, *Pasteurella pneumotropica*, *Pasteurella trehalosi*, *Salmonella* spp., *Streptobacillus moniliformis*, *Streptococci ß-hemolytic* (not group D), *Streptococcus pneumoniae*; <u>2)</u> <u>Endoparasites</u>: Protozoa, *Entamoeba* spp, Flagellates, Coccidia, Helminths, Cestodes, Nematodes; <u>3) Ectoparasites</u>: Mites, Fur-dwelling mites, Surface-dwelling mites, Follicle-dwelling mites, Lice, Fleas; <u>4) Viruses</u>: Hantaviruses, K virus (Mouse pneumonitis virus), Lactate deshydrogenase elevating virus (LDV), Lymphocitic choriomenigitis virus (LCMV), Minute virus of mice (MVM), Mouse adenovirus (MAD) type 1 (FL), Mouse adenovirus (MAD) type 2 (K87), Mouse cytomegalovirus (MCMV), Mouse hepatitis virus (MHV), Mouse parvovirus (MPV), Mouse polyomavirus, Mouse rotavirus (EDIM), Mouse thymic virus (MTV), Mousepox (Ectromelia) virus, Murine norovirus (MNV), Pneumonia virus of mice, Reovirus type 3 (Reo 3), Sendai virus, Theiler’s murine encephalomyelitis virus (TMEV).

All mouse models were obtained from the collaborators and have been previously described in published studies, as cited below. We grouped the mouse models used in this study into accelerated aging (aAGE) and neurodegenerative diseases (NDDs). The accelerated aging models include: senescence-accelerated prone mouse 8 (SAMP8) and SAMR1 (aAGE_1)^58,59^, AIRAPL (aAGE_2)^60^, LmnaG609G/G609G (LAKI, aAGE_3)^61^, and Zmpste24-/- and Zfand2b-/- (Zmpste24 KO, aAGE_4)^62^. The NDD models include APPPS1 (NDD_1)^63^, WT-aSyn(Line 61, NDD_2)^64^, A30P (HM2, NDD_3)^65^ and the UCP2-/-(Ucp2 KO, NDD_4)^66^. All models were used with their respective littermate controls.

### Tissue isolation and protein extraction

Upon sacrifice, brains were quickly removed on ice, snap-frozen in liquid nitrogen and stored at −80°C until further processing. Before processing for biochemistry and sequencing, the olfactory bulbs were removed from the frozen brains, and one hemisphere, alternating left and right, was selected for lysate preparation. The biochemical fractionation protocol was optimized based on the protocol by Bandopadhyay (2016)^87^. Hemispheres were washed in 320 mM sucrose buffer to remove blood and superficial vessels, then homogenized in 2 mL of 320 mM sucrose buffer containing protease inhibitors (cOmplete, Roche) and phosphatase inhibitors (PhosSTOP, Roche). Homogenization was carried out using a Teflon pestle with ten strokes at 900 rpm. To the homogenate, 10X RIPA lysis and extraction buffer (ThermoFisher) was added to achieve a final concentration of 1X RIPA. Benzonase (Merck) was then added, and the samples were mixed on a rotation mixer for 30 minutes at room temperature (RT). Following this lysis and extraction process, samples were centrifuged at 2000 g at 4°C to remove cell debris. The supernatant was transferred to a new tube, and 20% SDS was added to achieve a final concentration of 5% SDS. The samples were then rotated again for 30 minutes at RT. A 400 µl aliquot was collected for the ‘total’ fraction, and the remaining sample was ultracentrifuged at 100,000 g for 30 minutes at RT. The supernatant was collected as the ‘soluble’ fraction, while the pellet was resuspended in 5% SDS and placed on a shaker at 800 rpm for 30 minutes, with intermittent vigorous pipetting to dissolve the ‘insoluble’ fraction. Protein concentration for all samples was estimated using the BCA assay.

### Trypsin digestion and SP3 cleanup

For SP3 cleanup and trypsin digestion, protein samples were adjusted to 50 µg, following the protocol by Hughes et al. (2019). The samples were heated to 95°C with RapiGest SF (Waters), reduced with dithiothreitol (DTT; Sigma-Aldrich), and alkylated with iodoacetamide (IAA; Sigma-Aldrich) while shaking at 600 rpm. RapiGest was then diluted before the addition of magnetic carboxylate-modified beads (Sera-Mag SpeedBeads; GE Healthcare). The SP3 cleanup procedure was performed according to^88^, with minor modifications. Specifically, 25 µg of beads were used per 50 µg of protein. Protein binding to the beads was induced by adding ethanol to a final concentration of 50%, followed by washes with 80% ethanol using a magnetic rack. After the ethanol was removed, 200 µl of digestion solution containing 1 µg of trypsin in 100 mM ammonium bicarbonate buffer was added. Samples were briefly sonicated and incubated overnight at 37°C while shaking. The next day, samples were centrifuged, separated on a magnetic rack, and the supernatant was collected and dried using a SpeedVac vacuum concentrator (Eppendorf).

### Peptide library preparation for BoxCar

To generate a peptide library for spectral matching we used the protocol by Meier and colleagues^89^, digested peptides from the total, soluble, and insoluble fractions of different animals and mouse models were pooled. To enhance the depth of LC-MS analysis, these pooled peptide samples were fractionated offline by high-pH reverse-phase chromatography. Fractionation was performed using an Agilent 1100 HPLC system, a Waters XBridge BEH C18 column, and an in-house constructed fraction collector rotating every minute, yielding 10 or 20 concatenated fractions. The flow rate was set at 60 µl/min. Peptides were separated using a linear gradient from Buffer A (10 mM ammonia in water) to Buffer B (10 mM ammonia in 80% acetonitrile) up to 70% Buffer B. Collected fractions were dried in a SpeedVac and reconstituted in LC-MS/MS loading buffer (0.05% trifluoroacetic acid, 5% acetonitrile in water).

### Analytical proteomics and LC-MS/MS analysis

For LC-MS/MS analysis, peptides were resuspended in loading buffer (0.05% trifluoroacetic acid, 5% acetonitrile in water), sonicated, and approximately 1 µg of peptide per sample was injected in duplicate (technical replicates) into an UltiMate 3000 RSLCnano HPLC system coupled to a Q Exactive HF mass spectrometer. Peptides were first desalted on a reverse-phase C18 pre-column and then separated over 88 minutes on a 30 cm ReproSil-Pur C18 AQ 1.9 µm reverse-phase analytical column. MS data acquisition was performed using a standard data-dependent acquisition (DDA) method, scanning precursor ions from 350 to 1600 Da at a resolution of 60,000 at m/z 200, with an automated gain control (AGC) target of 3 × 10^6. The top 30 precursor ions were selected for MS2 at a resolution of 15,000 at m/z 200, with a maximum ion injection time (IT) of 50 ms and an AGC target of 1 × 10^6. Alternatively, the BoxCar method was used with MaxQuant Live^38^. MS1 resolution was set to 120,000 across an m/z range of 300 to 1650, with a maximum IT of 250 ms and an AGC target of 3 × 10^6. MS2 was acquired at a resolution of 15,000 with a maximum IT of 28 ms and an AGC target of 1 × 10^6. Preset BoxCar settings included 3 scans, 12 BoxCar boxes, and a 1 Thomson overlap. The datasets generated and analyzed are available in PRIDE under accession number PXD030904.

### Label-free quantification of proteins

Mass spectra were analyzed using MaxQuant v1.6.17.0, with BoxCar selected as the search mode, and the high-pH reverse-phase fractionated sample data utilized as the peptide library. Label-free quantification (LFQ) was performed. The mouse reference proteome was obtained from UniProt in August 2020. Search parameters were selected based on Meier et al. (2018). The ‘match between runs’ feature was enabled with a 0.5-minute window following retention time alignment, and the ‘LFQ min ratio’ was set to 1. MS/MS spectra were not required for LFQ. The false discovery rate (FDR) for peptide and protein identification was set to 0.01. Methionine oxidation and N-terminal acetylation were included as variable modifications, while carbamidomethylation was set as a fixed modification. The first search tolerance was set to 20 ppm, and the main search tolerance was set to 4.5 ppm. The minimum peptide length was set to seven amino acids, with a maximum peptide mass of 4600 Da.

### Filtering and normalization for proteomic datasets

For data preprocessing, we applied stringent filtering criteria to ensure the quality and reliability of our proteomic dataset. Proteins were retained for analysis if they exhibited at least 50% valid values across biological replicates across all conditions. We excluded from the analysis known contaminant blood proteins such as hemoglobin subunits and albumin (Hbb-b1, Hbb-b2, Hbb-y, Hbb-bh1, Hbz, Hbb-bh0, and Alb), to minimize potential confounding factors. Additionally, proteins flagged as ‘+’ in the columns “Reverse”, “Potential Contaminant”, and “Only Identified by Site” were removed from further analyses. To address systematic biases and enhance inter-sample comparability, we implemented a two-step normalization process. First, quantile normalization was applied to align the distribution of protein abundances across all samples, effectively mitigating technical variations. Subsequently, we performed row-wise median normalization to facilitate robust comparisons across biological replicates. This comprehensive normalization approach ensured that downstream analyses were conducted on a dataset optimized for biological interpretation and statistical reliability.

### Parallel reaction monitoring (PRM) for confirmation of selected targets level changes

The SpikeTides L (JPT, Berlin) were solubilized as instructed by JPT. Shortly, each well containing a heavily labelled peptide 100 µl of solubilization buffer containing 20% Acetonitrile (ACN) and 80% 0.1M ammonium bicarbonate (ABC) were added. This was incubated for 10 minutes at RT. Subsequently, the buffer was pipetted up and down and then the solubilized peptides were transferred to Eppendorf cups. For the following analyses, 10 µl of each peptide were mixed and the volume of the mixture was brought to 100 µl by adding solubilization buffer. From this, 50 aliquots containing approximately 20 (peptides are not accurately quantified) pmol of each heavy peptide were prepared. The stocks and the aliquots were then lyophilized and stored at −20°C.

The tryptic-digested and SP3 cleaned up mouse brain samples were desalted and further cleaned up by using Harvard Apparatus C18 micro SpinColumns. After cleaning, samples were lyophilized. For Mass Spectrometry (MS) analysis, the heavy peptides were solubilized first using loading buffer (4% ACN and 0.1% trifluoroacetic acid (TFA)). Concentration of heavy peptides were adjusted to 10 fmol per 7 µl. The heavy peptide solution was then used to solubilize the sample containing the endogenous peptides derived from mouse brains (final concentration: 10 fmol of heavy peptide and 1 µg of endogenous peptides in 7 µl).

The samples were measured on an Orbitrap Exploris™480 mass spectrometer (Thermo Fisher Scientific). Peptide were separated on a self-packed RP C18 analytical column (75 µm inner diameter, ReproSil-PurC18-AQ3 resin) packed into a Silica Tip emitter (FS360–50-8-N, New Objective). The peptide mixture was separated over a 58 minute 0-42% ACN gradient and analyzed via an acquisition method containing a MS1 survey scan and targeted MS2 scans. For MS1 scans, the Orbitrap resolution was 120,000 FHWM, the scan range (*m/z*) was adjusted to the *m/z* range of the peptides to be detected, a standard AGC target was applied, and the mass spectrometric analysis was performed in positive mode. For the MS^2^ scans, the isolation window (*m/z*) was set to 1.4 *m/z*, a normalized collision energy of 30% was applied, a normalized AGC target of 100% implemented, the maximum injection time was adjusted to 100 ms, an Orbitrap resolution of 60,000 FHWM was used, and data was acquired in positive mode. For targeting, the retention time, charge state and mass of the peptides were defined in the PRM methods. Retention time windows of 8 minutes were used. The retention time windows were newly determined for each set of measurements, as the retention time of the peptides can be easily influenced by e.g. buffer change in the system.

One part of data analysis was performed using the software Skyline^90^ (version 21.1.0.278). The peptide sequences can be imported into Skyline, the isotope modifications for ^13^C, ^15^N-labelled arginine and lysine can be added in the settings section of the program and instrument and measurement details can be adjusted in the transition section of the program. When Skyline is provided with all the information about peptides and measurement settings, the MS raw files can be imported into Skyline. Retention times of the LC-MS runs were aligned. Endogenous and labelled signals were compared, transitions from endogenous peptides were only considered as such, when they had the same retention time and the same fragmentation pattern as their labelled counterpart. For further analyses, only peaks were considered, which showed at least three transition ions and three to five data points. The total integrated area of the light and heavy peaks was then used to calculate the ratios between light and heavy peptide.

Normalization of the data was performed using the total ion current (TIC) area from each measurement. For this, the TIC area of one measurement was set to 1 and based on that, a normalization factor was calculated for every measurement. The total area of the peaks corresponding to the endogenous peptides were then divided by their according normalization factor

### mRNA isolation

RNA was isolated in parallel from the matched hemisphere of the “physiological aging” cohort, from which the protein data was generated, to ensure perfect matching. The protocol for obtaining total and nuclear fractions was adapted from Gagnon, Song and their colleagues^91,92^. Briefly, brain tissue was washed in 320 mM sucrose buffer (320 mM sucrose, 5 mM HEPES, pH 7.4) and homogenized using a Teflon pestle (10 strokes at 900 rpm) in 2 mL of sucrose buffer supplemented with RNase inhibitor. The lysate was filtered through a 100 µm cell strainer, washed with 2 mL of 2× hypotonic lysis buffer (HLB; 10 mM Tris, pH 7.5, 10 mM NaCl, 2 mM MgCl₂, 0.3% NP-40, and 10% glycerol), briefly vortexed, and incubated on ice for 10 minutes. A 500 µL aliquot was taken for ‘total’ RNA, mixed with 800 µL Trizol LS, and stored at −80°C overnight. The remaining lysate was adjusted to 7 mL with HLB and centrifuged at 1,000g for 3 minutes at 4°C. The cytosolic fraction (2 mL of the supernatant) was collected and stored at −20°C. The remaining supernatant was discarded, and the nuclear pellet was washed twice more in HLB, followed by centrifugation at 1,000g for 2 minutes. After removing the supernatant, the pellet was resuspended in 1.8 M sucrose buffer (1.8 M sucrose, 50 mM Tris, pH 7.5, 1 mM MgCl₂) and transferred to an ultracentrifugation tube. The sample was ultracentrifuged at 70,900g for 60 minutes at 4°C, after which the upper phase was removed. The nuclear pellet was washed and resuspended in 320 mM sucrose buffer, mixed with 800 µL Trizol LS, and stored at −80°C for at least one night. RNA was extracted using the Qiagen RNeasy Mini Kit according to the manufacturer’s instructions. RNA was maintained on ice during processing and stored at −80°C.

### Library preparation and mRNA sequencing

RNA sequencing was performed using Illumina’s next-generation sequencing technology. Total RNA was quantified and assessed for quality using a Tapestation 4200 instrument with RNA ScreenTape (Agilent Technologies). For library preparation, 500 ng of total RNA was used with the NEBNext Ultra II Directional RNA Library Preparation Kit, combined with the NEBNext Poly(A) mRNA Magnetic Isolation Module and NEBNext Multiplex Oligos for Illumina (96 Unique Dual Index Primer Pairs), following the manufacturer’s instructions (New England Biolabs). Libraries were quantified and assessed for quality using the Agilent 4200 Tapestation Instrument and D1000 ScreenTapes (Agilent Technologies). The prepared libraries were pooled and sequenced on an Illumina HiSeq4000 with single-end reads of 50 bp. Sequence data were converted to FASTQ format using bcl2fastq v2.20.0.422. The datasets generated and analyzed are available in the Gene Expression Omnibus (GEO) under accession numbers GSE249499 (total fraction, 6m, 24m^24^), GSE278427 (total fraction, 12m) and GSE278509 (nuclear fraction, 6m, 12m and 24m).

### RNA-Seq data processing and quantification

The quality of raw sequencing data was assessed using FastQC v0.11.91^93^. Reads were then aligned to the GRCm39.103 reference mouse genome using HISAT2^94^. Gene-level read counts were quantified with featureCounts from the Subread package^95^. Low-count genes (≤5 reads) were filtered out, followed by quantile normalization and row-wise median normalization to facilitate cross-replicate comparisons. Principal component analysis was performed in R using the prcomp() function to visualize the normalized data and detect potential outliers. No outliers were identified in this quality control step.

### Differential protein expression and gene ontology analysis for proteomics datasets

To account for variability in peptide quantification, we utilized both quantile-normalized LFQ intensities and Razor+unique peptides for differential protein expression (DPE) analysis using the DEqMS package version 1.10.0^96^ in R. Proteins with significant differential expression (**Supplementary Table 1, 6 and 9**), defined by p-values ≤ 0.05, were selected for further analysis. To determine the functional significance of these proteins, we performed gene ontology analysis using the clusterProfiler package v4.5.0^97^. Gene over-representation analysis was conducted with the enrichGO() function using default parameters, applying a p-value cutoff of ≤ 0.05. P-values were adjusted using the Benjamini–Hochberg method (**Supplementary Table 1, 6 and 9**). The background set for over-representation analysis consisted of genes identified as significantly differentially expressed.

### Differential gene expression and gene ontology analysis for transcriptomics datasets

Differential gene expression (DGE) analysis was performed using DESeq2 v1.30.1^98^ in R, utilizing raw count data. The package’s internal median ratio method was employed for normalization. Genes were considered significantly differentially expressed if they met the following criteria: adjusted p-value (*padj*) ≤ 0.05 and absolute log2 fold change (|log2FC|) ≥ 0.58 (**Supplementary Table 1, 5**). Functional analysis of the differentially expressed genes was conducted using the clusterProfiler package v4.5.0^97^. Gene ontology (GO) over-representation analysis was performed using the enrichGO() function with default parameters. A p-value cutoff of ≤ 0.05 was applied, and p-values were adjusted using the Benjamini– Hochberg method (**Supplementary Table 1, 5**). The background gene set for the over-representation analysis consisted of all genes identified as significantly differentially expressed.

### Gene ontology analysis

ORA was conducted using the enrichGO() function from the clusterProfiler package v4.5.0^97^ with default parameters. A significance threshold of p ≤ 0.05 was applied, with p-values adjusted using the Benjamini-Hochberg method. Gene Set Enrichment Analysis (GSEA) was performed using the gseGO() function, also from clusterProfiler, with default parameters. The significance threshold was set at p ≤ 0.05, and p-values were adjusted using the Benjamini-Hochberg method.

For multi-list Gene Ontology (GO) analysis, we utilized the multi-list utility in WebGestalt^99^ graphical user interface to conduct ORA and pathway analysis (**for** Fig 6b, c). This approach allowed for the integration of multiple gene lists, enabling a comprehensive analysis across different datasets.

### mRNA-total and protein-total data filtering

We filtered the datasets to ensure high-quality gene expression profiles for further analysis. For the mRNA datasets, we applied a threshold of standard error (STDERR) < 0.12, while for the protein dataset, we used a threshold of STDERR < 0.4. This filtering process resulted in a refined set of ∼3500 genes, which were then used for subsequent analyses. These stringent criteria allowed us to focus on genes with reliable expression measurements across both mRNA and protein levels, enhancing the robustness of our findings in the context of age-related expression dynamics

### Temporal dynamics analysis

To identify gene groups with similar or divergent trends in mRNA-total and protein-total fraction datasets during physiological aging, we employed a Pearson correlation analysis. For each gene, we calculated the correlation between its Z-score and an idealized trend vector representing various expression patterns across the 6-, 12-, and 24-month time points. We defined eight distinct trends: four with similar expression patterns in both mRNA and protein datasets (continuous upregulation, continuous downregulation, and two non-linear patterns: ‘down-up’ and ‘up-down’; i-iv), and four with divergent patterns between mRNA and protein datasets (up-transcript and down-protein, down-transcript and up-protein, down-up transcript and up-down protein, up-down transcript and down-up protein v-viii). For each trend, we created an idealized vector (e.g., 0, 0.5, and 1 for continuous upregulation at 6, 12, and 24 months, respectively) and calculated the Pearson correlation coefficient between each gene’s Z-score and this vector for both mRNA and protein datasets. Genes were then assigned to the trend group with the highest correlation coefficient, allowing us to categorize genes based on their expression dynamics during aging in both mRNA and protein levels (used in Fig. 1d-f).

### Novel proteoforms analysis

First, splice junctions were identified using the command line tool rMATs^100^ in the transcriptomic datasets of the 6m and 24m old brains^24^, as these were paired-end with a read length of 150bp. These were later quantified as alternative splicing (AS) events such as mutually exclusive exon (MXE), retention intron (RI), skipped exon (SE), alternative splicing at the 5’ or 3’ splice site (A5SS, A3SS). Second, based on the splice sites identified in the transcriptomics datasets, the reads were filtered by junction sites, later the translation sites were identified using JCAST^101^ and translated to create a custom protein database. Third, the custom protein database was searched against the uniport Mus musculus sequence using blastp and only proteoforms with approximately less than 90% similarity were used to ensure that they were non-cannonical isoforms. Fourth, the raw files from the proteomics datasets were re-quantified using the custom protein database created from the transcriptomics datasets. Fifth, the LFQ intensities for the proteoforms were used to perform the differential protein expression analysis using the DEqMS package version 1.10.0^96^ in R (used in Extended Data Fig. 12).

### Protein-mRNA coefficient (p/m coefficient)

To assess protein-to-mRNA (p/m) coefficient, we first applied percentile rank normalization to both protein and mRNA datasets. For each biological replicate, we computed the p/m ratio, which we termed “p/m coefficient”. This ratio data underwent a subsequent renormalization step to prepare it for further analysis. We then calculated Z-scores for each biological replicate based on p/m coefficient. This process involved computing the mean and standard deviation of p/m coefficient across all ages and replicates. For each gene and biological replicate, we subtracted this mean from the individual p/m coefficient value and divided by the standard deviation:

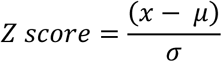

Where x is the individual p/m coefficient value, μ is the mean, and σ is the standard deviation. The resulting Z-score reflects the overall change in p/m coefficient for each gene across all ages, with a score of 0 indicating no difference between groups. Positive or negative values represent the direction and magnitude of the change in p/m coefficient.

### Weighted correlation network analysis (WGCNA) analysis of p/m coefficient

To construct co-expression networks that capture genes with similar p/m coefficient in the aged brain, we first normalized the count data using percentile rank normalization, followed by row-wise median normalization to adjust for protein-specific differences in the aged brain. The normalized gene-level counts were then used as input for identifying functional groups with the WGCNA v1.7.1^43^ R package.

We created p/m coefficient modules within a signed network based on the topological overlap matrix (TOM), applying a soft-threshold power of 14 to approximate a scale-free topology. Modules were defined with a minimum size of 30 and a cutHeight of 0.97. Proteins were clustered based on dissimilarity (1-TOM), and similar modules were merged using the mergeCloseModules() function with a cutHeight of 0.26 (**Supplementary Table 2,** Fig. 2**, Extended Data** Fig. 13**, 14**).

To evaluate the functional significance of these modules, we conducted a gene over-representation analysis using the enrichGO() function from the clusterProfiler v4.5.0^97^ R package. The background gene set included all genes assigned to any module in the WGCNA analysis, ensuring a thorough assessment of functional enrichment (**Supplementary Table 2**). We also explored the relationship between these modules and brain pathologies by analyzing curated disease datasets. The pathologies considered were Parkinson’s disease (PD, N = 229), mental retardation (MR, N = 457), autism (N = 924), ataxia (N = 245), amyotrophic lateral sclerosis (ALS, N = 50), and Alzheimer’s disease (AD, N = 82). Overlap ratios were computed using the GeneOverlap v1.26.0^102^ package in R.

### Nuclear vs. total mRNA coupling (n/t coefficient)

To evaluate the coupling between nuclear and total mRNA (n/t), we implemented a stringent filtering process. We initially focused on approximately 3,000 genes detected in the nuclear fraction. These genes were selected based on two criteria: (1) they met the standard error (STDERR) threshold established for the mRNA-total and protein-total datasets, and (2) they exhibited a minimum count of 5 in the nuclear fraction. This approach ensured the inclusion of reliably quantified transcripts, minimizing potential artifacts from low-abundance or poorly detected mRNAs. We then applied percentile rank normalization to both nuclear and total fraction mRNA datasets. For each biological replicate, we computed the n/t ratio, which we termed “n/t coupling”. This ratio data underwent a subsequent renormalization step to prepare it for further analysis. We then calculated Z-scores for each biological replicate based on n/t coupling. This process involved computing the mean and standard deviation of n/t coupling across all ages and replicates. For each gene and biological replicate, we subtracted this mean from the individual n/t coupling value and divided by the standard deviation:

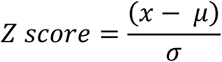

Where x is the individual n/t coupling value, μ is the mean, and σ is the standard deviation. The resulting Z-score reflects the overall change in n/t coupling for each gene across all ages, with a score of 0 indicating no difference between groups. Positive or negative values represent the direction and magnitude of the change in n/t coupling.

### Biochemical properties and protein feature analysis

In our analysis of biochemical properties and protein features (Fig. 6), we employed a custom Python script to extract amino acid sequences and compute sequence length and average amino acid compositions from UniProt^103^ identifiers. We used the ProtParam module in Biopython^104^ to calculate isoelectric point, secondary structure and GRAVY (Grand Average of Hydropathy) score. The disordered fraction (IUPL) was determined using IUPred^105^, while the intrinsically disordered score was acquired from MobiDB^106^. For each protein sequence, we calculated the average biosynthetic cost (avECPA) using previously described amino acid cost values ^107^. Spearman’s rank correlation coefficients (ρ) were then computed to assess the relationships between these properties and features (used in Fig. 5a, b).

### Model calculations for metabolic costs

The energetic costs associated to mRNA and protein synthesis and degradation (used in Fig. 5a) are computed following Marino, Bergmann and collegues^108,109^. Here, we focused on the costs for the biosynthesis of proteins, since they by far exceed the mRNA-related costs^108^. The costs for protein synthesis and degradation in turn are dominated by those for amino acid chain elongation^108^, which linearly depend on the number of amino acids that are added to and removed from peptide chains during synthesis and degradation. The rate of amino acid addition and removal to a single protein species is given by the its protein degradation rate 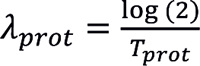 (with protein half-life *T_prot_*) and the number of amino acids per protein *N_aa_*:

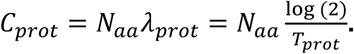

For a protein species with *N_prot_* copies this results in total costs of

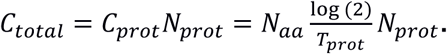

Note that the actual costs (in ATP per time) are 5 times this value^108,110,111^. We here omit this factor.

### Weighted correlation network analysis (WGCNA) for mouse models and late aging

To construct co-expression networks to find proteins with similar fold change (log2FC) differences in late aging and eight mouse models, we filtered genes with significant differences (adjusted P ≤ 0.05) in physiological aging or any mouse model in total and insoluble fractions, yielding ∼1,000 genes. We used log2FC values for late aging and 8 mouse models in the total fraction as input for identifying functional groups using WGCNA v1.7.11^43^. Modules were created within a signed network based on the topological overlap matrix (TOM), applying a soft-threshold power of 16 to approximate scale-free topology. We defined modules with a minimum size of 50 and a cutHeight of 0.993. Proteins were clustered based on dissimilarity (1-TOM, used in Extended Data Fig. 32).

To evaluate the functional significance of these modules, we conducted gene over-representation analysis using the enrichGO() function from clusterProfiler v4.5.0^97^. The background gene set included all genes in the genome (Supplementary Table 10, used in Fig. 6f). Using the modules defined in the total fraction, we examined log2FC differences in our insoluble fraction to check if there are changes in the solubility of these protein modules.

### Statistical analysis

All statistical analyses were conducted using R version 4.0.

### Structural visualization and figure preparation

Structural representations of mitochondrial complex I were generated using PyMOL (Schrödinger, LLC). Graphical icons were generated using envato (https://elements.envato.com/) and BioRender (BioRender.com). Figures were generated using R version 4.0 and subsequently edited and finalized using Adobe Illustrator (Adobe Inc.).

## Acknowledgements

We would like to acknowledge the help of the Scientific Compute Cluster (SCC) of the GWDG for the high-performance computing system. The Bioanalytical Mass Spectrometry group at the Max Planck Institute for Biophysical Chemistry, Göttingen is gratefully acknowledged for their technological support in mass spectrometry analyses. The Doppelganger Biosystem GmbH is gratefully acknowledged for their technological support in calculating the metabolic essentials. We are grateful to Alexander Peter Wegner for helping with IT to set up the BrainAging-MAP website. We are grateful to Nadine Kurz and Dr. Ali Shaib for their input on the initial draft of the BrainAging-MAP website. We are very grateful to Dr. Cristina Cheroni and Dr. Giuseppe Testa for their input on the gene overlap analysis. We are very grateful to all the members of the Fornasiero lab for the useful comments that were made after a critical reading of our original draft. We thank all members of the Fornasiero and Tchumatchenko lab for fruitful discussions.

## Author contributions

Conceptualization (E.F.F), Data Curation (N.H.K and V.K), Formal Analysis (N.H.K, V.K, M.C.M, M.T, P.G, C.B, K.M and T.T), Funding Acquisition (E.F.F), Investigation (N.H.K and VK), Methodology (V.K, E.F.F, S.V.K and H.U), Project Administration (E.F.F), Resources (A.F, T.F.O, A.C, M.H, M.P, C.S, S.M.I, and C.L.O), Software (N.H.K), Supervision (E.F.F), Validation (V.K and N.H.K), Visualization (N.H.K and E.F.F), Writing – Original Draft (N.H.K and E.F.F), Writing – Review & Editing (N.H.K and E.F.F with the input of all authors).

## Ethics Statement

All animal experiments were conducted in accordance with the guidelines and regulations of the Lower Saxony State Office for Consumer Protection and Food Safety (Niedersächsisches Landesamt für Verbraucherschutz und Lebensmittelsicherheit). The animal license for WT-aSyn is ‘20.3469’, A30P is ‘19.3213’, and for UCP2-/- is ‘V242-7224,122.5’.

## Code availability

This study used open-source software and codes, specifically R (https://www.r-project.org/), DESeq2 (https://bioconductor.org/packages/release/bioc/vignettes/DESeq2/inst/doc/DESeq2.html), DeqMS (https://bioconductor.org/packages/release/bioc/vignettes/DEqMS/inst/doc/DEqMS-package-vignette.html), WebGestalt 2024 (https://www.webgestalt.org/), clusterProfiler (https://github.com/YuLab-SMU/clusterProfiler), and WGCNA (https://cran.r-project.org/web/packages/WGCNA/index.html).

## Supplementary Figures

**Extended Data Figure 1:**
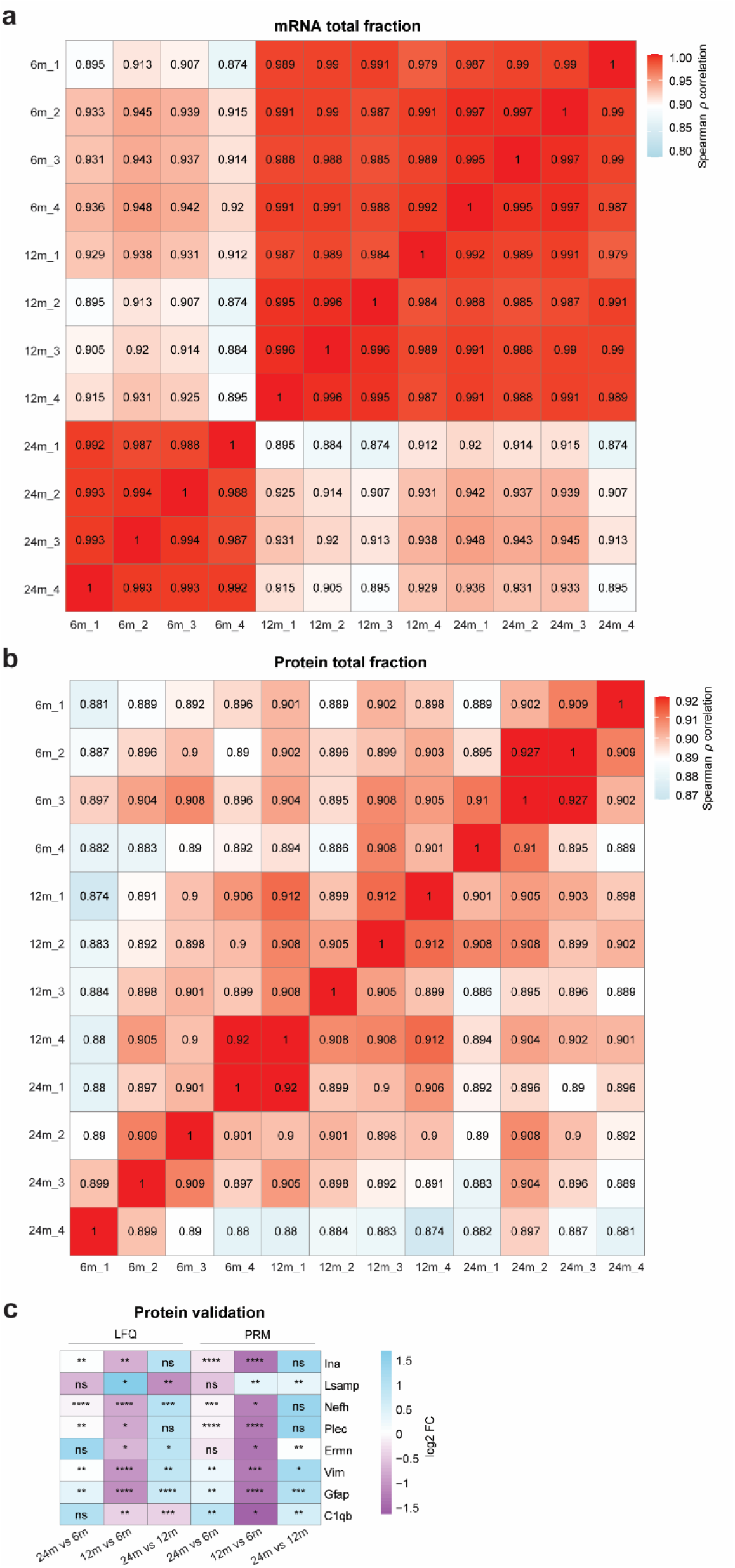
mRNA and protein correlation and validation. (a) Heatmap depicting Spearman ρ correlation across biological replicates for the mRNA-total fraction. (b) Heatmap for the protein-total fraction, with red indicating high correlation and sky blue indicating low correlation. (c) Validation of proteins using label-free quantification (LFQ) and Parallel Reaction Monitoring (PRM) analyses, where sky blue represents increased expression with aging and medium orchid denotes decreased expression with aging.

**Extended Data Figure 2:**
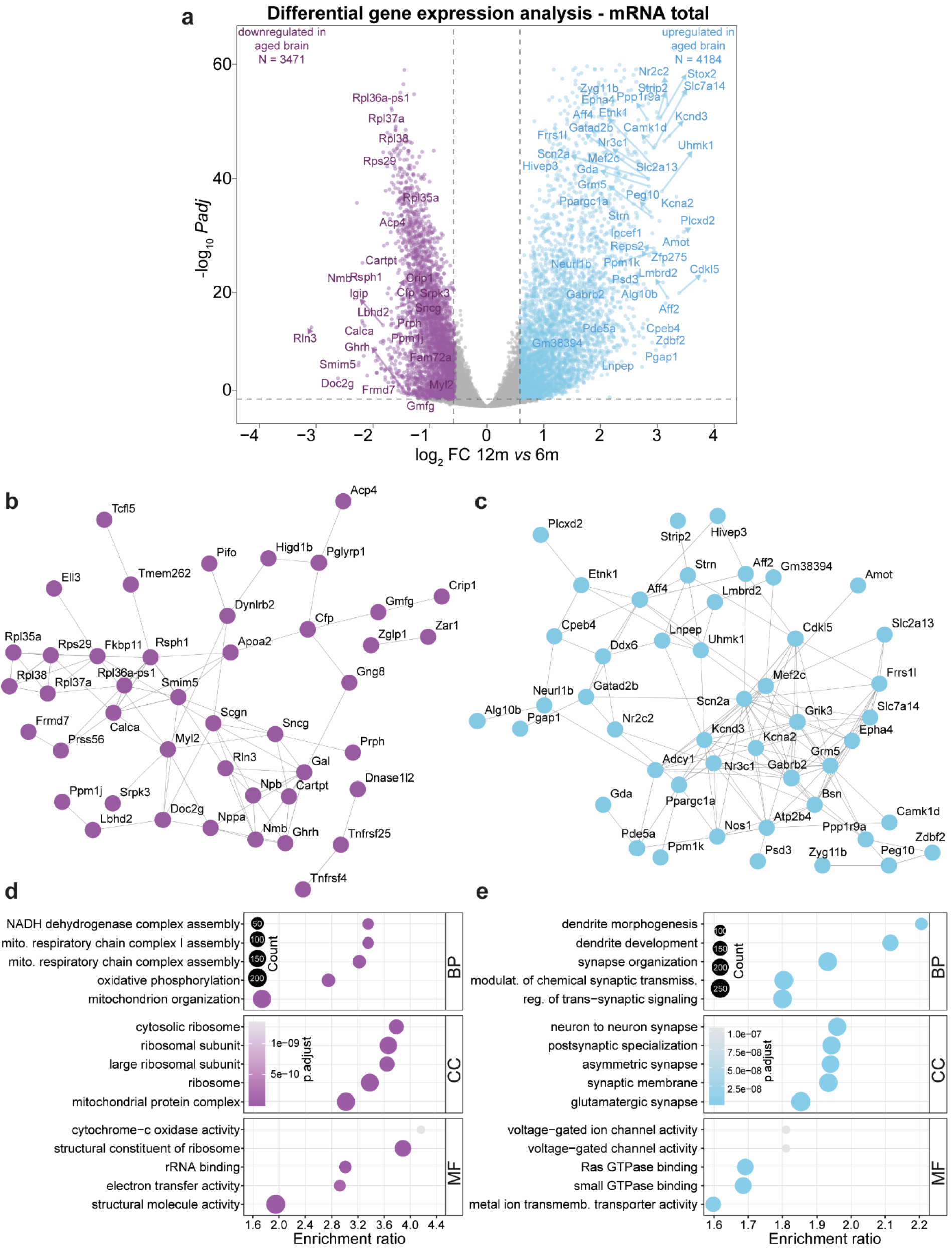
Gene expression changes in the aging mouse brain for 12m vs 6m in mRNA-total fraction. (**a**) Volcano plot displaying the differentially expressed genes for 12m vs 6m. Inset: dot plot for biological replicate variability calculated using PCA, where sky blue represents increased expression with aging and medium orchid denotes decreased expression with aging. (**b, c**) STRING representation for the top 50 most (**b**) down- or (**c**)upregulated genes in 12m brain (respectively medium orchid and light blue). (**d, e**) GO-ORA of genes that are significantly down- or upregulated in 12m (*padj* ≤ 0.05, |log_2_FC| ≥ 0.58.

**Extended Data Figure 3:**
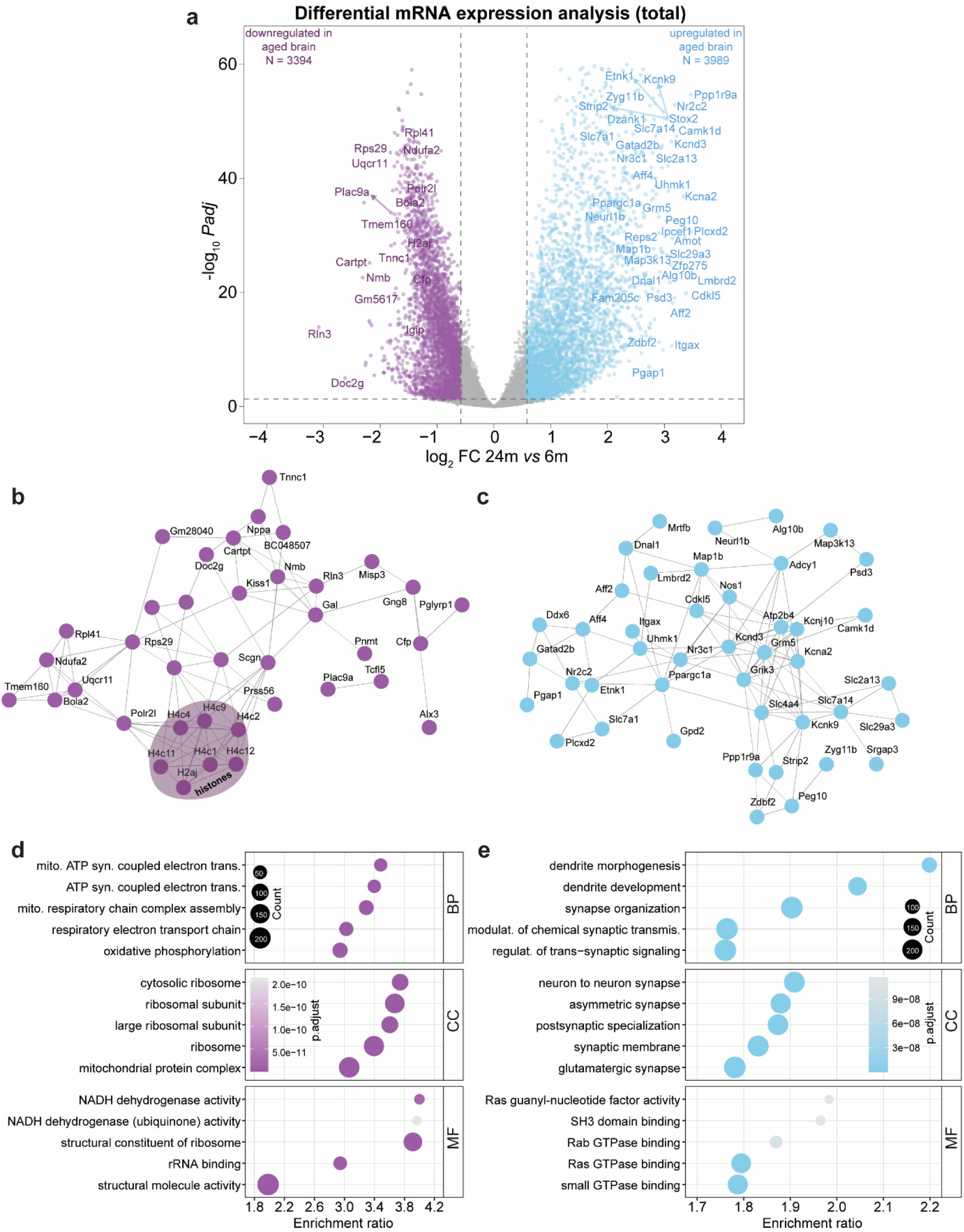
Gene expression changes in the aging mouse brain for 24m vs 6m in mRNA-total fraction. (**a**) Volcano plot displaying the differentially expressed genes for 24m vs 6m. Inset: dot plot for biological replicate variability calculated using PCA, where sky blue represents increased expression with aging and medium orchid denotes decreased expression with aging. (**b, c**) STRING representation for the top 50 most (**b**) down- or (**c**)upregulated genes in 24m brain (respectively medium orchid and light blue). (**d, e**) GO-ORA of genes that are significantly down- or upregulated in 24m (*padj* ≤ 0.05, |log_2_FC| ≥ 0.58).

**Extended Data Figure 4:**
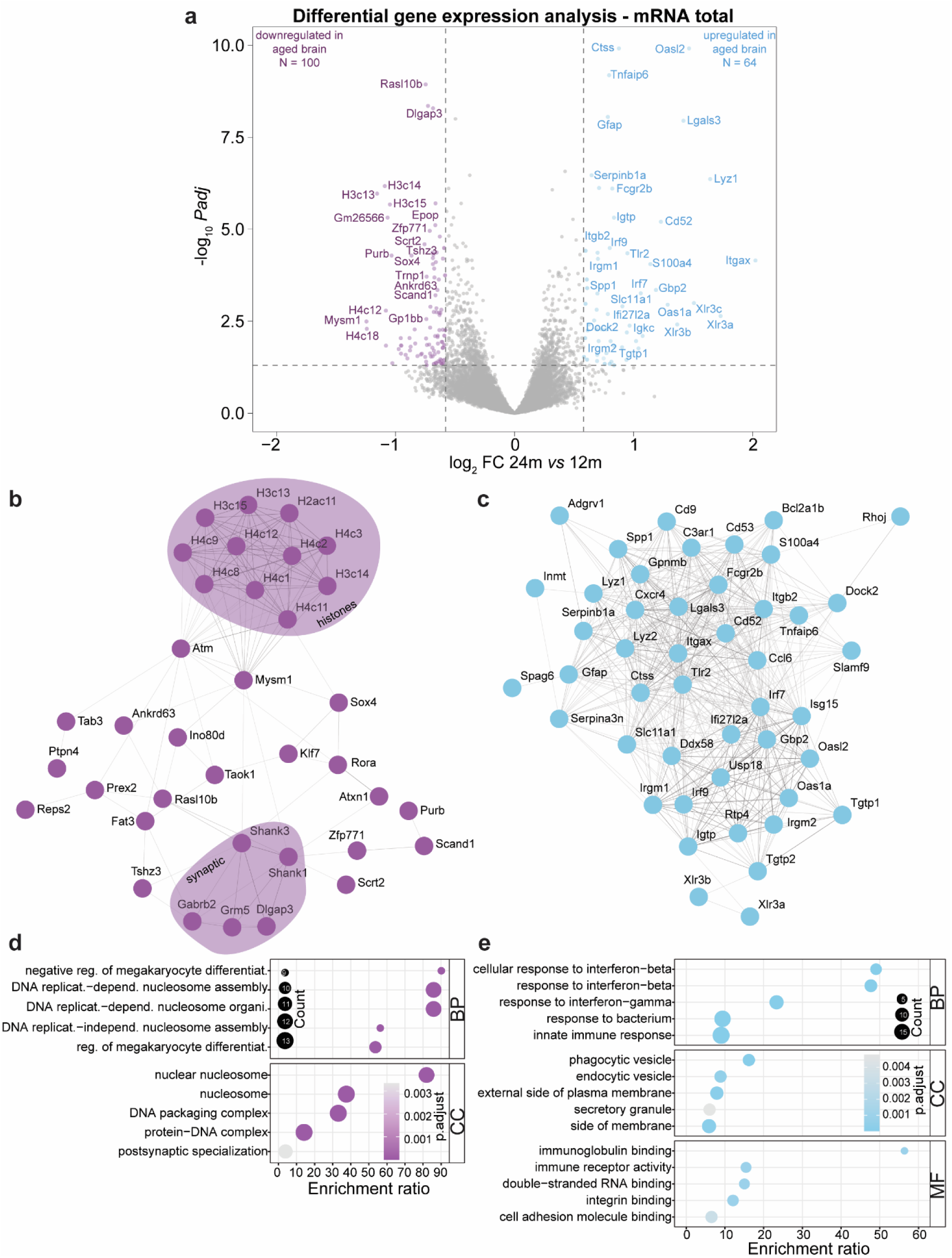
Gene expression changes in the aging mouse brain for 24m vs 12m in mRNA-total fraction. (**a**) Volcano plot displaying the differentially expressed genes for 24m vs 12m. Inset: dot plot for biological replicate variability calculated using PCA, where sky blue represents increased expression with aging and medium orchid denotes decreased expression with aging. (**b, c**) STRING representation for the top 50 most (**b**) down- or (**c**)upregulated genes in 24m brain (respectively medium orchid and light blue). (**d, e**) GO-ORA of genes that are significantly down- or upregulated in 24m (*padj* ≤ 0.05, |log_2_FC| ≥ 0.58.

**Extended Data Figure 5:**
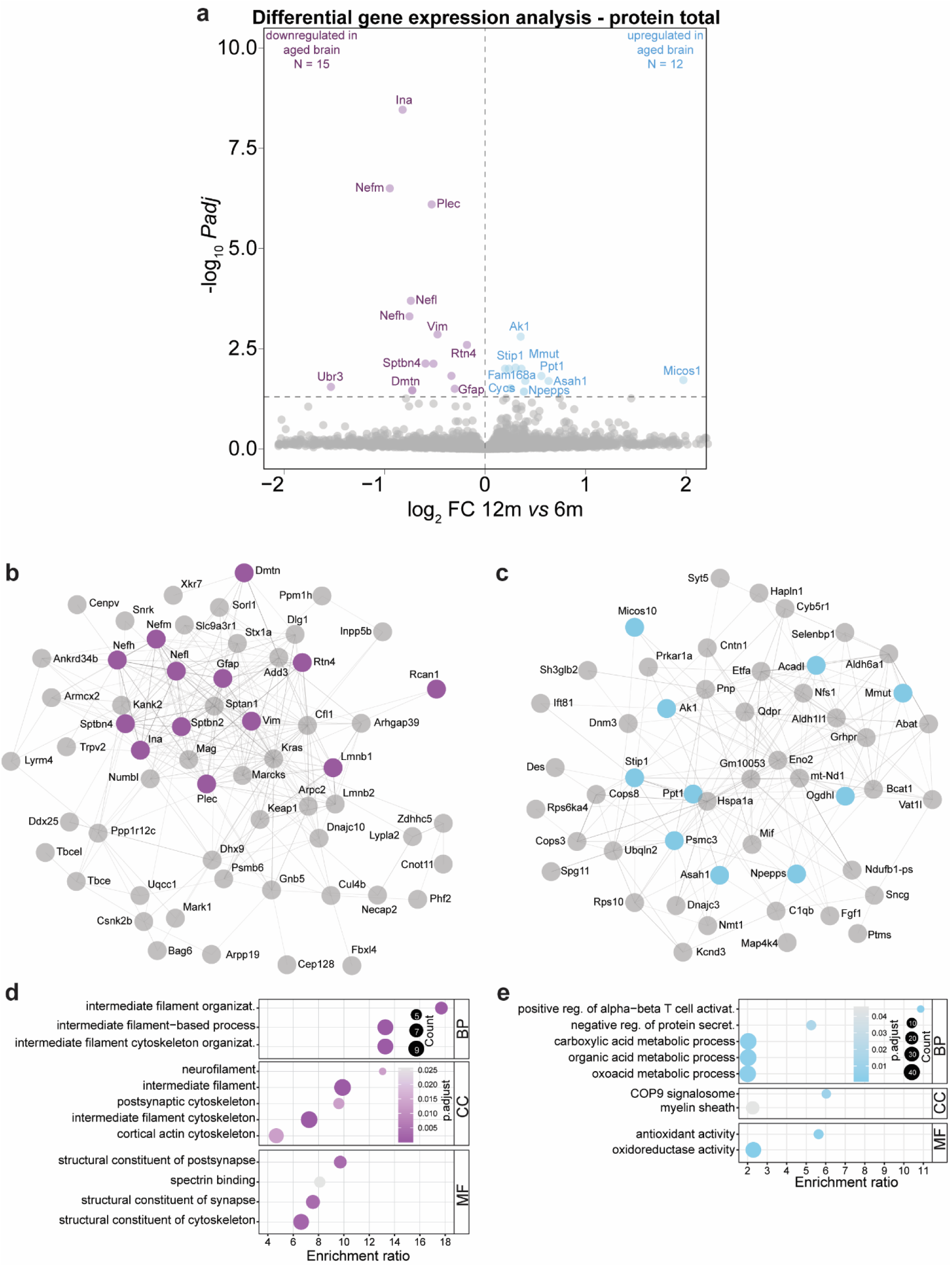
Protein expression changes in the aging mouse brain for 12m vs 6m in protein-total fraction. (a) Volcano plot displaying the differentially expressed proteins for 12m vs 6m. Inset: dot plot for biological replicate variability calculated using PCA, where sky blue represents increased expression with aging and medium orchid denotes decreased expression with aging. (b, c) STRING representation for the top 50 most (b) down- or (c)upregulated proteins in 12m brain (respectively medium orchid and light blue), grey are the proteins that are not significant. (d, e) GO-ORA of the top150 proteins that are down- or upregulated in 12m.

**Extended Data Figure 6:**
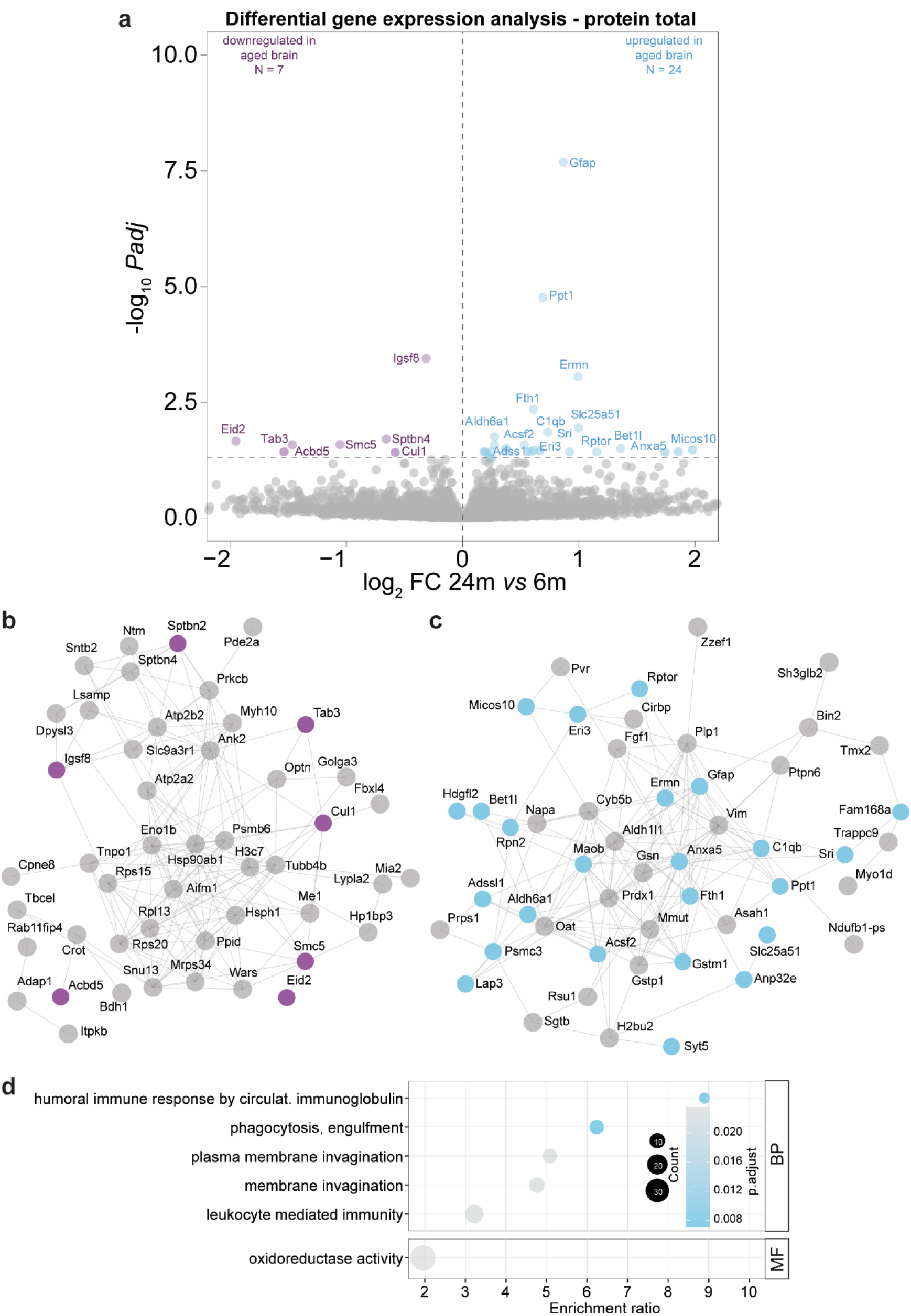
Protein expression changes in the aging mouse brain for 24m vs 6m in protein-total fraction. (a) Volcano plot displaying the differentially expressed proteins for 24m vs 6m. Inset: dot plot for biological replicate variability calculated using PCA, where sky blue represents increased expression with aging and medium orchid denotes decreased expression with aging. (b, c) STRING representation for the top 50 most (b) down- or (c)upregulated proteins in 24m brain (respectively medium orchid and light blue), grey are the proteins that are not significant. (d, e) GO-ORA of the top150 proteins that are down- or upregulated in 24m.

**Extended Data Figure 7:**
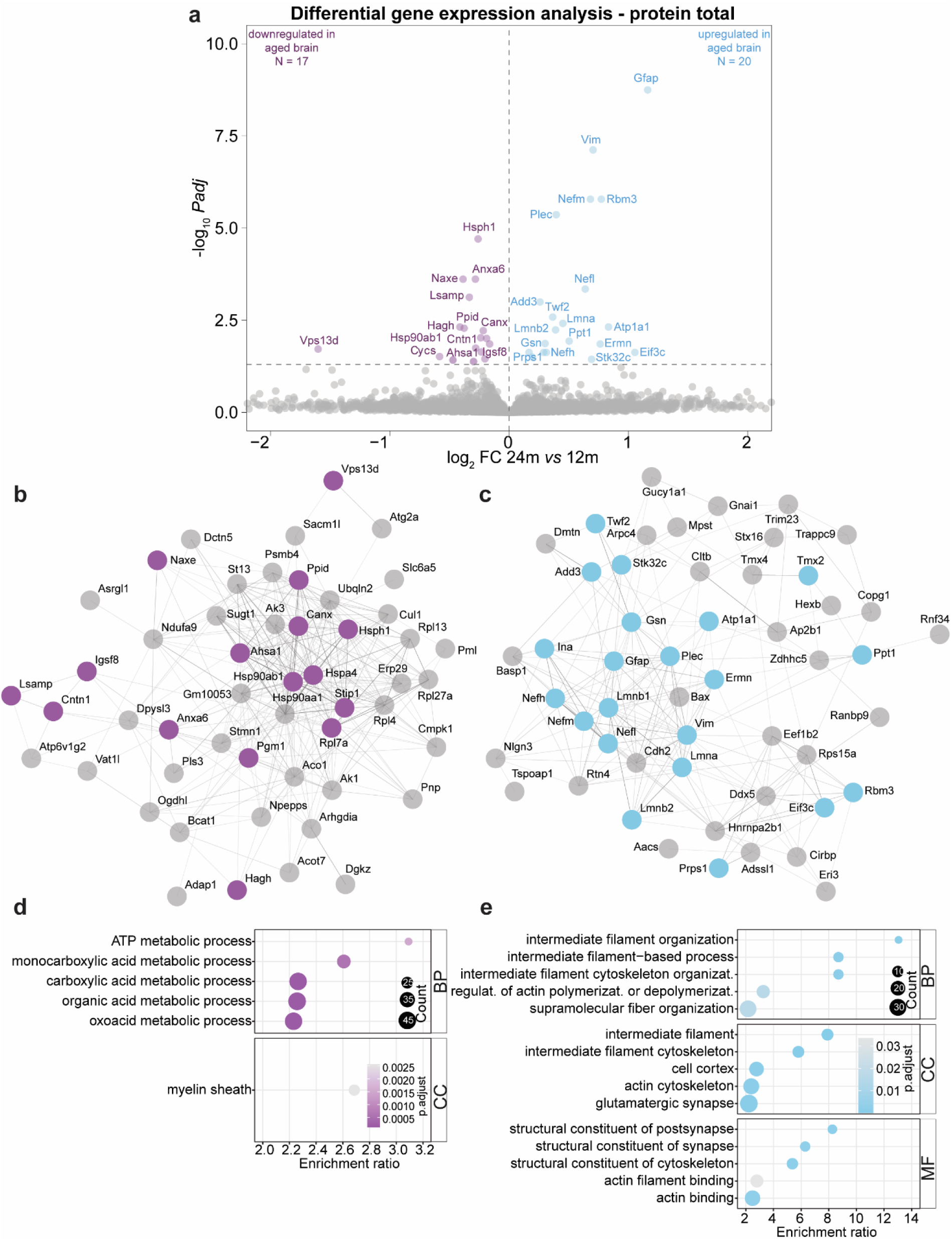
Protein expression changes in the aging mouse brain for 24m vs 12m in protein-total fraction. (a) Volcano plot displaying the differentially expressed proteins for 24m vs 12m. Inset: dot plot for biological replicate variability calculated using PCA, where sky blue represents increased expression with aging and medium orchid denotes decreased expression with aging. (b, c) STRING representation for the top 50 most (b) down- or (c)upregulated proteins in 24m brain (respectively medium orchid and light blue), grey are the proteins that are not significant. (d, e) GO-ORA of the top150 proteins that are down- or upregulated in 24m.

**Extended Data Figure 8:**
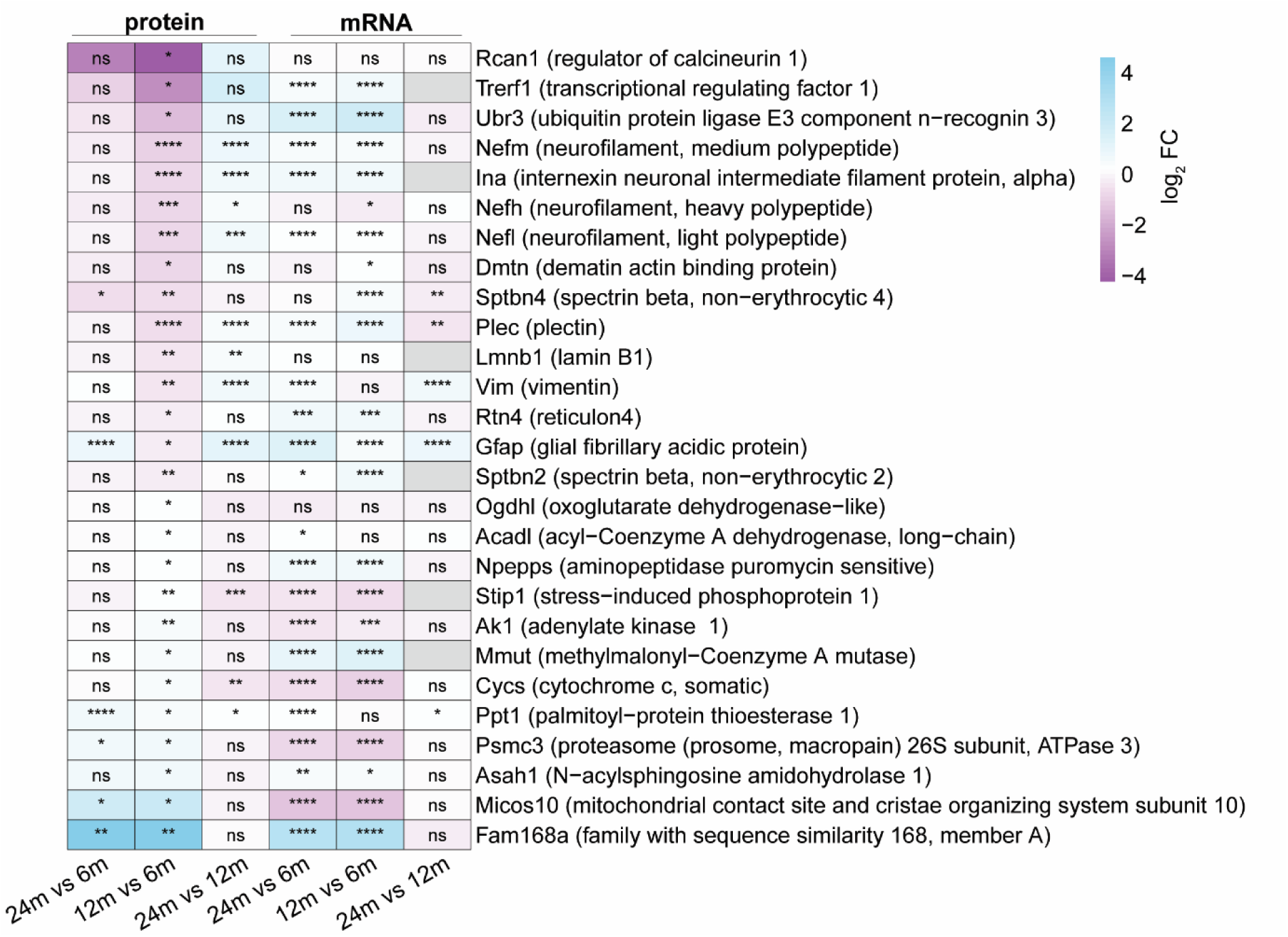
Heatmap of proteins significantly altered in the 12m vs. 6m comparison for the protein- and mRNA-total Fraction. The heatmap displays proteins that show significant changes between 12 months and 6 months in the protein-total fraction. Sky blue indicates proteins with increased expression during aging, while medium orchid denotes proteins with decreased expression during aging.

**Extended Data Figure 9:**
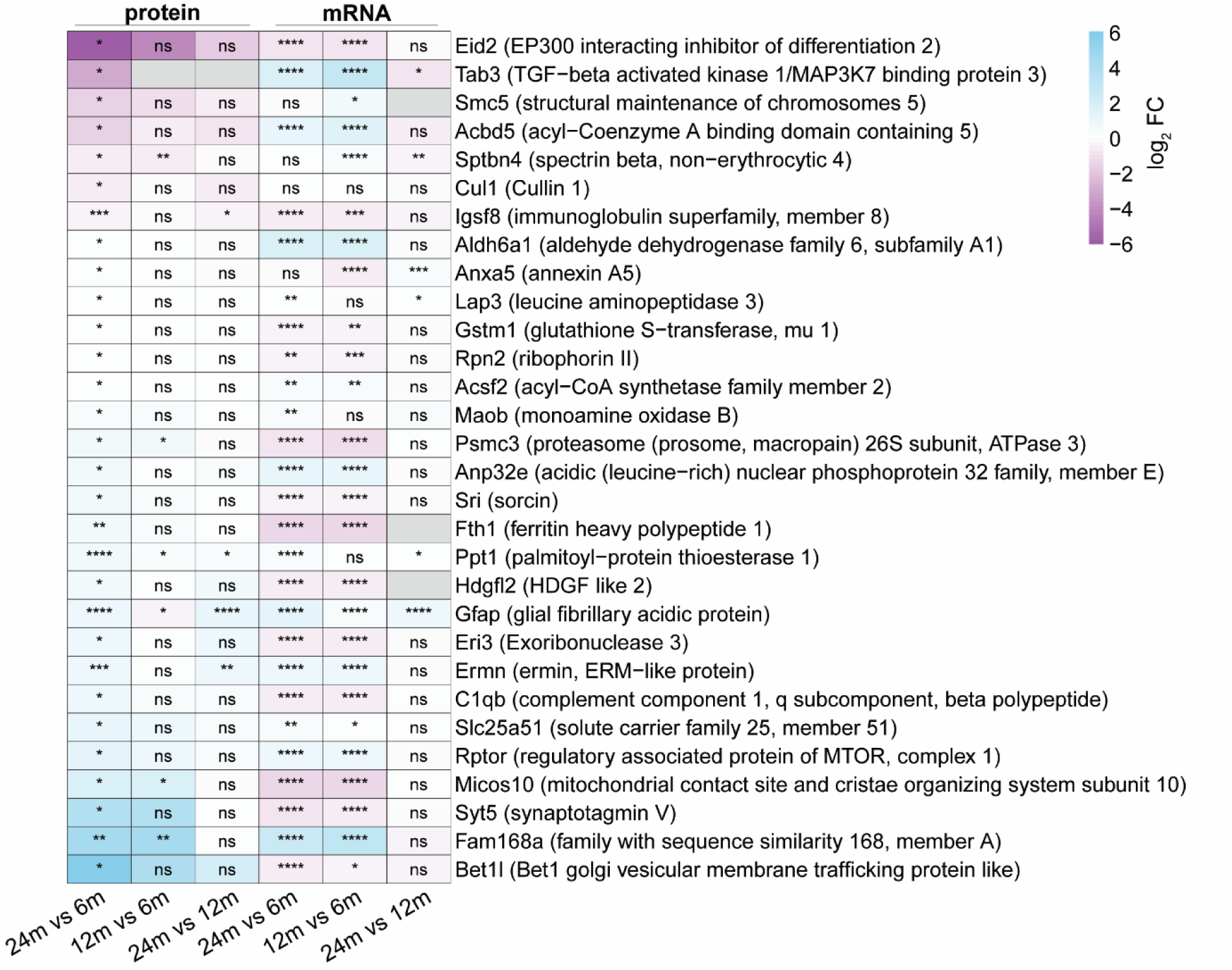
Heatmap of proteins significantly altered in the 24m vs. 6m comparison for the protein- and mRNA-total Fraction. The heatmap displays proteins that show significant changes between 24 months and 6 months in the protein-total fraction. Sky blue indicates proteins with increased expression during aging, while medium orchid denotes proteins with decreased expression during aging.

**Extended Data Figure 10:**
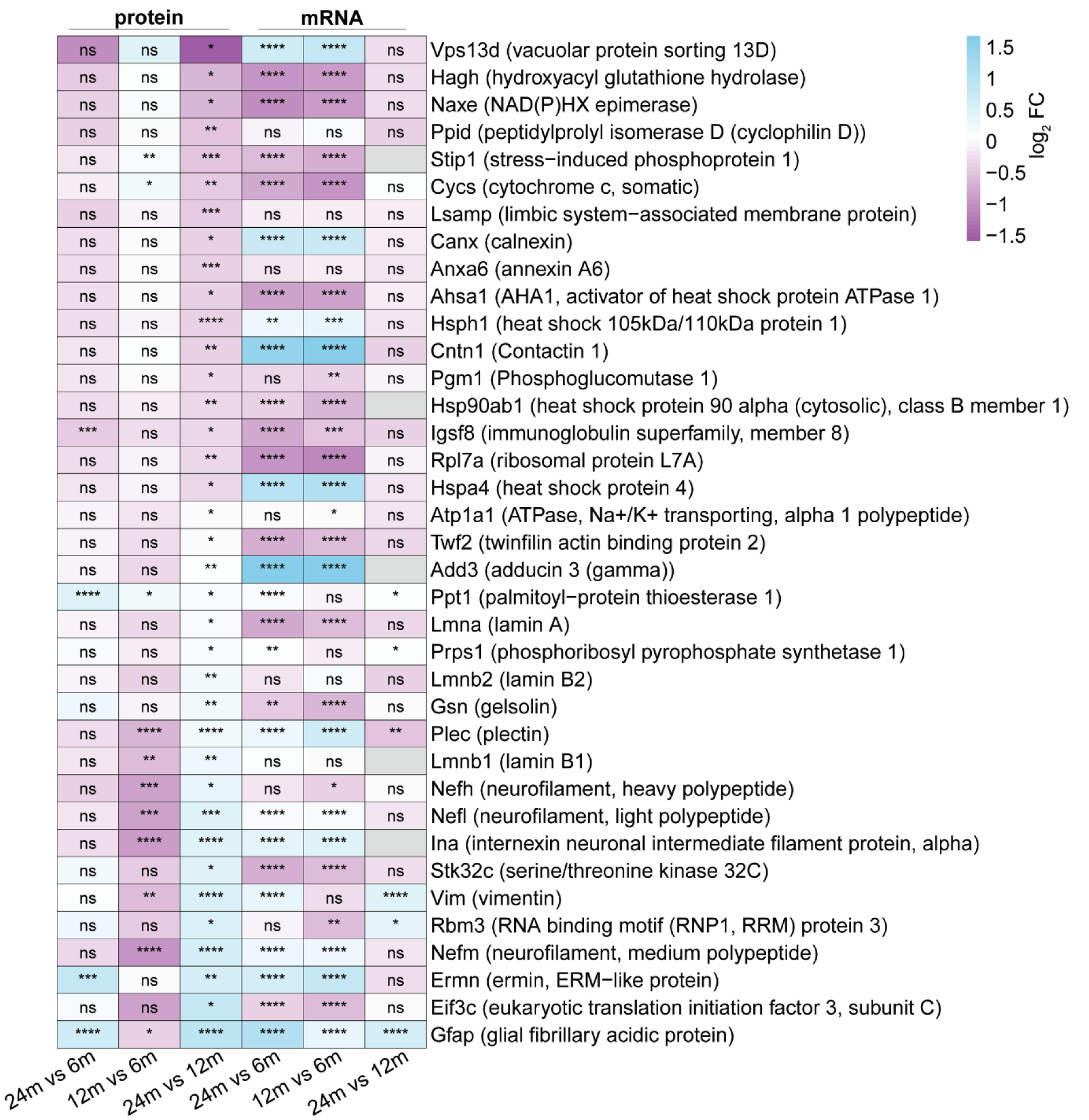
Heatmap of proteins significantly altered in the 24m vs. 12m comparison for the protein- and mRNA-total Fraction. The heatmap displays proteins that show significant changes between 24 months and 12 months in the protein-total fraction. Sky blue indicates proteins with increased expression during aging, while medium orchid denotes proteins with decreased expression during aging.

**Extended Data Figure 11:**
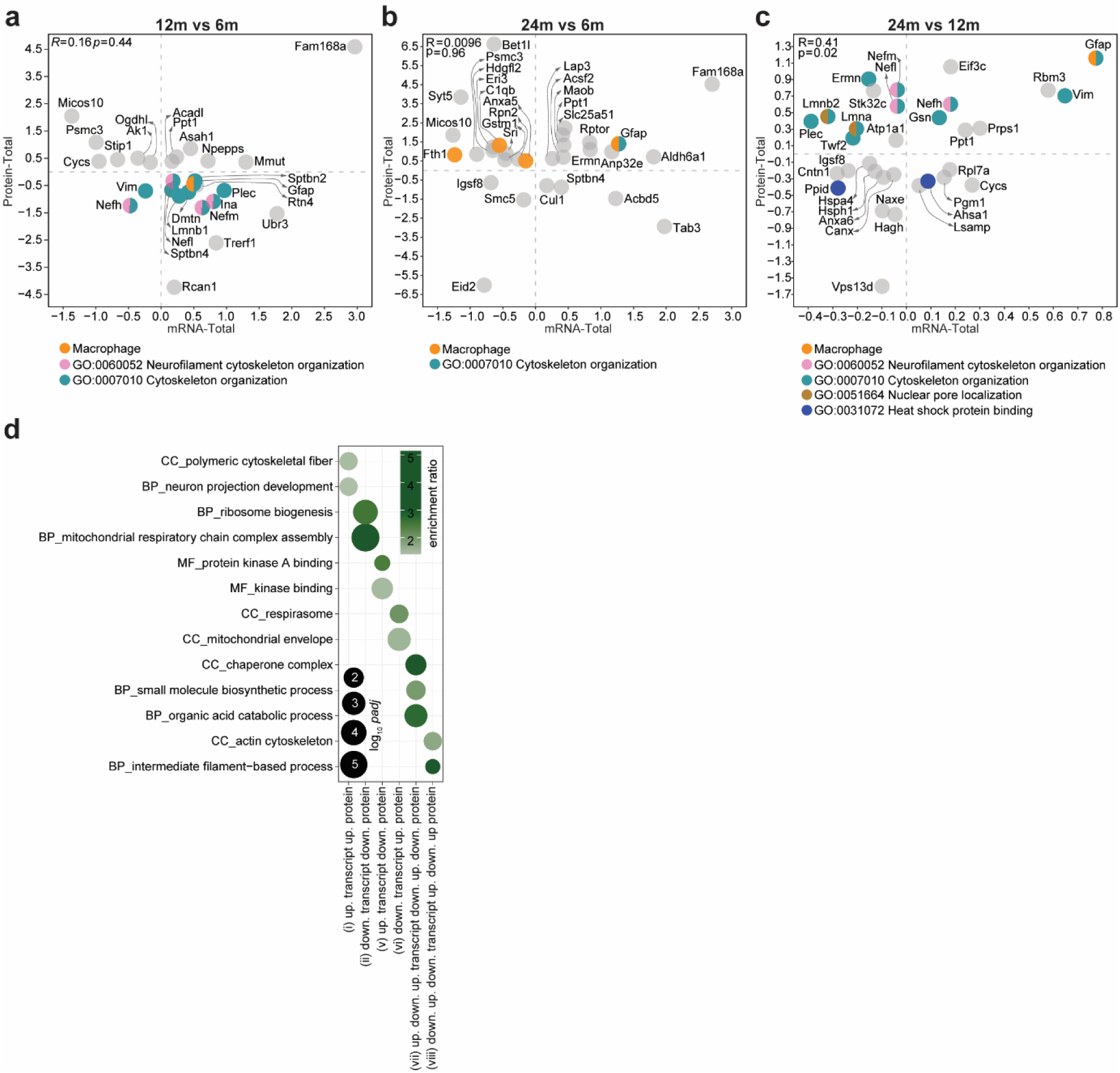
Comparison of the protein- and mRNA-total Fraction. (a-c) Scatter plot between mRNA and protein for the proteins that show significant changes between (a) 12m and 6m (b) 24m and 6m (c) 24m and 12m in the protein-total fraction. (d) GO-ORA of the non-linear dynamic patterns, here are top ∼2 GO terms based on the enrichment ratio. For all patterns in detail refer to Supplementary Table 1. Dot sizes in the enrichment graphs correspond to the padj and color scales represents the enrichment ratio for each GO term, dark green – highly enriched and gray – less enriched.

**Extended Data Figure 12:**
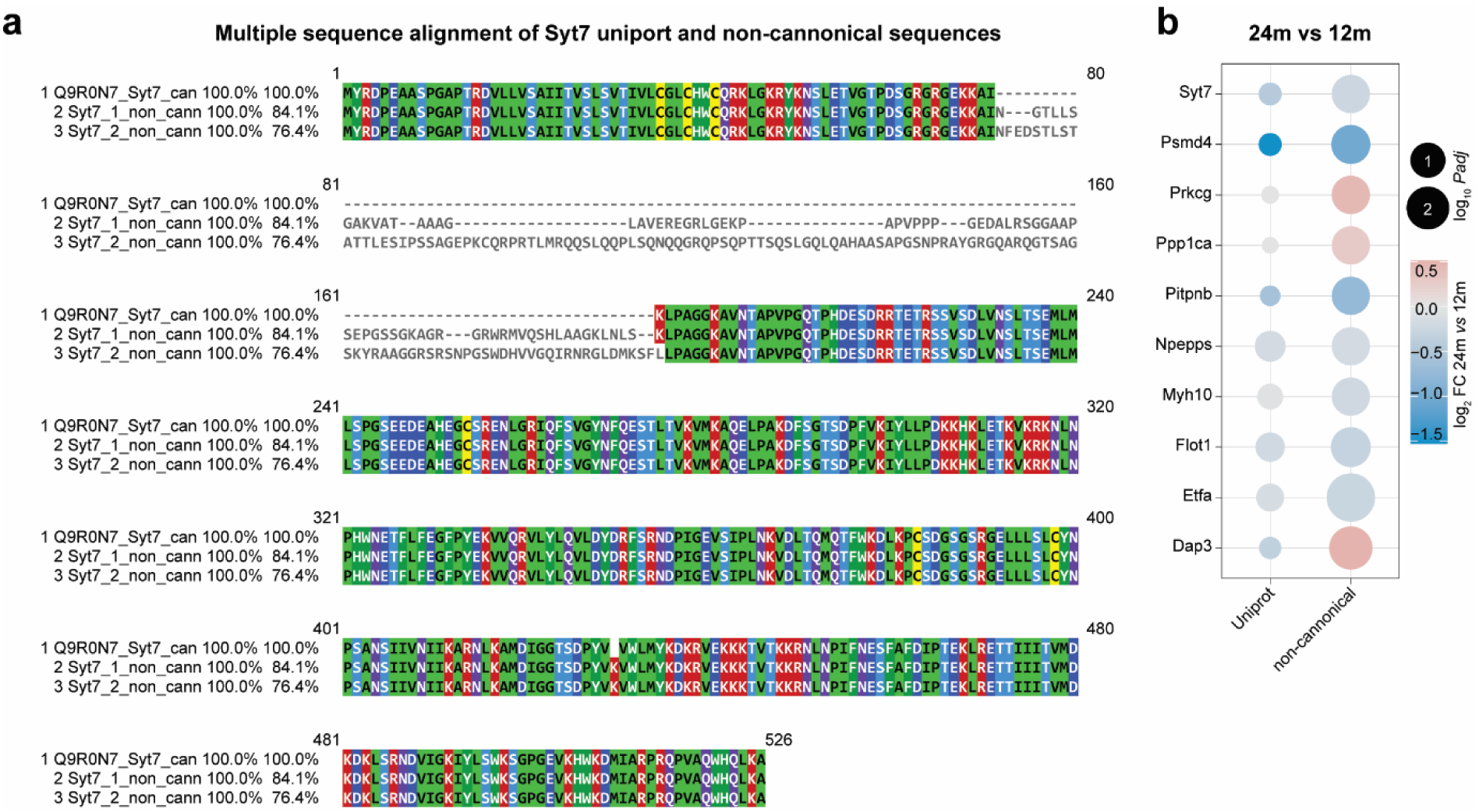
Integration of proteo-transcriptomic datasets leads to the identification of novel proteoforms with aging. (**a**) Multiple sequence alignment of Syt7 canonical (can. the uniport sequence, Q9R0N7) and 2 non-canonical (non_cann.) proteoforms (Syt7_1, Syt7_2). (**b**) Dot plot for proteoforms that are differentially expressed at 24m vs. 12m (here the Syt7_1 is shown for the non-canonical as this showed significant difference). Dot sizes correspond to the absolute log_10_padj and color scales represents the log_2_FC for protein, red – upregulated and blue – downregulated at 24m compared to 12m.

**Extended Data Figure 13:**
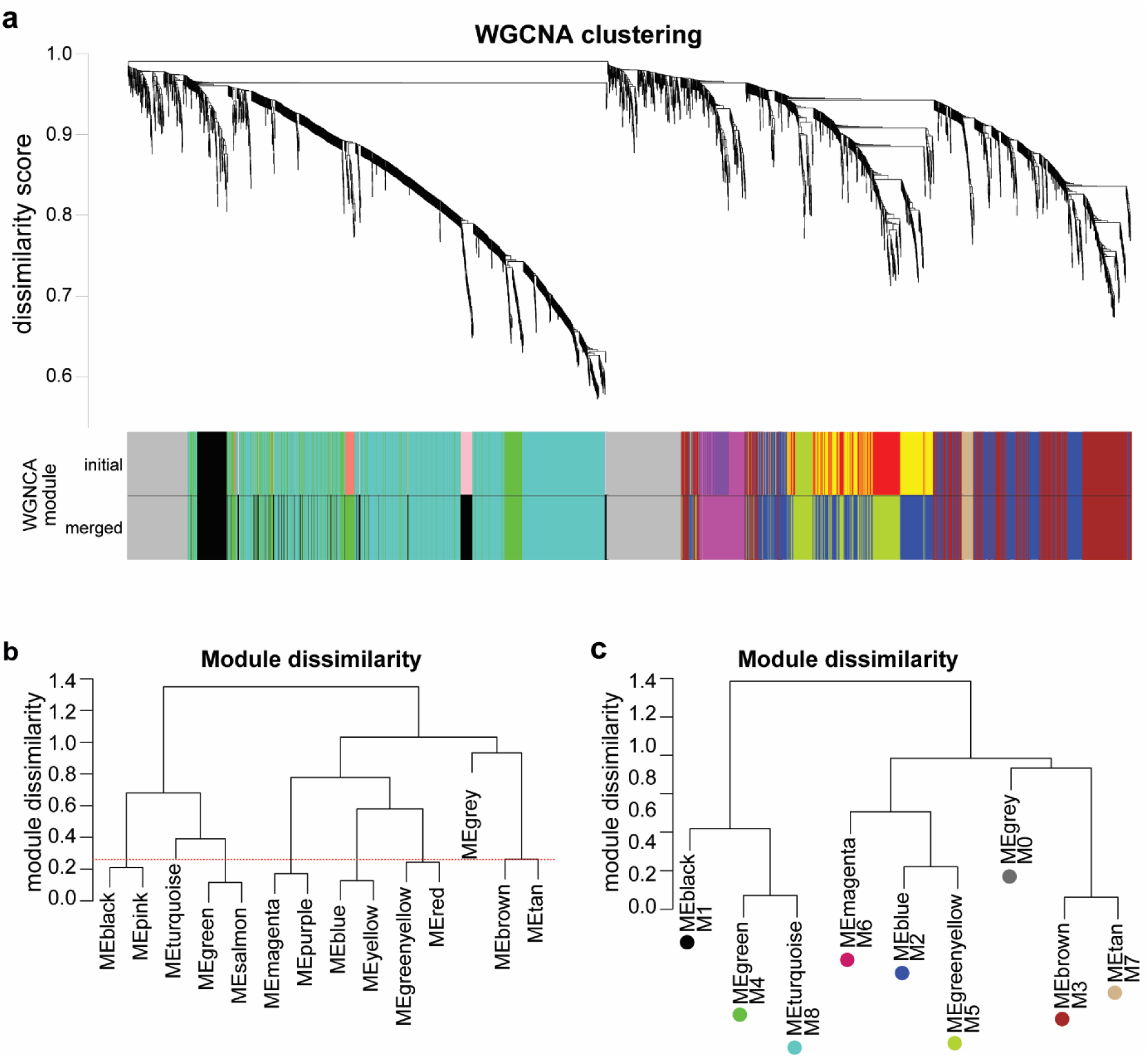
Co-expression modules WO merging for p/m coefficient in the aging mouse brain. (**a**) WGNCA dendrogram with highlighted modules (lower colored bar). Genes were clustered based on a dissimilarity measure. The branches are modules of closely correlated gene groups that have a similar p/m coefficient. (**b**) Module dissimilarity based on module eigengene distances WO merge. Modules are grouped based on their p/m coefficient. (**c**) Module dissimilarity based on module eigengene distances after merging. Modules are grouped based on their p/m coefficient.

**Extended Data Figure 14:**
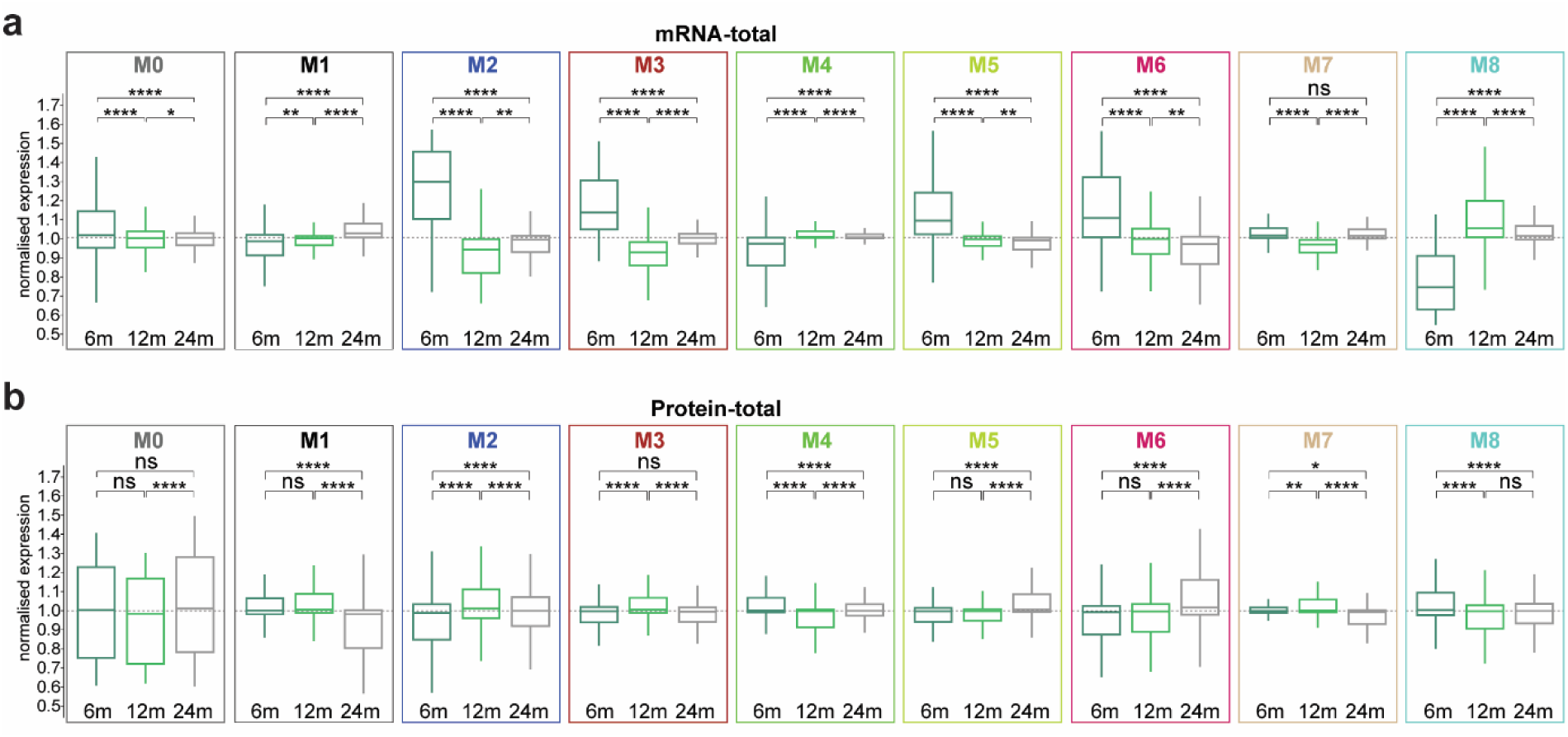
mRNA and protein level changes in the co-expression modules across ages. **(a, b)** Boxplots of normalized (**a**) mRNA-total (**b**) protein-total in 6m, 12m and 24m mouse brain, grouped by their modules detected in the WGNCA method. Tukey posthoc test P-value * ≤ 0.05, ** ≤ 0.01, *** ≤ 0.001 and **** ≤ 0.0001.

**Extended Data Figure 15:**
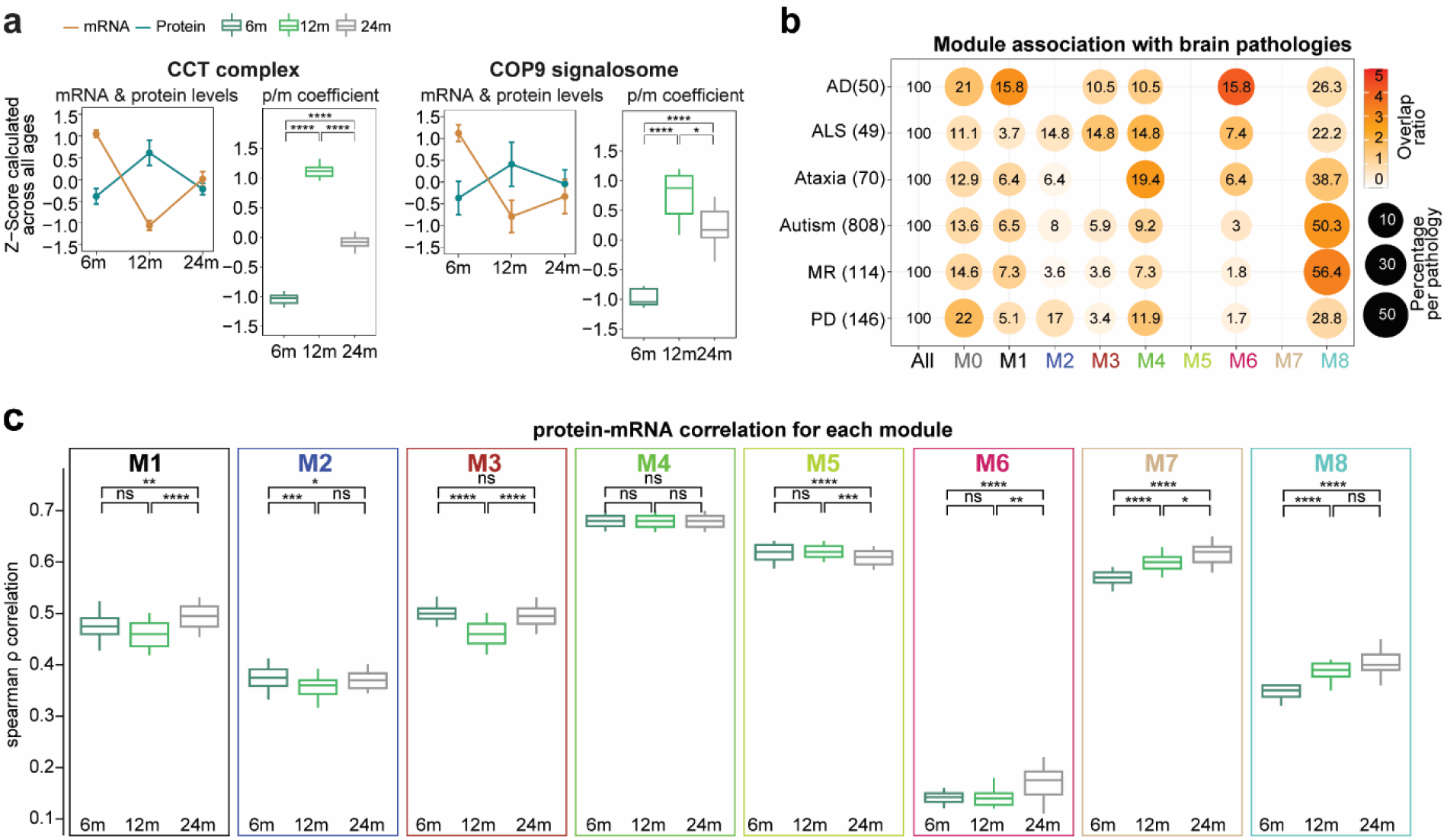
mRNA, protein and p-m coefficient change in the aging brain and its association to brain pathologies. (a) Line plots for mRNA and protein trajectories and boxplot for p/m coefficient for mitochondrial complexes. Line colors reflect R color codes, with “peru” representing mRNA and “turquoise4” representing protein. P-values indicate the results of paired t-test followed by Tukey posthoc test P-value * ≤ 0.05, ** ≤ 0.01, *** ≤ 0.001 and **** ≤ 0.0001. (b) Dot plot for brain pathologies. Dot sizes correspond to the percentage of genes per pathology and color scales represents the overlap ratio for each pathology, dark orange (higher overlap) and white (lower overlap). (c) Boxplots of spearman ρ correlation for each age group between protein and mRNA within each module. P-values indicate the results of paired t-test followed by Tukey posthoc test P-value *≤ 0.05, ** ≤ 0.01, *** ≤ 0.001 and **** ≤ 0.0001.

**Extended Data Figure 16:**
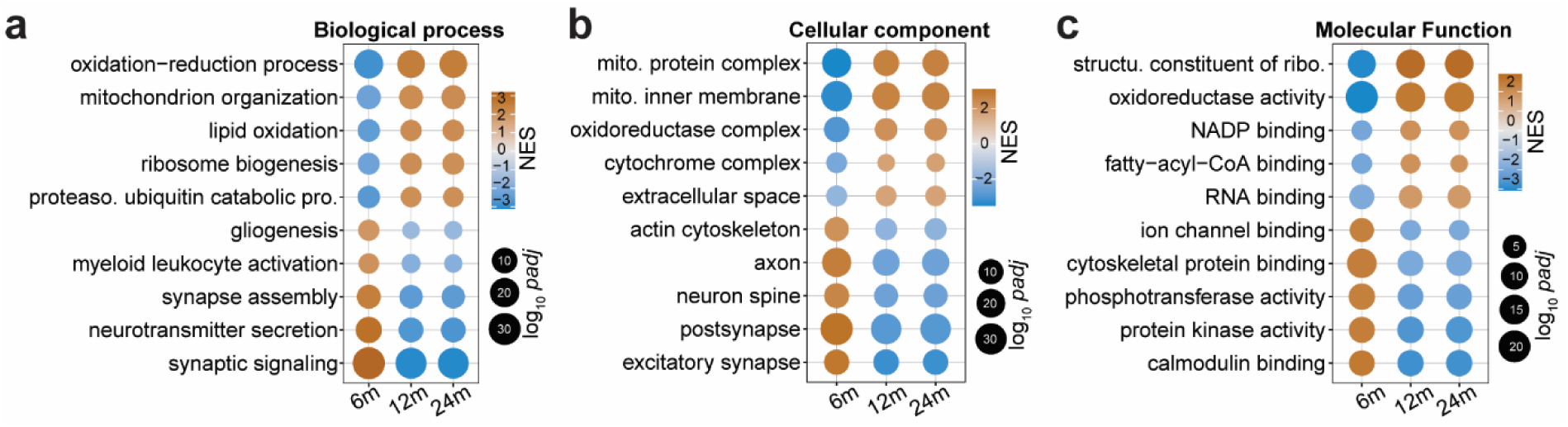
p/m coefficient changes in each age group. **(a-c)** Gene ontology (GO)-Gene set enrichment analysis (GSEA) of genes that have an increased or decreased p/m coefficient for each age. Positive coefficient values (goldenrod) indicate increased protein levels relative to mRNA, suggesting efficient translation; negative values (skyblue) indicate reduced protein levels relative to mRNA, indicating inefficient translation. The size of the dots corresponds to log_10_*padj*, bigger the size of the bubble means they are highly significant (**a**) Biological process, (**b**) Cellular component and (**c**) Molecular function.

**Extended Data Figure 17:**
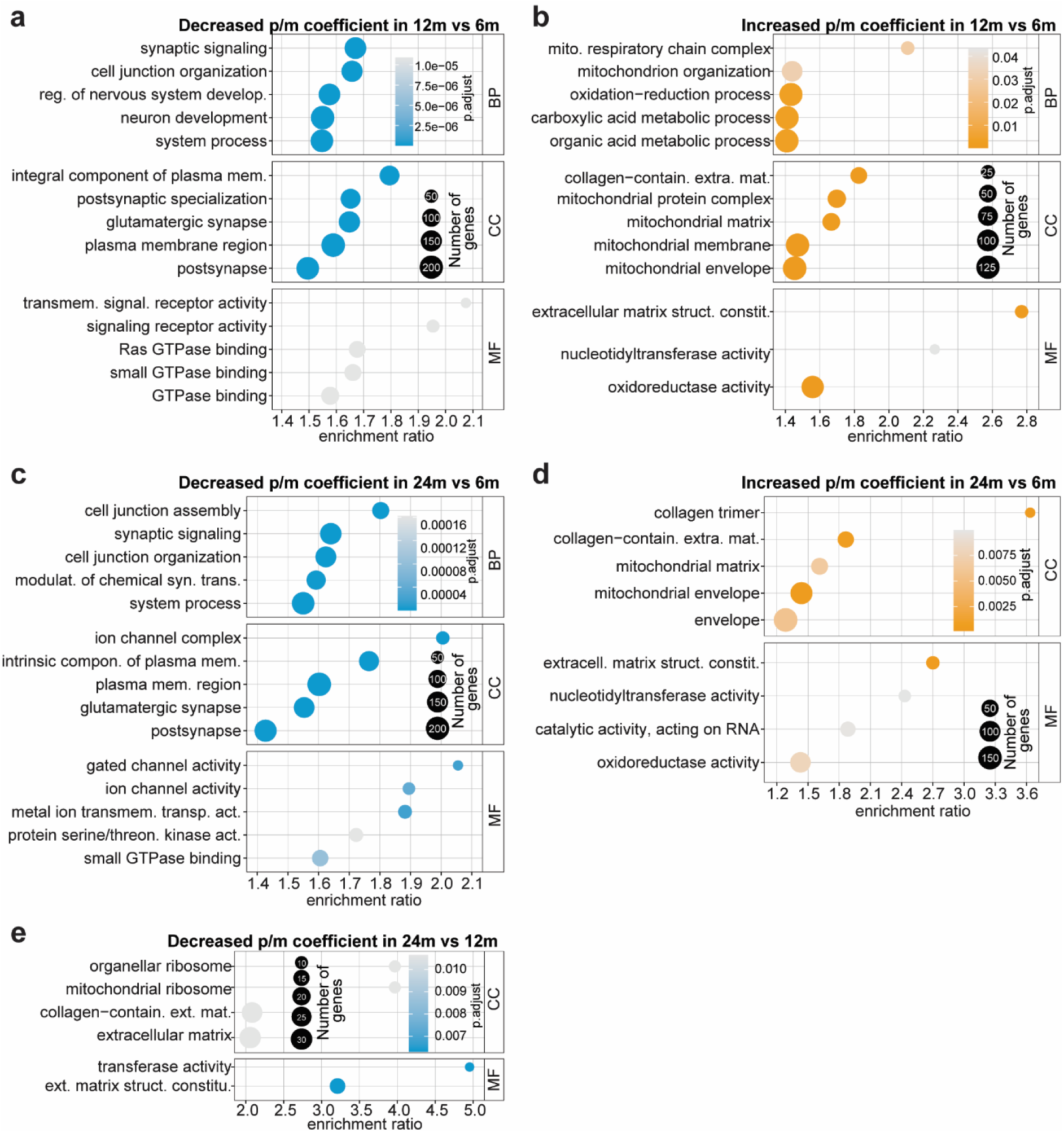
GO-ORA of genes that have an increased or decreased p/m coefficient for each age comparison. (**a)** Decreased p/m coefficient 12m vs 6m (**b**) Increased p/m coefficient 12m vs 6m (**c)** Decreased p/m coefficient 24m vs 6m (**d**) Increased p/m coefficient 24m vs 6m (**e)** Decreased p/m coefficient 24m vs 12m. Colour scale corresponds to log_10_*padj*, Goldenrod color indicate increased p/m coefficient (protein levels increased relative to mRNA), suggesting efficient translation; Skyblue color indicate decreased p/m coefficient (reduced protein levels relative to mRNA), indicating inefficient translation. Gray is less significant. The size of the dots corresponds to the number of genes, bigger the size of the bubble means they are highly significant.

**Extended Data Figure 18:**
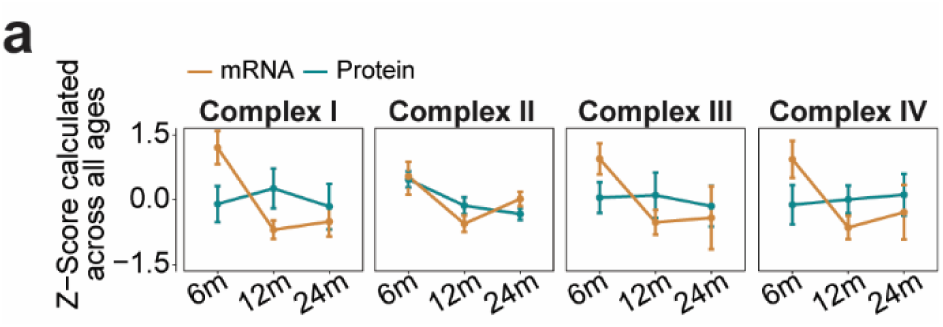
mRNA and protein levels in mitochondrial complexes. (a) Line plots for mRNA and protein trajectories for mitochondrial complexes. Line colors reflect R color codes, with “peru” representing mRNA and “turquoise4” representing protein.

**Extended Data Figure 19:**
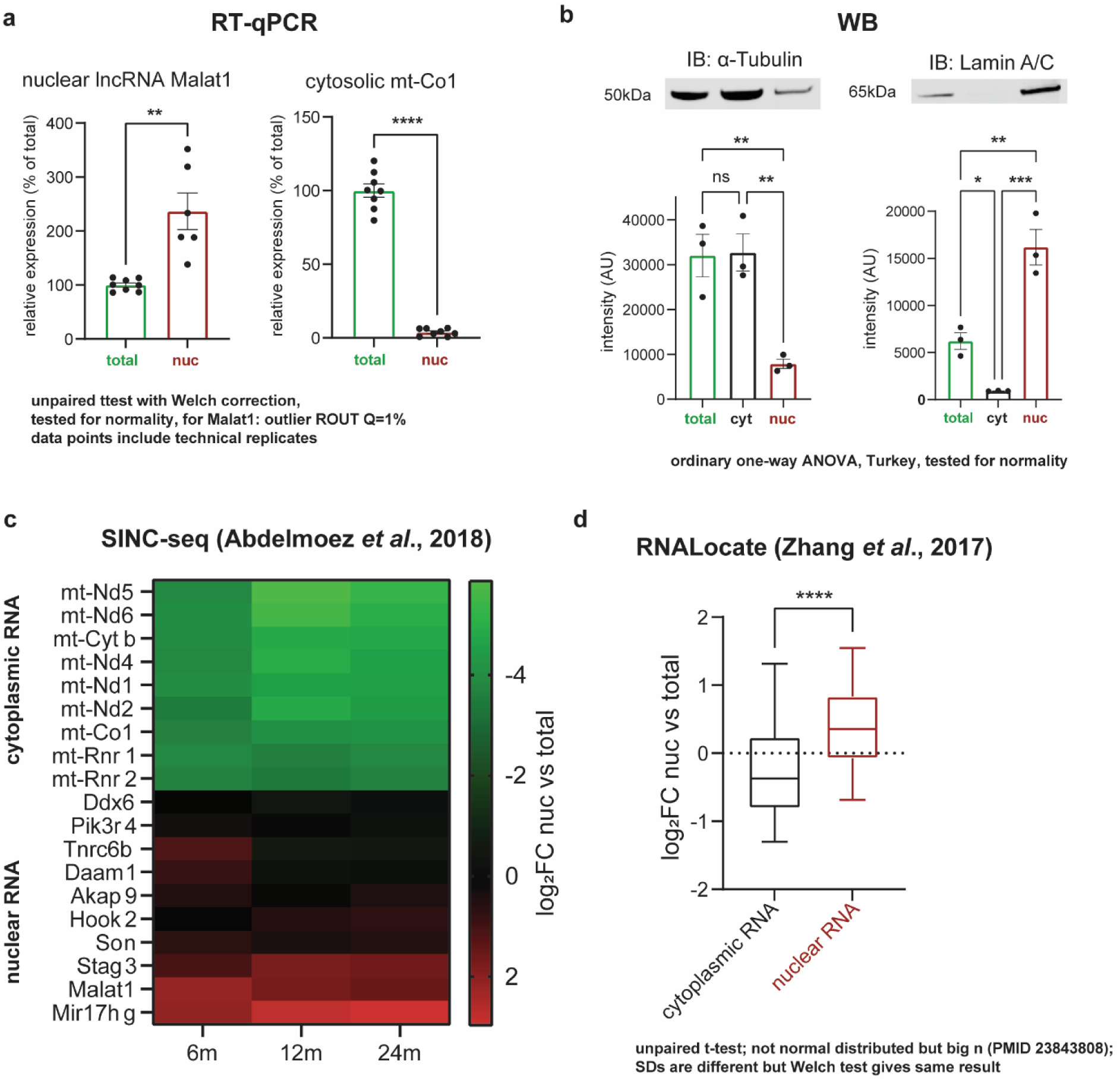
Quality control of subcellular fractionation for RNA-Isolation and subsequent sequencing. (a) Reverse-transcription qualitative PCR (RT-qPCR) confirms enrichment of nuclear transcript *Malat1* in the nuclear fraction (nuc). Reversely, cytosolic *mt-Co1* is strongly depleted in the nuclear fraction. Unpaired t-test with Welch correction; n=6 technical replicates from 3 biological samples. (b) Western blot analysis of cytosolic protein α-Tubulin, which is depleted in the nuclear fraction (nuc), and nuclear lamin Lamin A/C, which is depleted in the cytosolic fraction (Cyt) and relatively enriched in Nuc. Ordinary one-way ANOVA with Turkey correction; n=3 biological replicates. (c) Comparison of nuclear vs total enrichment (log_2_FC Nuc vs Total) of our data with single-cell integrated nuclear RNA and cytoplasmic RNA sequencing (SINC-seq^53^) showing that transcripts identified by SINC-seq as nuclear are indeed enriched in the nucleus in my dataset and cytoplasmic transcripts are de-enriched. (d) Comparison with annotations from RNALocate, a web-accessible database providing subcellular RNA localization^54^. Fold changes of nuclear enrichment (log_2_FC Nuc vs Total) are significantly higher, meaning nuclear enrichment, among transcripts that are annotated as ‘nuclear RNA’. Unpaired t-test with Welch correction of 1465 and 2259 transcripts with cytoplasmic and nuclear annotation, respectively. Boxplots show median, 25^th^ to 75^th^ percentile as box and 5^th^ to 95^th^ percentile as whiskers. * *p*<0.5, ** *p*<0.01, *** *p*<0.001, **** *p*<0.0001.

**Extended Data Figure 20:**
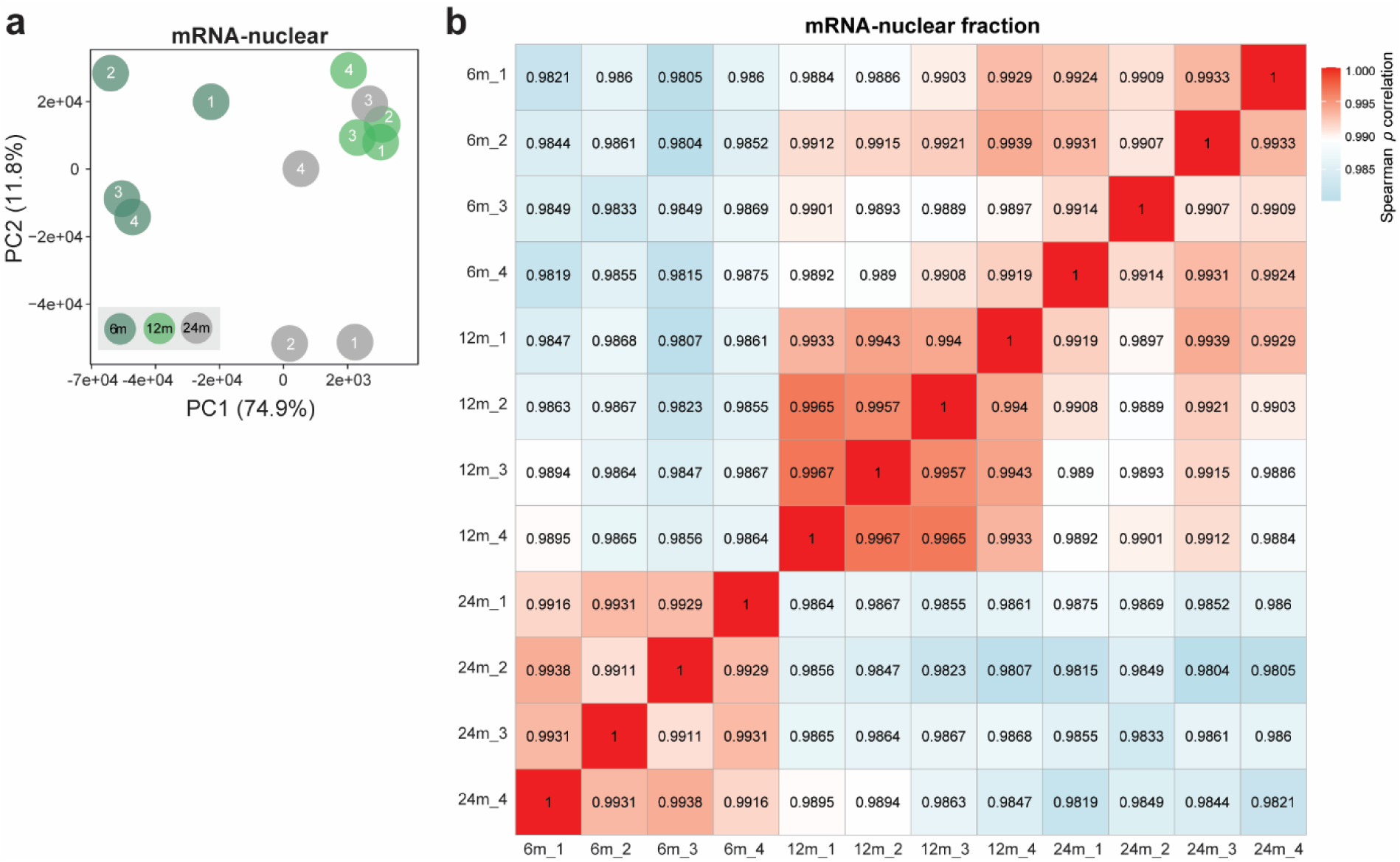
nuclear-mRNA fraction correlation and principal component analysis. **(a)** Dot plots depict biological replicate variability via principal component analysis (PCA) for the mRNA – nuclear dataset. **(b)** Heatmap for the mRNA-nuclear fraction, with red indicating high correlation and sky blue indicating low correlation.

**Extended Data Figure 21:**
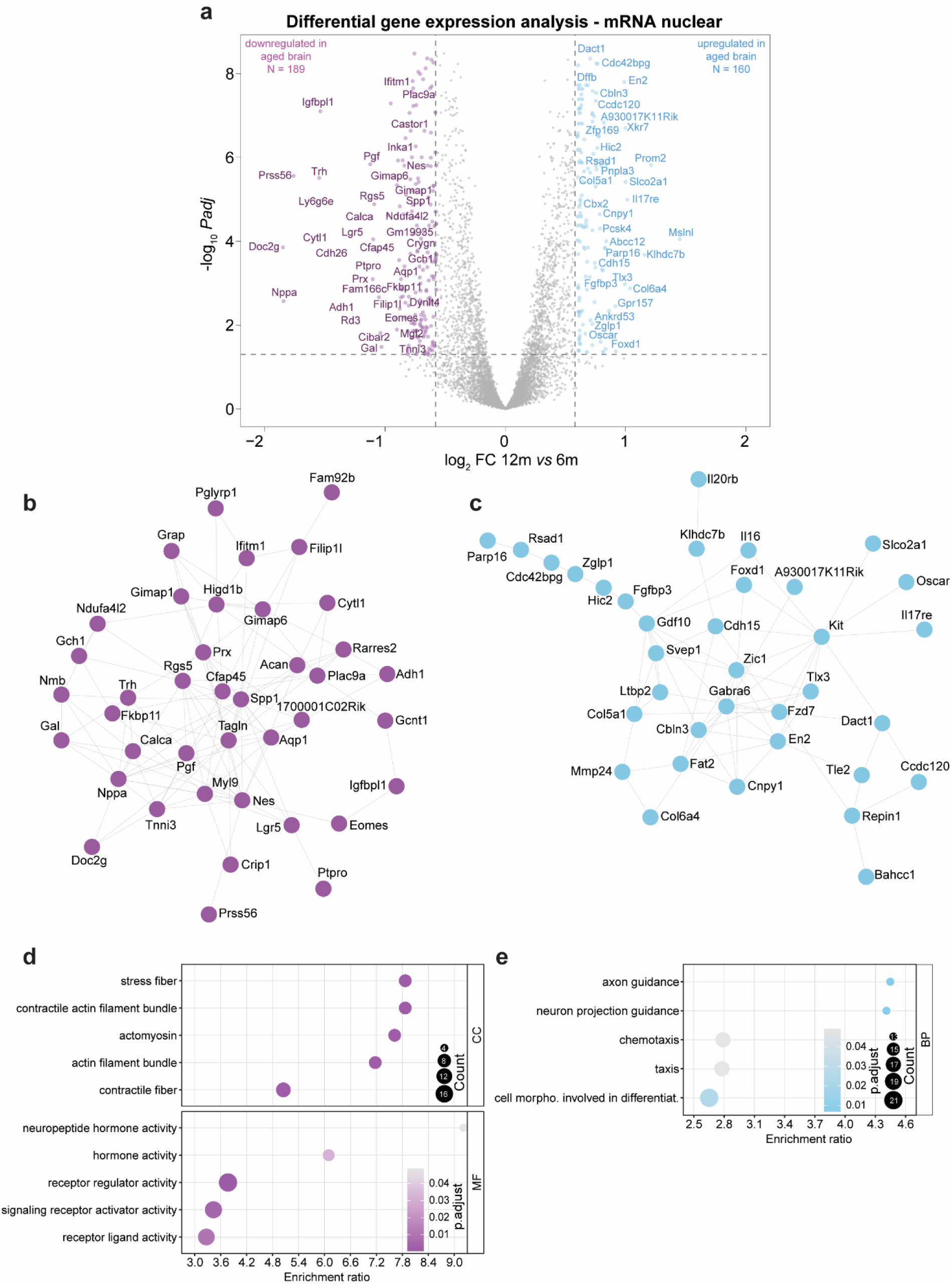
Gene expression changes in the aging mouse brain for 12m vs 6m in mRNA-nuclear fraction. (**a**) Volcano plot displaying the differentially expressed genes for 12m vs 6m. Inset: dot plot for biological replicate variability calculated using PCA, where sky blue represents increased expression with aging and medium orchid denotes decreased expression with aging. (**b, c**) STRING representation for the top 50 most (**b**) down- or (**c**)upregulated genes in 12m brain (respectively medium orchid and light blue). (**d, e**) GO-ORA of genes that are significantly down- or upregulated in 12m (*padj* ≤ 0.05, |log_2_FC| ≥ 0.58.

**Extended Data Figure 22:**
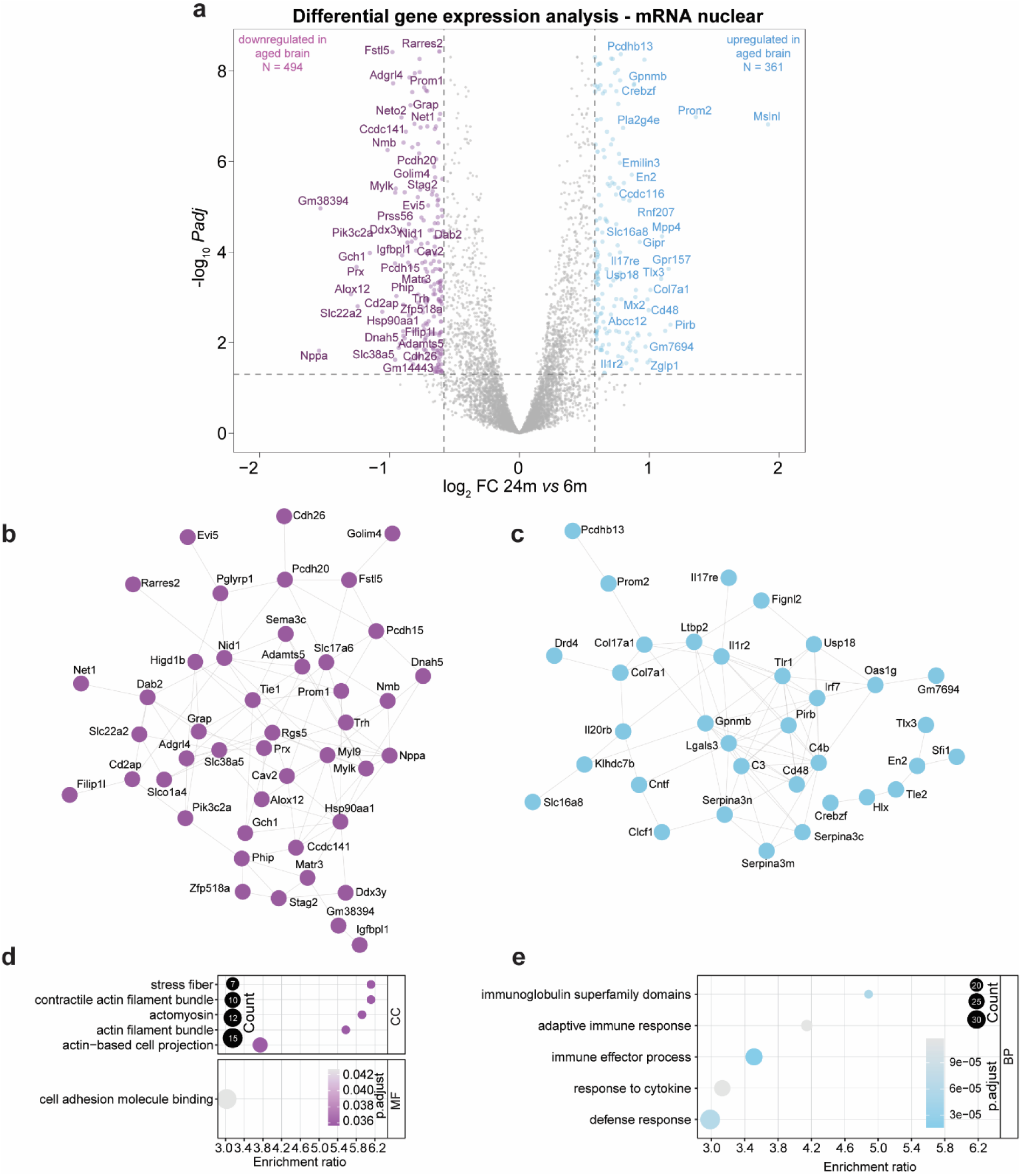
Gene expression changes in the aging mouse brain for 24m vs 6m in mRNA-nuclear fraction. (**a**) Volcano plot displaying the differentially expressed genes for 24m vs 6m. Inset: dot plot for biological replicate variability calculated using PCA, where sky blue represents increased expression with aging and medium orchid denotes decreased expression with aging. (**b, c**) STRING representation for the top 50 most (**b**) down- or (**c**) upregulated genes in 24m brain (respectively medium orchid and light blue). (**d, e**) GO-ORA of genes that are significantly down- or upregulated in 24m (*padj* ≤ 0.05, |log_2_FC| ≥ 0.58.

**Extended Data Figure 23:**
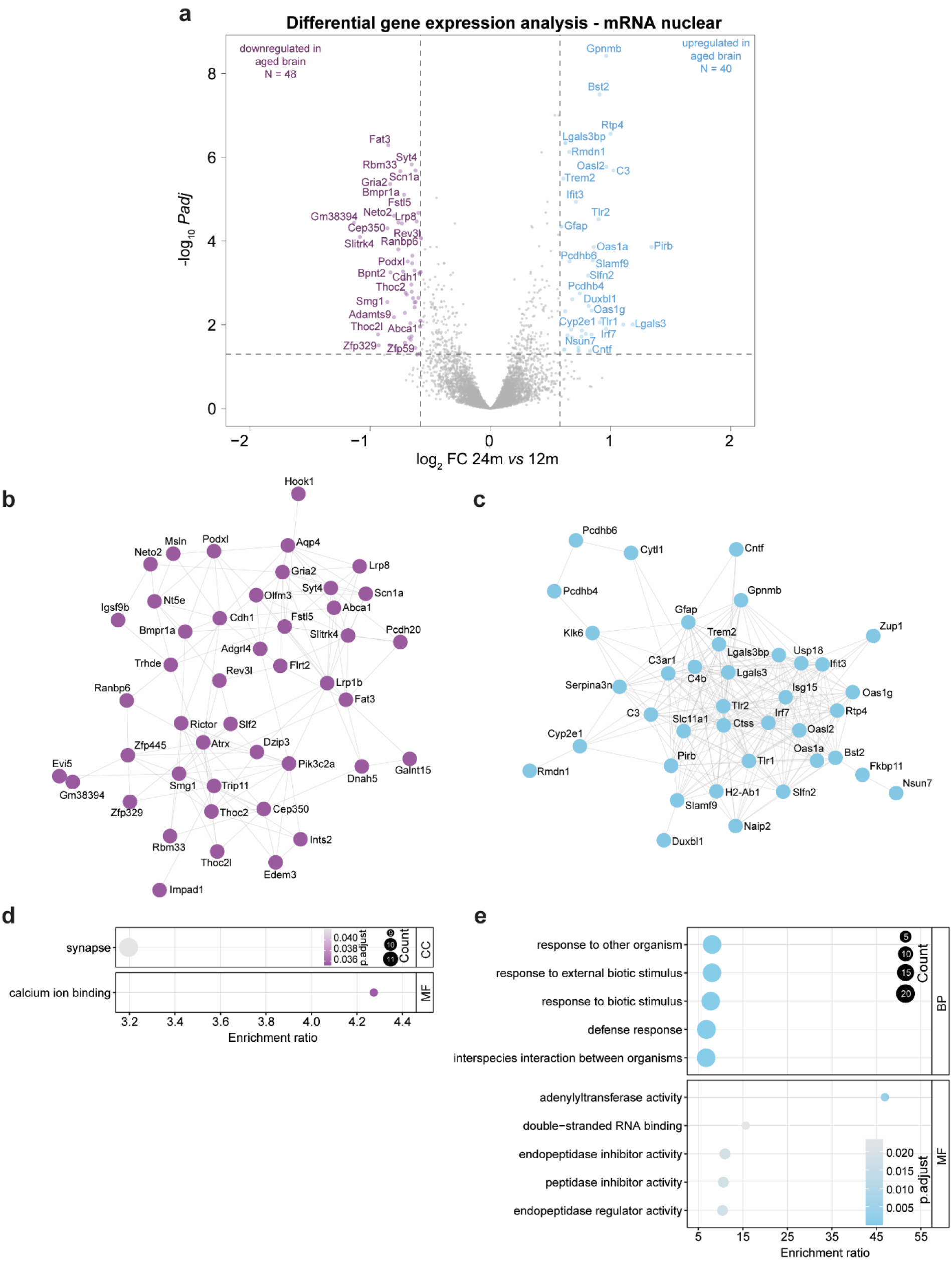
Gene expression changes in the aging mouse brain for 24m vs 12m in mRNA-nuclear fraction. (**a**) Volcano plot displaying the differentially expressed genes for 24m vs 6m. Inset: dot plot for biological replicate variability calculated using PCA, where sky blue represents increased expression with aging and medium orchid denotes decreased expression with aging. (**b, c**) STRING representation for the top 50 most (**b**) down- or (**c**)upregulated genes in 24m brain (respectively medium orchid and light blue). (**d, e**) GO-ORA of genes that are significantly down- or upregulated in 24m (*padj* ≤ 0.05, |log_2_FC| ≥ 0.58.

**Extended Data Figure 24:**
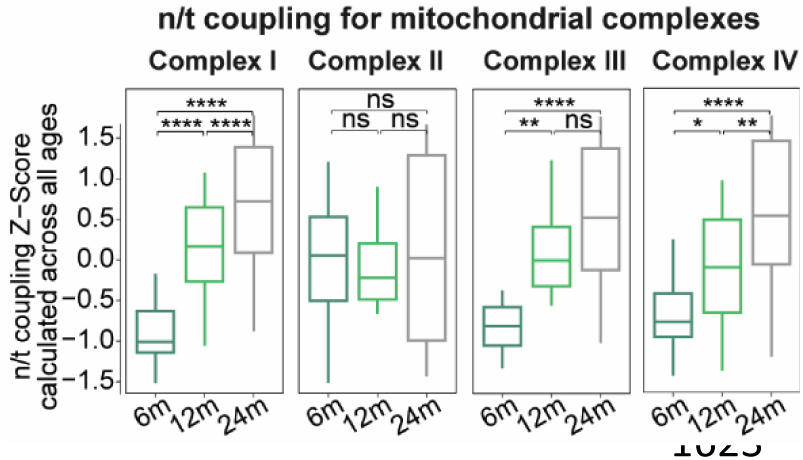
n/t coupling for mitochondrial complexes. Boxplots for n/t coupling for mitochondrial complexes. P-values indicate the results of paired t-test followed by Tukey posthoc test P-value * ≤ 0.05, ** ≤ 0.01, *** ≤ 0.001 and **** ≤ 0.0001.

**Extended Data Figure 25:**
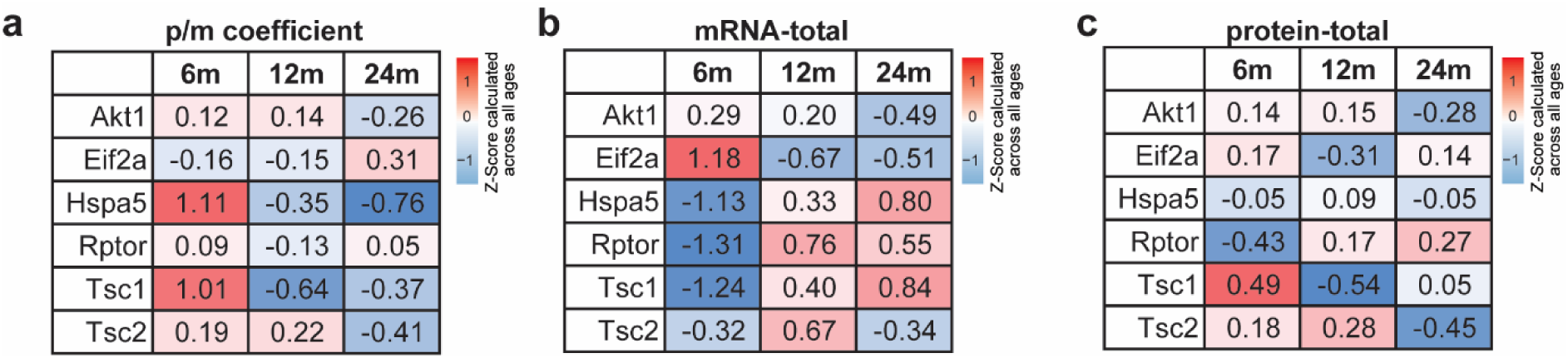
Heatmap of mTOR and ER stress genes. (a) p/m coefficient (b) mRNA-total and (c) protein-total. Colors based on the color names in R indicate Z-scores across all ages, where red signifies increased abundance relative to all ages, and skyblue signifies decreased abundance.

**Extended Data Figure 26:**
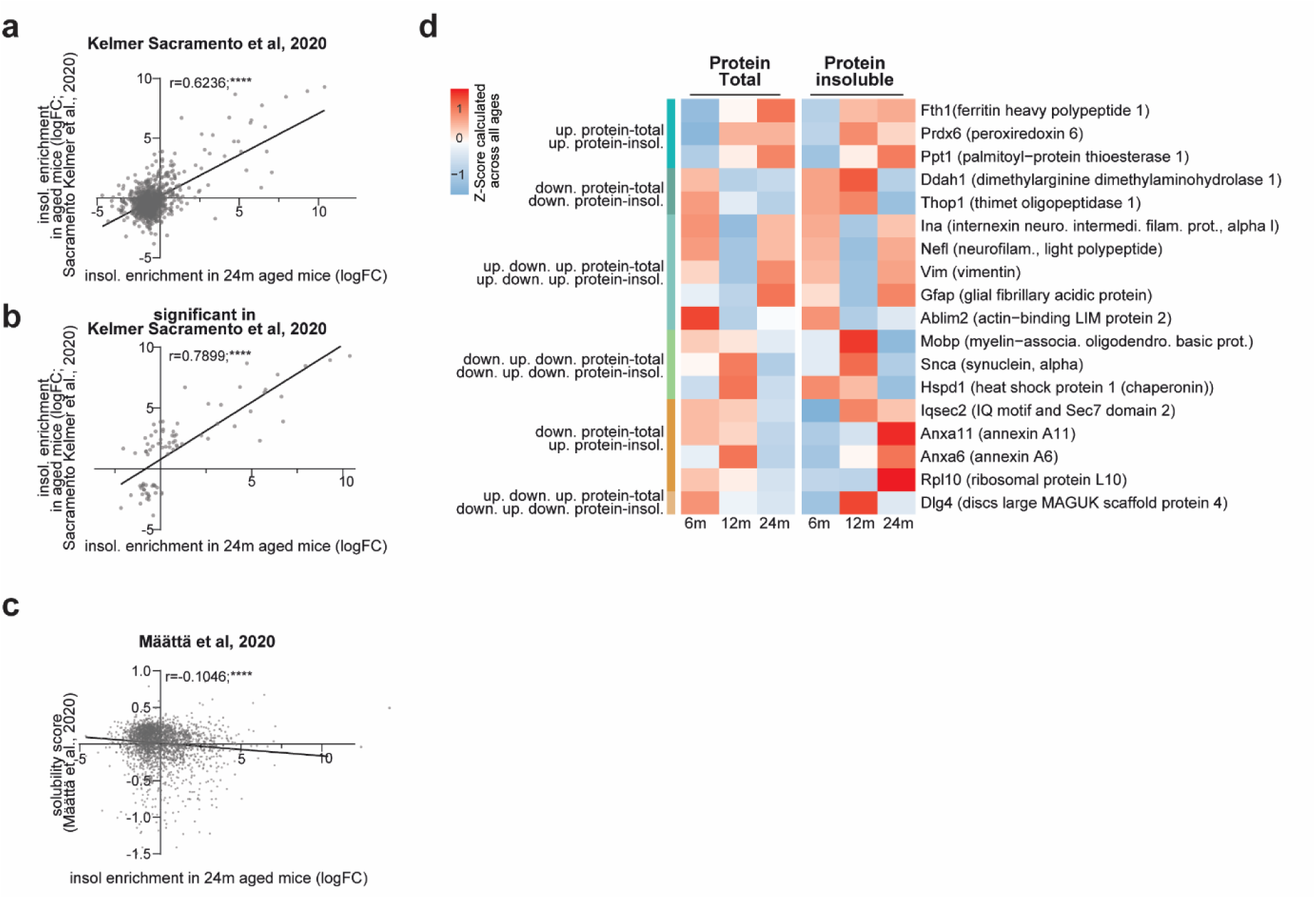
Protein level changes in the aging mouse brain in protein-insoluble fraction. (a, b) Comparison of insoluble-enriched proteins between our study and the study by Kelmer Sacramento et al. (a) shows the correlation of all common proteins and their insoluble enrichment at 24 months between the two studies, (b) shows the correlation between the significantly enriched proteins from the Kelmer Sacramento study and our data. (c) Correlation of our insoluble enrichment data to the protein solubility score introduced by Määttä at al., 2020. Note that increasing negative solubility score indicates higher aggregation propensity. Correlations are indicated as Pearson correlation coefficient. * p<0.5, ** p<0.01, **** p<0.0001. (d) Heatmap illustrating six distinct patterns in protein total and insoluble levels across ages. Color gradient for the Z-scores across all ages (red increased and blue decreased).

**Extended Data Figure 27:**
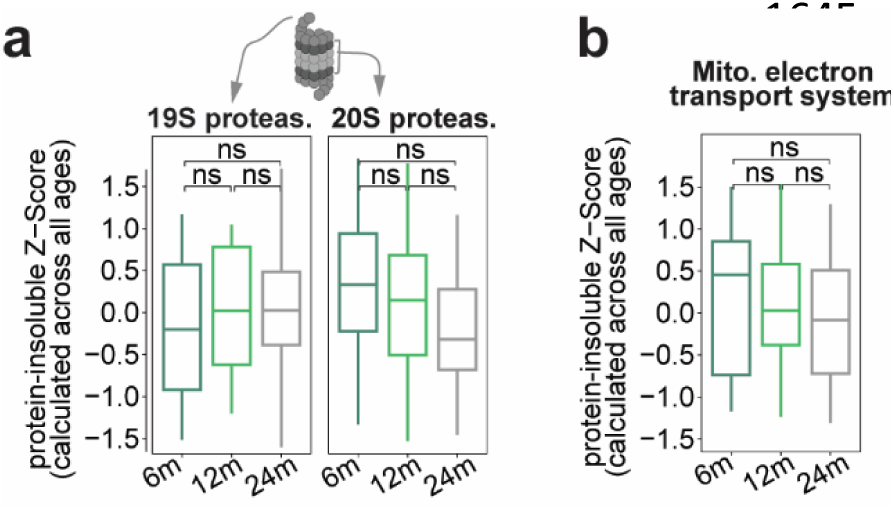
Changes in the protein - insoluble levels for the proteasome and mitochondrial proteins. (a, **b)** Boxplots for protein aggregate for (**a**) 19S and 20S proteasome complexes, and (**b**) proteins of the mitochondrial electron transport system (ETS). P-values indicate the results of paired t-test followed by Tukey posthoc test P-value * ≤ 0.05, ** ≤ 0.01, *** ≤ 0.001 and **** ≤ 0.0001.

**Extended Data Figure 28:**
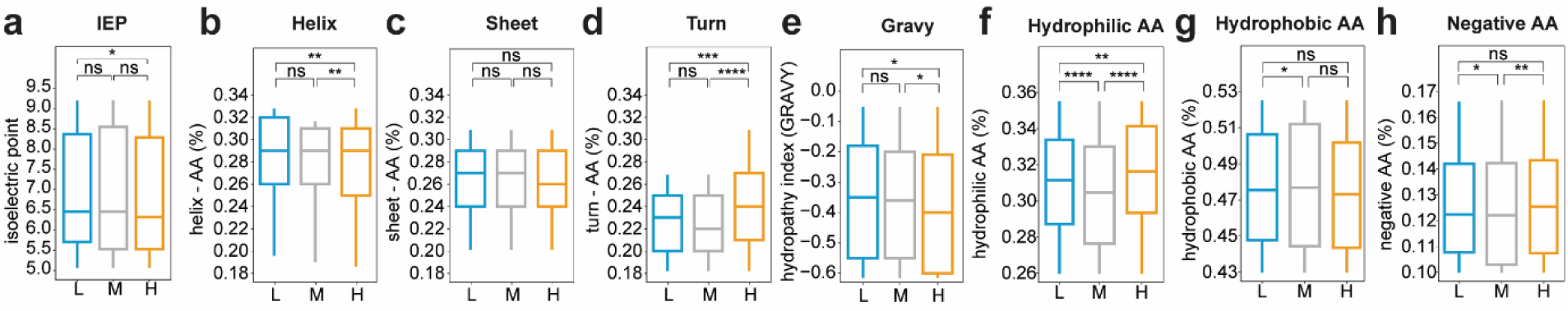
Biochemical property alterations in the aged brain. (a-h) Boxplot for proteins for high (H, increased), low (L, decreased) and middle (M, unchanged) in 24m compared to 12m. (a) Isoelectric point (IEP), (b) Amino acid percentage for Helix, (c) Amino acid percentage for Sheet, (d) Amino acid percentage for Turn, (e) Amino acid percentage for Hydropathy index (Gravy), (f) Amino acid percentage for hydrophilic amino acids (g) Amino acid percentage for hydrophobic amino acids, and (h) Amino acid percentage for negative amino acids. Colors based on the color names in R indicate, ‘goldenrod’ indicate high (H), ‘grey’ indicate middle (M) and ‘skyblue’ indicate low (L). P-values indicate the results of paired t-test followed by Tukey posthoc test P-value * ≤ 0.05, ** ≤ 0.01, *** ≤ 0.001 and **** ≤ 0.0001.

**Extended Data Figure 29:**
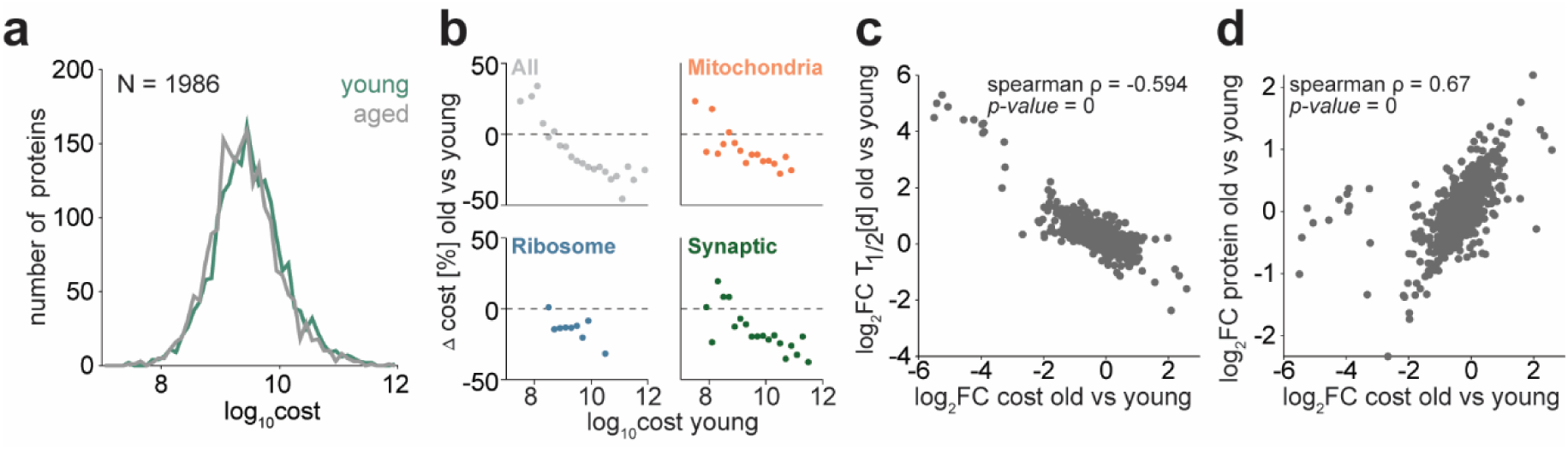
Energy imbalance in the aged brain. (**a**) The distribution of cost per protein species spans four orders of magnitude in young and older mice, hence the global cost strongly depends on the most expensive proteins. Bin size is 0.1 (in log_10_ scale). (**b**) Changes of cost per protein species are negatively correlated with its energetic burden in younger mice. Proteins are binned (bin size 0.2 in log_10_ scale) based on their associated cost in younger age and with changes shown in percentage. Grey – all proteins, orange - mitochondrial proteins, blue - ribosomal and green - synaptic proteins. (**c,d**) Spearman ρ correlation for the change in cost per protein with age is negatively associated with its (**c**) half-life and positively with its (**d**) log_2_FC.

**Extended Data Figure 30:**
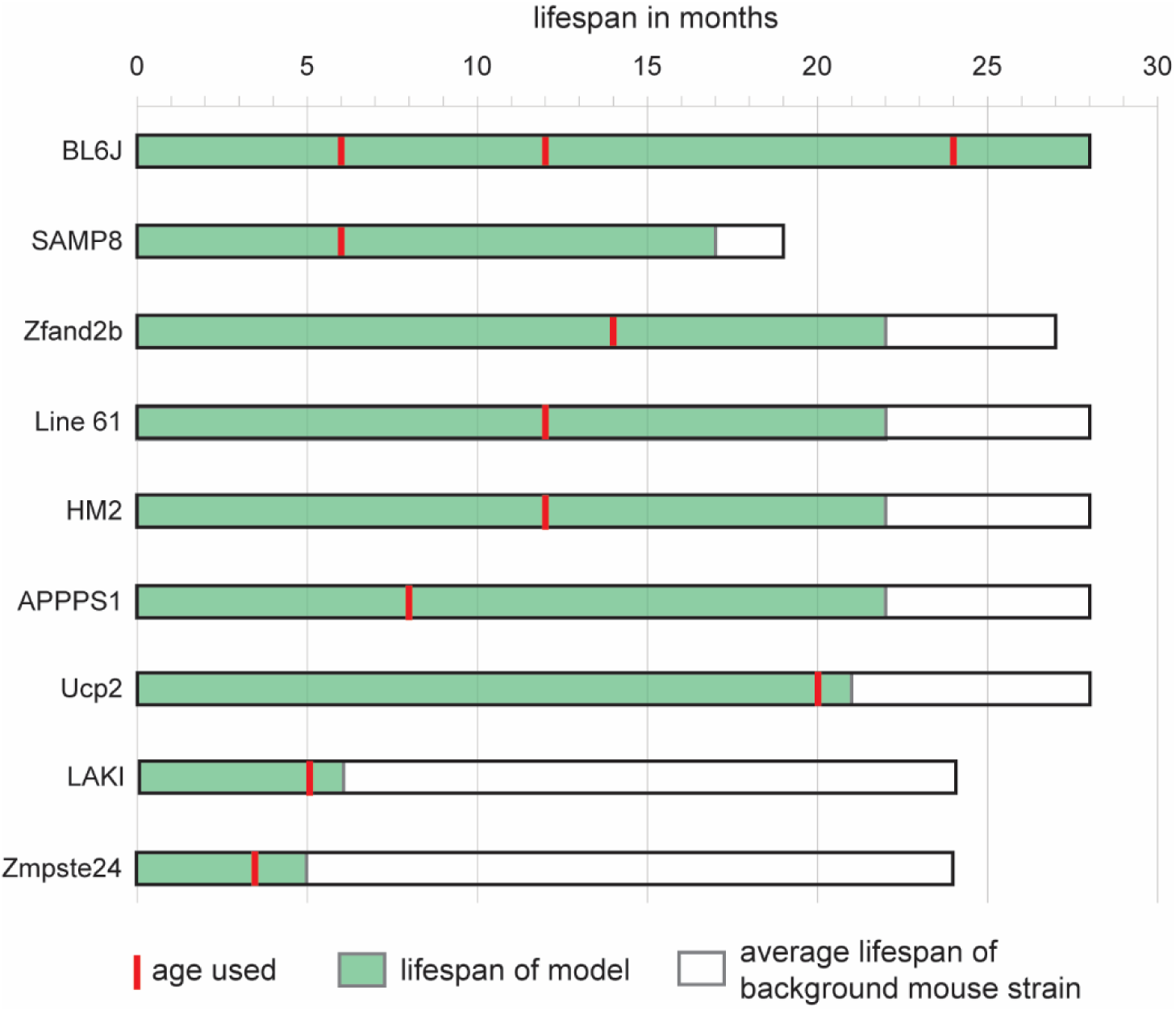
Overview of average lifespan and used ages of mouse model cohorts used in this study. General average lifespan of the specific mouse strain is indicated in white. If the model has extended or decreased lifespan, this is indicated in the overlaid green. The red line marks the age used here. Age in months. Data on lifespan was retrieved from the original publications for each model (see Methods).

**Extended Data Figure 31:**
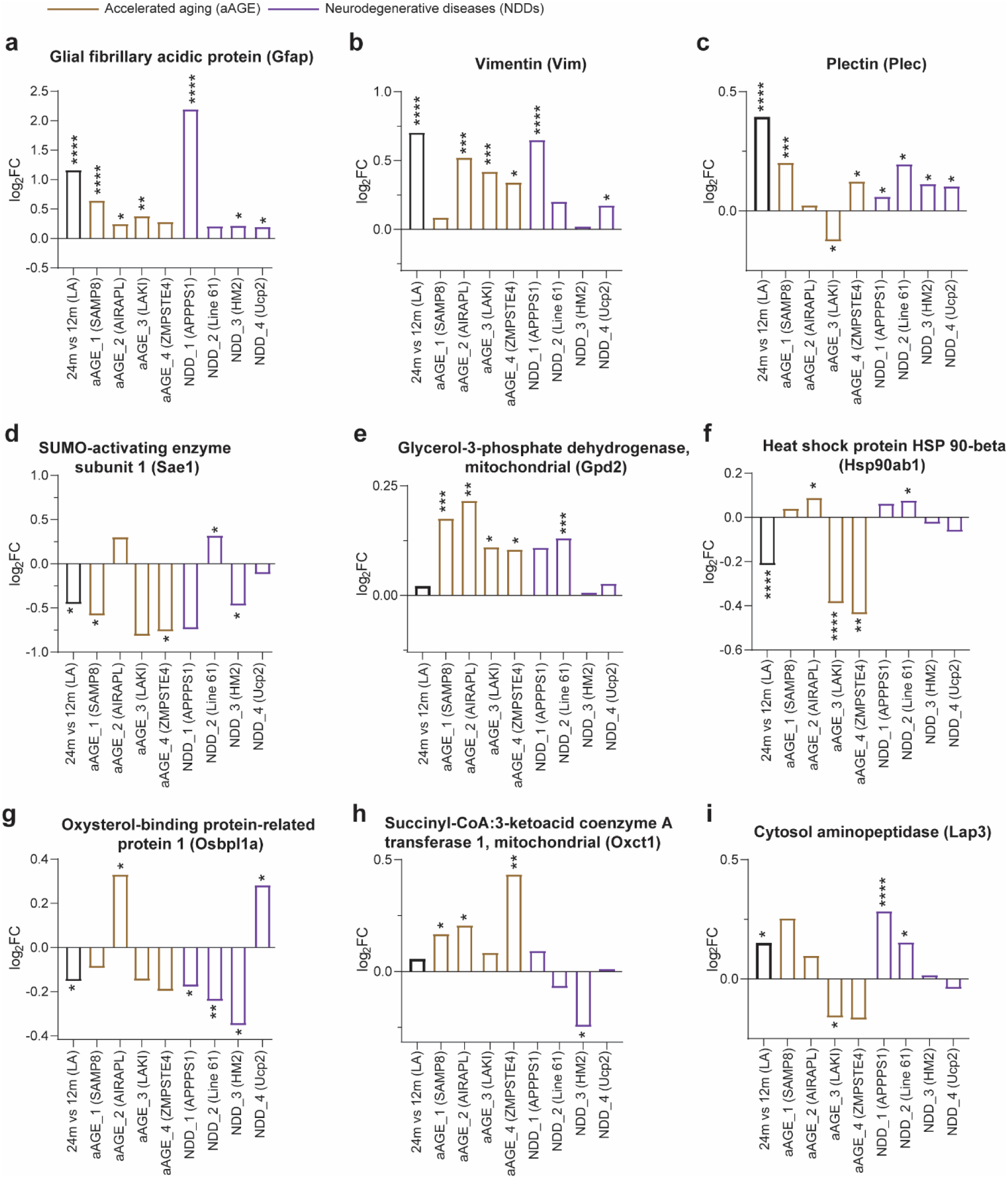
Proteins found significantly changed in at least 5 mouse models, including physiological aging 24 vs 12. Bar plot for proteins that are significantly changed in atleast 5 mouse models and late aging (**a**) Glial fibrillary acidic protein (Gfap), (**b**) Vimentin (Vim), (**c**) Plectin (Plec), (**d**) SUMO1 activating enzyme subunit 1 (Sae1), (**e**) Glycerol-3-phosphate dehydrogenase (Gpd2), (**f**) Heat shock protein 90 alpha family class B member 1 (Hsp90ab1), (**g**) Oxysterol binding protein like 1A (Osbpl1a), (**h**) Succinyl-CoA:3-ketoacid coenzyme A transferase 1, mitochondrial, (**i**) Leucine aminopeptidase 3 (Lap3). 24m vs. 12m (LA) – black, accelerated aging (aAGE) in ‘peru’, neurodegenerative diseases (NDD) in purple. Stars indicate DEqMS significance in the respective comparison of 24 vs 12 or of model vs littermate control: * p<0.05, ** p<0.01, *** p<0.001, **** p<0.0001.

**Extended Data Figure 32:**
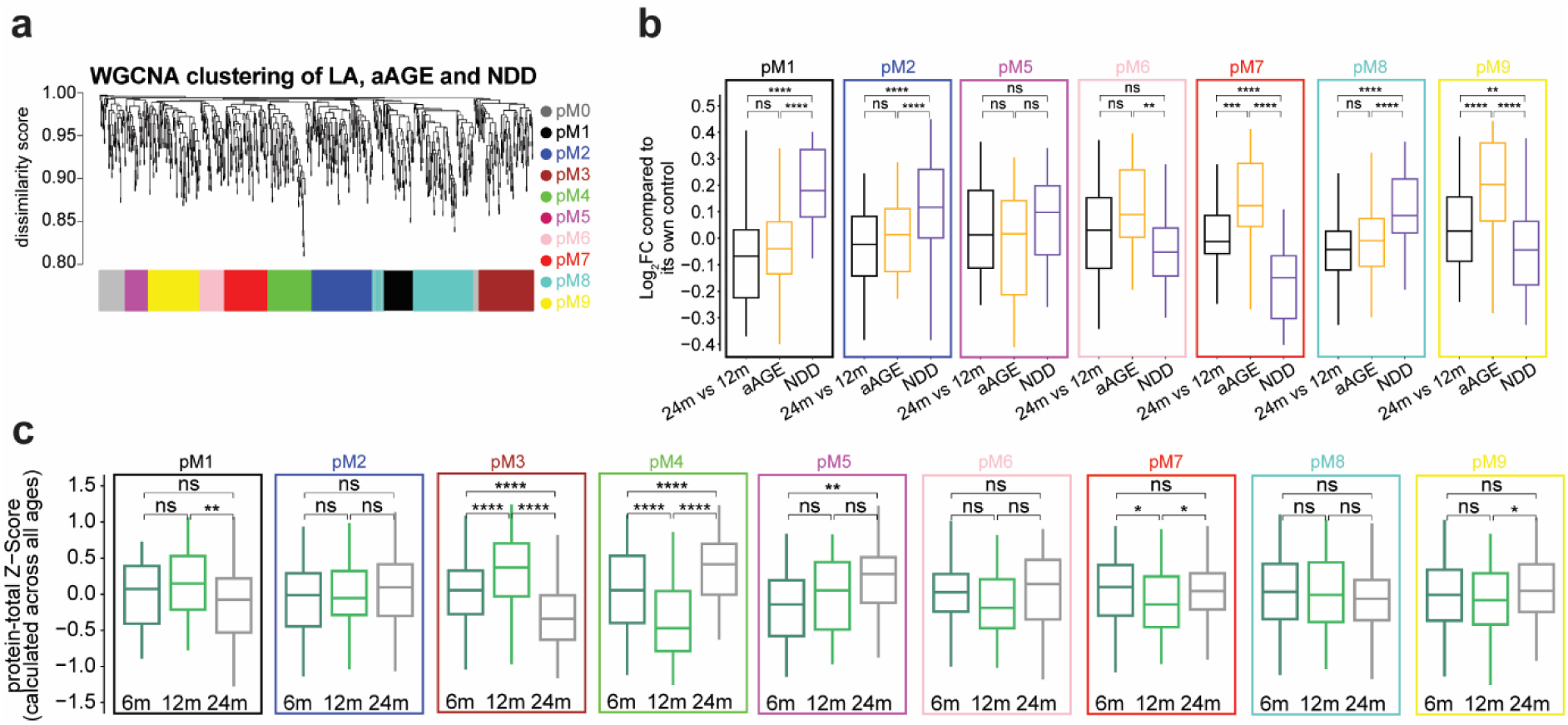
Module level alterations in late aging, aAGE and NDD brain proteomes. (**a**) WGNCA dendrogram with highlighted modules (lower colored bar). Proteins were clustered based on dissimilarity measures. The branches are modules of closely correlated proteomic groups that have a similar Log_2_FC in late aging and mouse models. Nine significant modules and M0 corresponding to ∼1000 genes were detected with WGCNA. M0 is a module with a less correlated protein group. (**b**) Boxplot for log2FC in 24m vs. 12m, average of aAGE and NDD models for pM1, pM2, pM5, pM6, pM7, pM8, and pM9 modules. (**c**) Boxplots of normalized protein-total levels in 6m, 12m and 24m mouse brain, grouped by their modules detected in the WGNCA method. P-values for paired t-test followed by Tukey post hoc test. P-value * ≤ 0.05, ** ≤ 0.01, *** ≤ 0.001 and **** ≤ 0.0001.

**Extended Data Figure 33:**
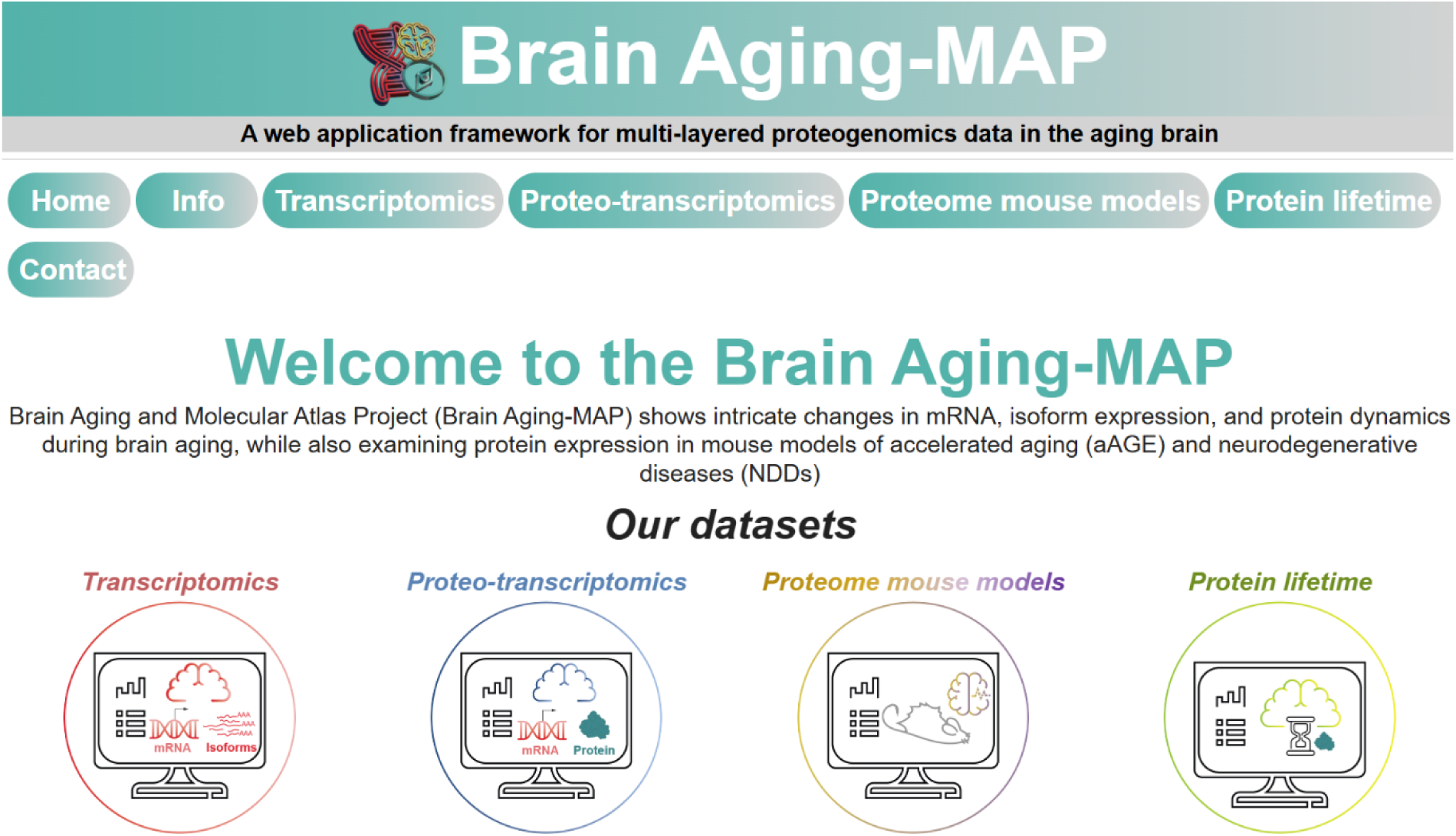
First page of the Brain Aging-MAP atlas. This page contains the navigation links to the individual datasets.

**Extended Data Figure 34:**
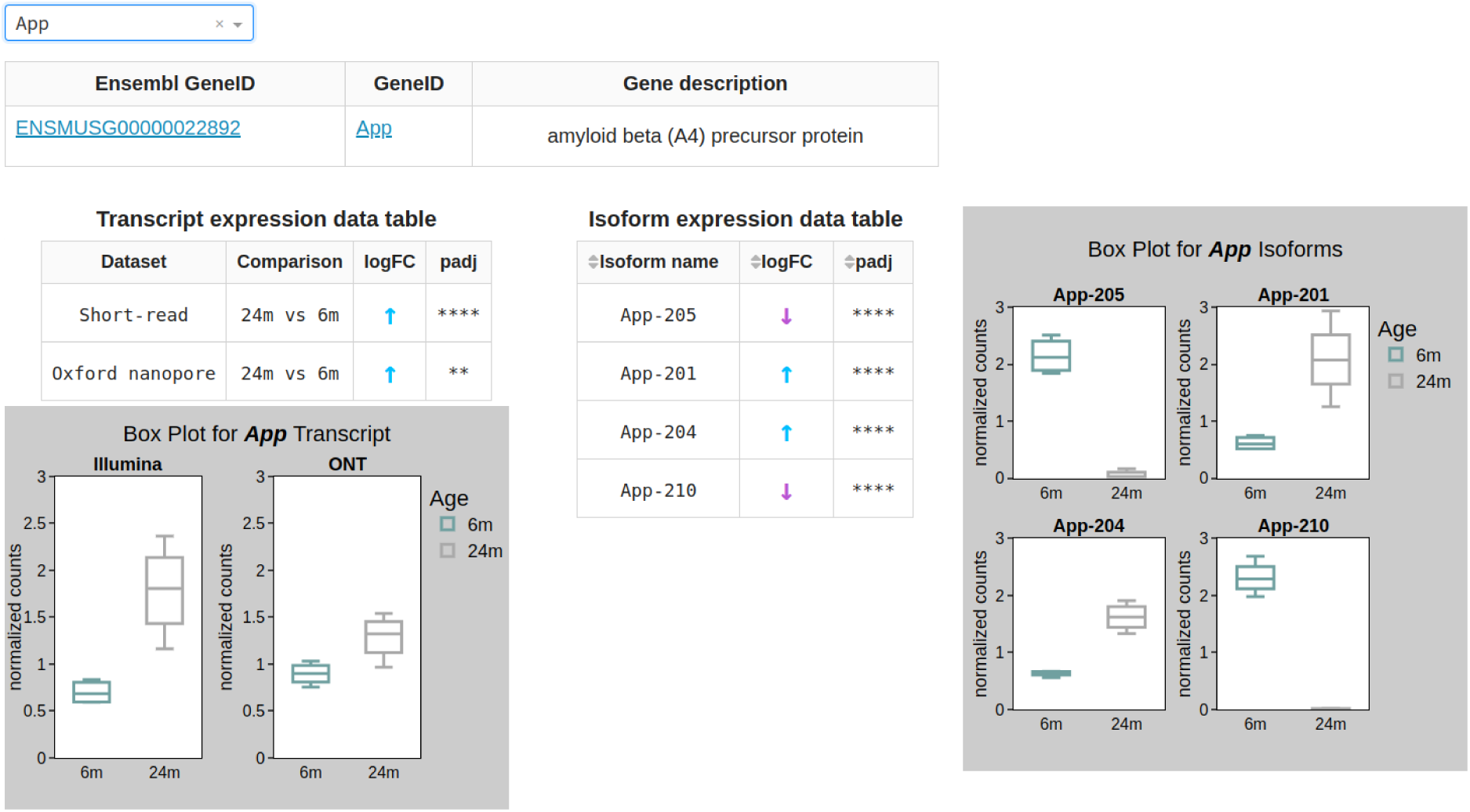
Exemplary transcriptomics page of the Brain Aging-MAP atlas. This page contains the gene and isoform expression information in physiological aging.

**Extended Data Figure 35:**
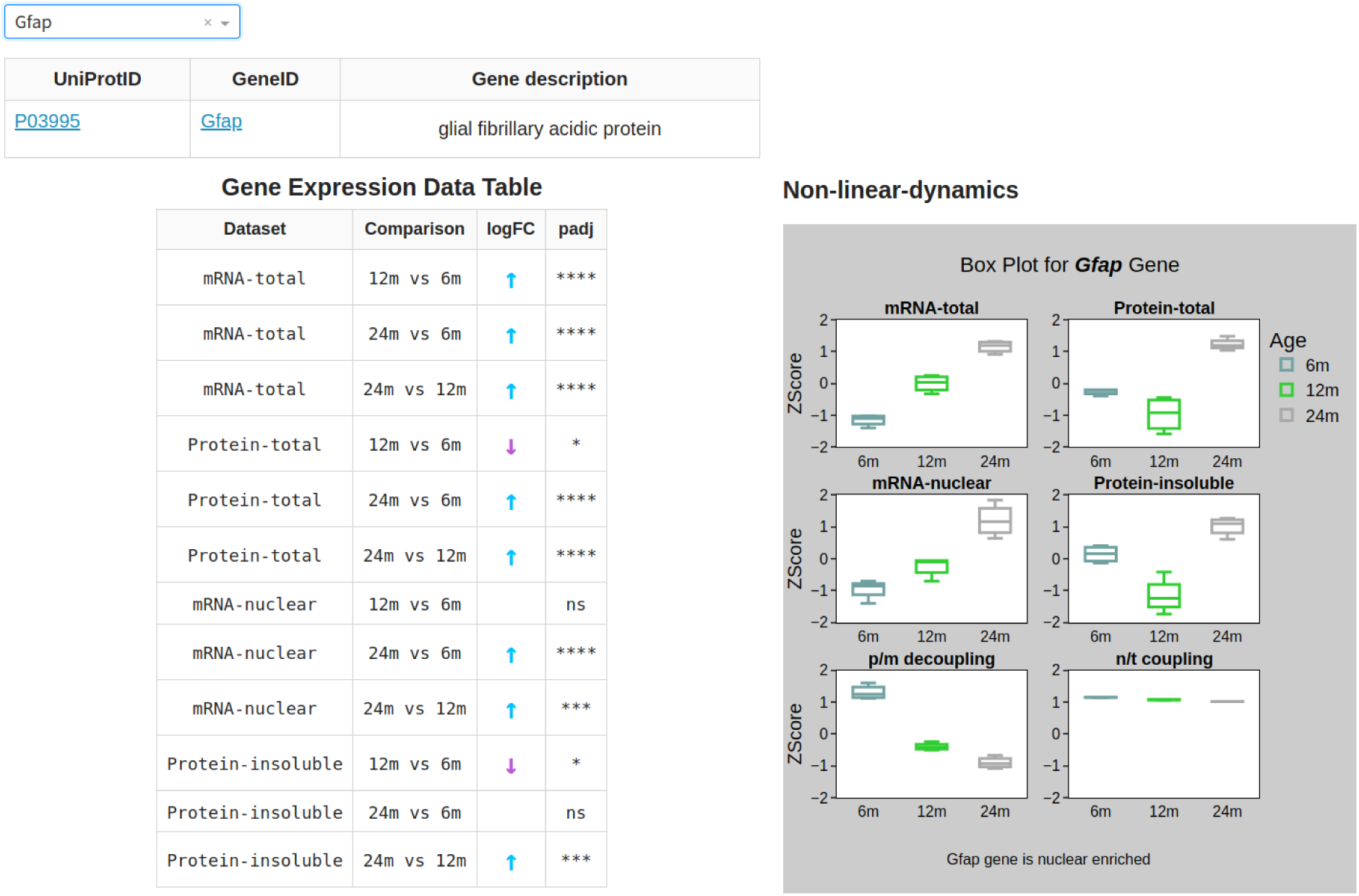
Exemplary proteo-transcriptomics page of the Brain Aging-MAP atlas. This page contains the multi-layered proteo-transcriptomics datasets, including total mRNA, nuclear mRNA, total protein, and insoluble protein datasets in physiological aging.

**Extended Data Figure 36:**
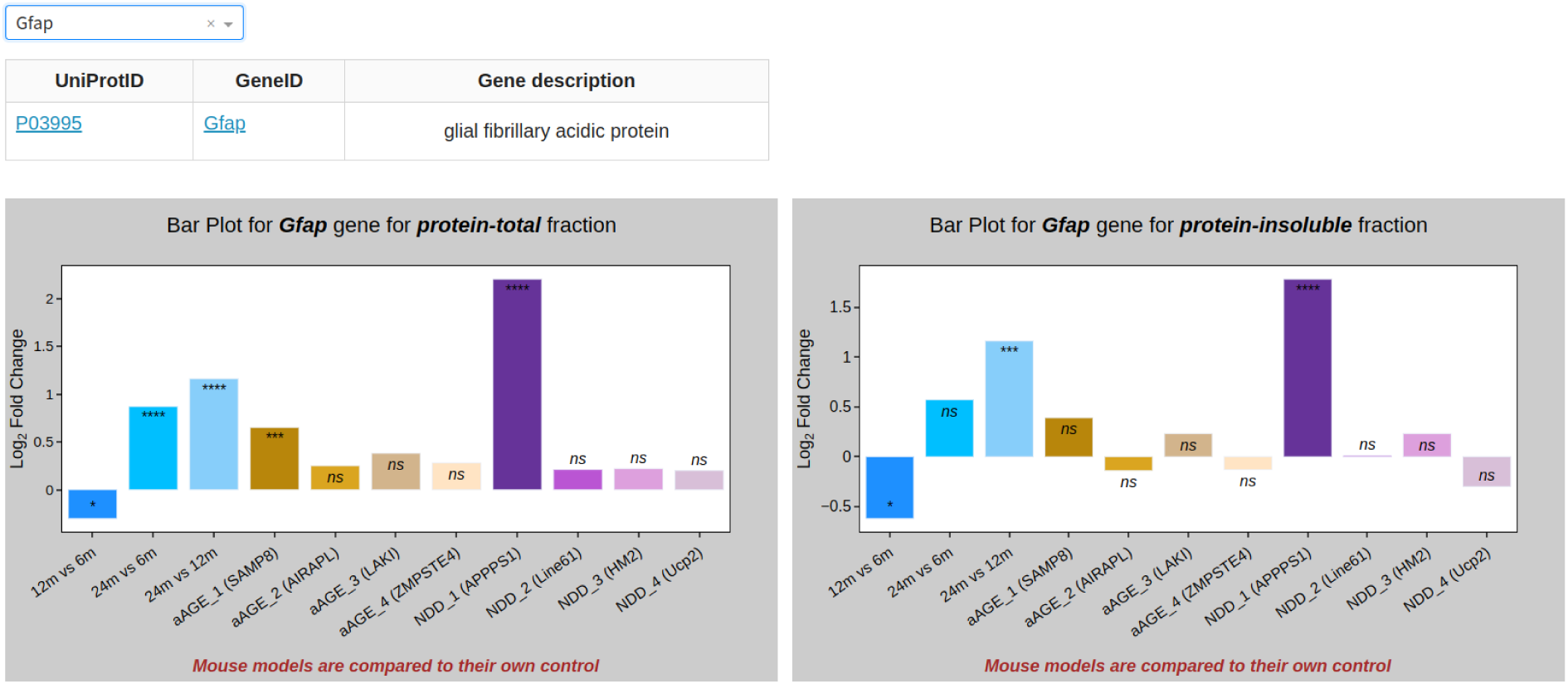
Exemplary proteome mouse models page of the Brain Aging-MAP atlas. This page contains the protein expression for the aAGE and NDD mouse models.

**Extended Data Figure 37:**
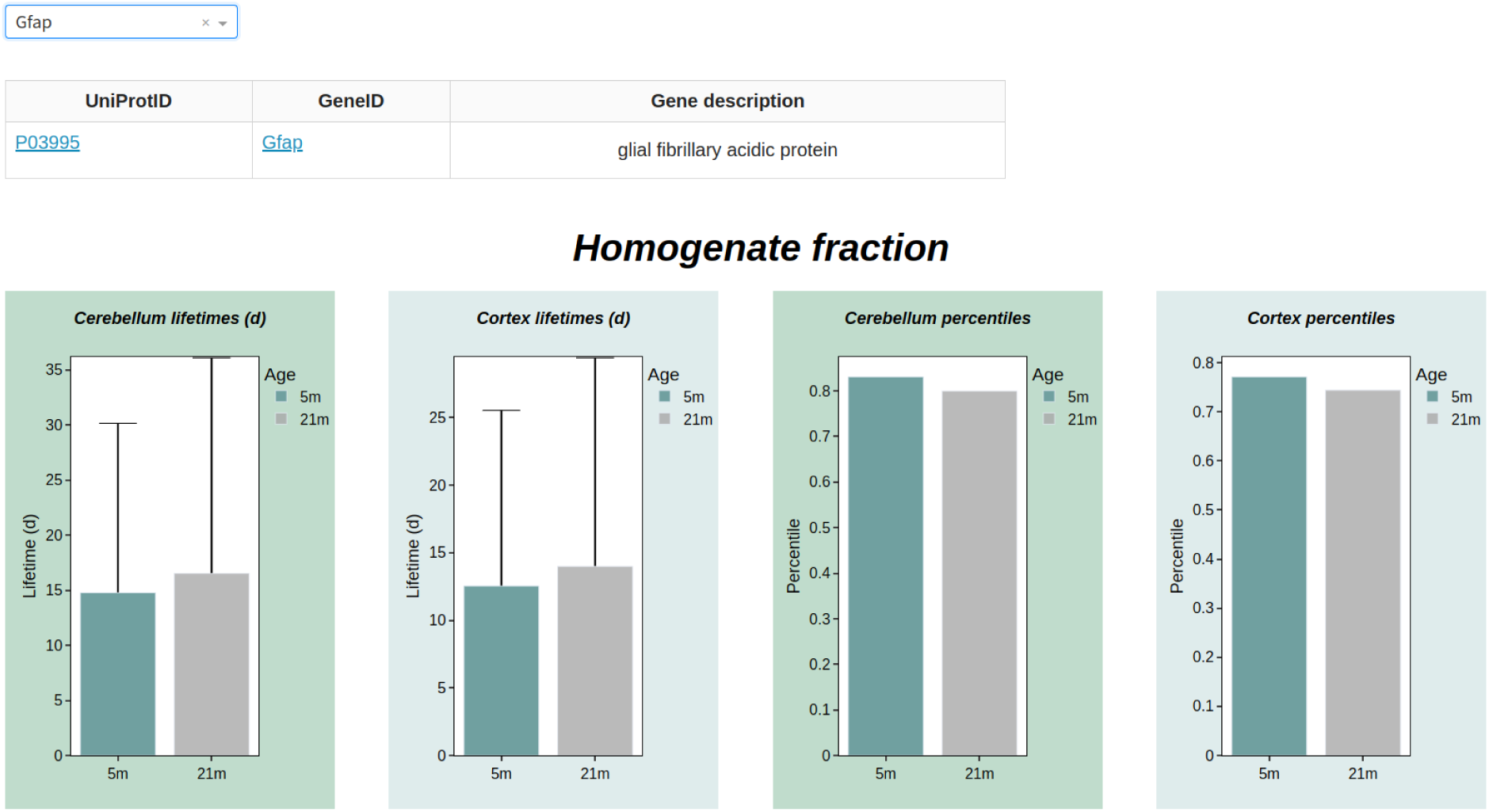
Exemplary protein lifetime page of the Brain Aging-MAP atlas. This page contains information about protein lifetime in physiological aging.

## Supplementary Text

### Supplementary Text 1: mRNA and protein level changes in the total fraction

Differential gene expression analysis in the aging brain was performed by comparing mRNA and protein levels across ages obtained through short-read sequencing and mass spectrometry methods.

We first evaluated genes that exhibited significant differences between 12m and 6m mice (early comparison or early aging) in our mRNA total fraction dataset, with a multiple comparison adjusted p-value (*padj*) ≤ 0.05 and an absolute log2 fold change (|log2FC|) ≥ 0.58 (**Supplementary Table 1**). When considering this subgroup of significantly differentially expressed mRNAs, we found 4184 upregulated transcripts corresponding to ∼55% and 3471 downregulated transcripts corresponding to ∼45%; **Extended Data Fig. 2a**, and **Supplementary Table 1**).

We then evaluated genes that exhibited significant differences between 24m and 6m mice (overall comparison or overall aging)^24^, selecting genes with a *padj* ≤ 0.05 and a |log2FC| ≥ 0.58 (**Supplementary Table 1**). When considering this subgroup of significantly differentially expressed mRNAs, we found 3989 upregulated transcripts corresponding to ∼54% and 3394 downregulated transcripts corresponding to ∼46% (**Extended Data Fig. 3a**, and **Supplementary Table 1**).

Overall, in 12m vs. 6m and 24m vs. 6m we found a slight over-representation of upregulated genes when compared to 6m. This small bias seems to be due to a bona-fide higher expression rather than a problem of normalization as observed in our previous studies^24^. When analyzing the highest and lowest 50 most significantly changed transcripts in 12m vs. 6m and 24m vs. 6m (Extended Data Fig. 2b, c and 3b, c, and Supplementary Table 1), we found some interesting hits that include Scgn and Rps29 downregulated in the aged brain. Scgn is a secreted calcium sensor that has a function related to neuroendocrine cells and has recently been shown to interfere with α-synuclein fibrillation ^112^. Rps29 is a ribosomal protein that is part of the small ribosomal subunit and is involved in translation to proteins. Among the 50 most upregulated mRNAs we observed a clear enrichment of transcripts encoding proteins with essential neuronal and synaptic functions^24^ (Extended Data Fig. 2c, 3c, and Supplementary Table 1). These include, for example, mRNAs related to the Gamma-Aminobutyric Acid (GABA) transporter (Slc2a13, and Slc7a14), potassium channels (Kcna2, and Kcnd3), and nuclear receptors (Nr2c2, Nr3c1). Other interesting hits include the ALF Transcription Elongation Factor 2 (Aff2) a RNA binding protein that could play a role in alternative splicing and the Neuralized E3 Ubiquitin Protein Ligase 1B (Neurl1b) that has a function related to ubiquitin-dependent endocytosis.

A more wide-ranging gene ontology ORA based on all changed transcripts confirmed a significant downregulation of mitochondrial and ribosomal mRNAs in the aged brain including several gene ontologies (GOs) associated to these processes (e.g., GO:0033108; GO:0007005, GO:0032981, GO:0042254, **Extended Data Fig. 2d, 3d**). The upregulated GOs point to synaptic transmission, organization, dendrite morphogenesis and development (GO:0050804, GO:0048813, GO:0016358, **Extended Data Fig. 2e, 3e**). These results indicate a possible engagement of neurons to produce more synaptic transcripts in the aged brain as observed in our previous study^24^.

Furthermore, we evaluated genes that exhibited significant differences between 24m and 12m mice (late comparisons or late aging), with a *padj* ≤ 0.05 and a |log2FC| ≥ 0.58 (Supplementary Table 1). Among these significantly differentially expressed mRNAs, we found 64 upregulated transcripts, corresponding to ∼39%, and 100 downregulated transcripts, corresponding to ∼61% (Extended Data Fig. 4a, and Supplementary Table 1). We observed a slight over-representation of downregulated genes. Notably, the 50 most downregulated transcripts were enriched for neuronal genes, including Shank1, Shank3, Gabrb2, Grm5, and Dlgap3. Additionally, we observed a downregulation of histone genes which play a central role in chromatin remodeling and transcription regulation^113^. These mRNAs include the histone H2, H3, and H4 families, for example, H2ac11, H3c13, H3c15, H4c1, H4c2, H4c3, H4c8, H4c9, H4c11, and H4c12. Interestingly, we also observed a downregulation with aging in two other comparisons: 12m vs 6m and 24m vs 6m (Extended Data Fig. 3b, Supplementary Table 1).

Among the 50 most upregulated mRNAs in the 24m vs 12m comparison, we observed genes related to the immune response, which is a hallmark of aging^39–41^. The ORA based on all significantly changed transcripts confirmed a significant downregulation of nucleosome and postsynaptic mRNAs in the aged brain, including several GOs associated with these processes (e.g., GO:0006334, GO:0097107; **Extended Data Fig. 4d**). The upregulated GOs point to the innate immune response and response to interferon beta and gamma (GO:0045087, GO:0034341, GO:0035456, **Extended Data Fig. 4e**). Our analysis revealed a significant upregulation of inflammatory and immune-related genes in the aging brain, consistent with previous reports of neuroinflammation as a hallmark of brain aging^39–41,72^. This immune activation may contribute to synaptic degeneration and functional impairment observed in both normal brain aging and neurodegenerative conditions. Notably, we found that these same pathways, when properly regulated, play crucial roles in neuroplasticity and neuronal stress resistance^39,114^. These findings suggest a delicate balance between beneficial and detrimental effects of immune activation in the aging brain, highlighting the complex interplay between inflammation and cognitive function.

We next analyzed proteins that exhibited significant differences between 12m and 6m mice focusing on proteins in our protein-total fraction dataset using with a *padj* ≤ 0.05 and a |log2FC| > 0 (Supplementary Table 1). In this comparison, we identified 12 upregulated proteins (∼44%) and 15 downregulated proteins (∼56%) (Extended Data Fig. 5a, and Supplementary Table 1). Similarly, we evaluated proteins with significant differences between 24m and 6m mice, with the same criteria of *padj* ≤ 0.05 and a |log2FC| > 0 (Supplementary Table 1). Among this subset of significantly differentially expressed proteins, we observed a slight over-representation of upregulated proteins, with 24 upregulated (∼77%) and 7 downregulated (∼23%) (Extended Data Fig. 6a, and Supplementary Table 1). Additionally, when comparing 24m and 12m mice, we found 20 upregulated proteins (∼54%) and 17 downregulated proteins (∼46%) (Extended Data Fig. 7a, and Supplementary Table 1).

We then evaluated genes that exhibited significant changes in mRNA and protein levels and observed an upregulation of astrocyte-specific markers. Glial fibrillary acidic protein (Gfap) often used as an astrocyte marker, showed the most pronounced upregulation, with increased expression at both mRNA and protein levels in 24m mice compared to 6m and 12m mice (**Extended Data Fig. 8, 9, 10 and 11b, c**). Similarly, another astrocyte-specific gene, vimentin (Vim), exhibited elevated expression at 24m compared to 12m (**Extended Data Fig. 10, 11c**) at both transcriptional and translational levels. These results are similar to previous studies^27,39–41,115,116^ which suggest glial activation in the aging brain.

Among the proteins significantly downregulated at 24m compared to 6m, we identified TGF-Beta Activated Kinase 1 Binding Protein 3 (Tab3), associated with neuronal apoptosis^117^, and Cullin1 (Cul1), involved in proteolysis^118^. These findings suggest alterations in homeostatic processes in the aging brain. We also observed an upregulation of Complement C1q B Chain (C1qb), which is involved in immune function at 24m compared to 6m.

Mitochondrial Contact Site and Cristae Organizing System Subunit 10 (Micos10), involved in inner mitochondrial membrane organization was upregulated at 24m compared to 6m and at 12m compared to 6m. Interestingly, Regulatory Associated Protein of mTOR Complex 1 (Rptor), a component of the mTORC1 complex that regulates cell growth and metabolism was upregulated at 24m compared to 6m at both mRNA and protein levels (Extended Data Fig 9, 11b). This suggests an impaired autophagy and altered protein synthesis potentially due to a hyperactivation of the mTOR pathway^119^ (Extended Data Fig 9, 11b). This upregulation of mTOR pathway is also associated with neurodegenerative diseases which might suggest an age-related cognitive decline.

ORA of the top 150 proteins across different age comparisons with a non-adjusted p-value < 0.05 revealed distinct of enrichment GO terms (Supplementary Table 1). In the early comparison, ‘oxoacid metabolic process’ (GO:0043436) and ‘carboxylic acid metabolic process’ (GO:0019752) were significantly enriched among upregulated proteins (Extended Data Fig 5e), while ‘intermediate filament-based process’ (GO:0045103) was enriched among downregulated proteins (Extended Data Fig 5d). The late comparison showed a shift, with ‘intermediate filament-based process’ (GO:0045103, Extended Data Fig 7e) becoming enriched among upregulated proteins, and several metabolic processes, including ‘monocarboxylic acid metabolic process’ (GO:0032787), ‘ATP metabolic process’ (GO:0046034), and ‘oxoacid metabolic process’ (GO:0043436), enriched among downregulated proteins (Extended Data Fig 7d). The overall comparison showed an activation of GO terms related to immune response (GO:0006956, GO:0002455, Extended Data Fig 6d).

Interestingly, several intermediate filaments, including neurofilament polypeptides (Nefl, Nefm, Nefh), vimentin (Vim), plectin (Plec), and α-internexin (Ina), exhibited decreased levels at 12m and were significant in both early and late comparisons (Extended Data Fig 10, 11a, c), though not between 24m vs. 6m. This suggests that aging is a non-linear process and highlights the importance of looking more closely at the genes that show a similar trend and the other non-linear changes in the aging process. When analyzing the downregulated proteins at 24m compared to 12m, we found proteins involved in protein folding and stress response such as heat shock proteins Hsp90ab1, Hspa4, and Hsph1, and co-chaperones Ahsa1, Ppid, and Canx (Extended Data Fig. 1c).

Furthermore, having perfectly-matched datasets allowed us to perform novel isoform discovery. After combining information from transcriptomics and proteomics, we were able to identify novel isoforms expressed at different ages. We identified several proteoforms (Supplementary Table 1) and a notable finding are the two non-canonical sequences of Syt7. Interestingly, the Syt7_1 proteoform, which had 84.1% similarity to the Syt_7 canonical sequence, and this proteoform were significantly downregulated in late aging. From these results, we could speculate that in general, Syt7 protein is downregulated with aging. In the future, having longer reads at 24m and 12m for the transcriptomics datasets could potentially help in the identification of novel isoforms. As these were quantified using the novel isoforms predicted from the 24m and 6m datasets^24^.

### Supplementary Text 2: mRNA level changes in the nuclear fraction

We then evaluated genes that exhibited significant differences between 12m and 6m mice, in our mRNA-nuclear fraction dataset with a *padj* ≤ 0.05 and a |log2FC| ≥ 0.58 (Supplementary Table 5). This comparison yielded 160 upregulated (∼47%) and 189 downregulated (∼53%) transcripts (Extended Data Fig. 21a and Supplementary Table 5). Similar criteria were applied to compare 24m and 6m mice, resulting in 174 upregulated (∼39%) and 204 downregulated (∼61%) transcripts (Extended Data Fig. 22a and Supplementary Table 5).

A slight over-representation of downregulated genes was observed at 24m and 12m when compared to 6m. Analysis of the 50 most significantly downregulated transcripts in both comparisons (**Extended Data Fig. 21b, 22b and Supplementary Table 5**) revealed notable genes including myosin light chain kinase (Mylk) and myosin light chain 9 (Myl9), which are localized in actin filaments (GO:0032432, GO:0097517; **Extended Data Fig. 21d, 22d and Supplementary Table 5**). Among upregulated genes, immune response-related transcripts were prominent in the 24m vs. 6m comparison (**Extended Data Fig. 22c, e and Supplementary Table 5**), while neuronal mRNAs were more affected in the 12m vs. 6m comparison (GO:0007411, GO:0097485; **Extended Data Fig. 21c, e and Supplementary Table 5**).

Comparison between 24m and 12m mice revealed 48 downregulated (∼55%) and 40 upregulated (∼45%) transcripts (**Extended Data Fig. 23a and Supplementary Table 5**). Analysis of the most significantly altered transcripts revealed upregulation of genes involved in immune and defense response (GO:0006955, **Extended Data Fig. 23c, e, Supplementary Table 5**) and endopeptidase regulator activity (GO:0004866**; Extended Data Fig. 23c, e; and Supplementary Table 5**) in the aged brain. These included complement C3 (C3), serpin family A member 3 (Serpina3n), bone marrow stromal cell antigen 2 (Bst2), and NLR family apoptosis inhibitory protein (Naip2). Increased levels of Serpina3n and C3 have been associated with age-related changes in brain function and the onset of brain amyloidosis in Alzheimer’s disease^120,121^ (AD). Notably, Ran-Binding Protein 6 (RanBP6), a nuclear transport receptor, was downregulated in both 24m vs. 6m and 12m vs. 6m comparisons, aligning with known nuclear transport dysfunction in the aged brain^122^. Additionally, we observed significant downregulation of synapse and calcium ion binding mRNAs in the aged brain for 24m vs 12m comparison (GO:0045202; **Extended Data Fig. 23b, d; and Supplementary Table 5**).

Overall, our analysis in the physiologically aging brain revealed a downregulation of genes involved in synapse function and an upregulation of immune response in the late comparisons, consistent with the previous studies^39–41^.

### Supplementary Text 3: Age-related changes in protein solubility

An upregulation at 24m for some proteins involved in cell signaling, trafficking, protein translation, and synaptic maintenance. For instance, Iqsec2, Anxa11, Anxa6 and Rpl10. Notably, we found that Iqsec2 has an IUPRED score of 0.7, indicating that it has a high amount of intrinsically disordered regions. Interestingly, 60S ribosomal proteins showed an increased insolubility at 24m (**Fig. 4e**) suggesting a reduction in functional ribosomes in the cytoplasm. However, we did not observe significant differences in the insolubility for the cytoskeletal, mitochondrial and proteasomal proteins (**Fig**. **4d****; Extended Data Fig. 27**).

### Supplementary Text 4: Age-related shifts in metabolic costs

We calculated the metabolic cost per proteo-transcriptome species based on protein turnover parameters previously measured in young and aged brains in a previous work^5^ and combined mRNA and protein level measurements obtained in this study. We found that the distribution of cost per proteo-transcriptome species is heavy-tailed, spans roughly four orders of magnitude and is superimposable at different ages (**Extended Data Fig. 29a**). When comparing the cost change in aged brains vs young ones, we found that specifically high-cost proteins experience a reduction in their energetic burden with age (**Extended Data Fig. 29a**), leading to a proteome-wide cost reduction of 26.24%. In contrast, proteins with low cost exhibited elevated energetic demands with age. Neither mitochondrial, ribosomal, nor synaptic proteins show strongly deviating trends from the overall background (**Extended Data Fig. 29b)** Overall, the changes in costs are driven by both variable parameters, protein half-life and protein levels (**Extended Data Fig. 29c, d**). If reduction in energy availability requires maintaining abundant but expensive proteins under increased translational pressure, this would lead to an increased translational efficiency over proteins with a low energetic burden, leading to a relative increase in their levels despite high biosynthetic cost. This rebalancing of proteome composition in aging is possibly an adaptive reaction to maintain cellular homeostasis during the condition of limited metabolic resources.

### Supplementary Text 5: Protein level changes in the mouse models

To obtain the proteins that are differentially expressed, we performed differential gene expression analysis and selected genes that showed significant changes (*padj* ≤ 0.05) in at least one mouse model or in physiological aging, in either protein total or insoluble fraction (**Supplementary Table 8**). This resulted in ∼1000 genes for our further analysis that includes multiomics gene set enrichment analysis (GSEA) and WGCNA analysis. Among the genes that were differentially expressed we observed significant changes for the genes involved in gliosis, inflammation, maintenance of neuronal structure, protein modification pathways, protein homeostasis, and lipid metabolism. For example, Gliosis and inflammation markers Gfap and Vim (**Extended Data Fig. 31a, b**), were upregulated in late aging (24m vs 12m), several aAGE models, and NDD models. Plectin (Plec, **Extended Data Fig. 31c**), involved in maintenance of cytoskeletal structure, was upregulated in late aging, all NDD models, and two aAGE models, but downregulated in aAGE_3. SUMO1 Activating Enzyme Subunit 1 (Sae1, **Extended Data Fig. 31d**) was downregulated in late aging and several models but upregulated in NDD_2. Glycerol-3-phosphate dehydrogenase (Gpd2, **Extended Data Fig. 31e**) was upregulated in late aging and all models, particularly significant in aAGE models and NDD_2. Chaperone regulation was impacted, as evidenced by the strong downregulation of Hsp90ab1 (**Extended Data Fig. 31f**) in late aging and nuclear envelope-related aAGE models. Oxysterol Binding Protein Like 1A (Osbpl1a, **Extended Data Fig. 31g**), gene involved in lipid metabolism was downregulated for late aging and all mouse models except aAGE_2 and NDD_4, which showed an upregulation. Succinyl-CoA:3-ketoacid coenzyme A transferase 1, mitochondrial (Oxct1, **Extended Data Fig. 31h**), involved in ketone body metabolism was significantly upregulated in three aAGE models but downregulated in NDD_3. Leucine Aminopeptidase 3 (Lap3, **Extended Data Fig. 31i**) was significantly upregulated in late aging and two NDD models but downregulated in nuclear envelope-related aAGE models (aAGE_3 and aAGE_4).

